# The P3 *O*-*Tert*-Butyl-Threonine is Key to High Cellular and Antiviral Potency for Aldehyde-Based SARS-CoV-2 Main Protease Inhibitors

**DOI:** 10.1101/2021.12.18.473326

**Authors:** Yuying Ma, Kai S. Yang, Zhi Zachary Geng, Yugendar R. Alugubelli, Namir Shaabani, Erol C. Vatansever, Xinyu R. Ma, Chia-Chuan Cho, Kaustav Khatua, Lauren Blankenship, Ge Yu, Banumathi Sankaran, Pingwei Li, Robert Allen, Henry Ji, Shiqing Xu, Wenshe Ray Liu

## Abstract

As an essential enzyme to SARS-CoV-2, main protease (M^Pro^) is a viable target to develop antivirals for the treatment of COVID-19. By varying chemical compositions at both P2 and P3 sites and the *N*-terminal protection group, we synthesized a series of M^Pro^ inhibitors that contain *β*-(S-2-oxopyrrolidin-3-yl)-alaninal at the P1 site. These inhibitors have a large variation of determined IC_50_ values that range from 4.8 to 650 nM. The determined IC_50_ values reveal that relatively small side chains at both P2 and P3 sites are favorable for achieving high *in vitro* M^Pro^ inhibition potency, the P3 site is tolerable toward unnatural amino acids with two alkyl substituents on the *α*-carbon, and the inhibition potency is sensitive toward the *N*-terminal protection group. X-ray crystal structures of M^Pro^ bound with 16 inhibitors were determined. All structures show similar binding patterns of inhibitors at the M^Pro^ active site. A covalent interaction between the active site cysteine and a bound inhibitor was observed in all structures. In M^Pro^, large structural variations were observed on residues N142 and Q189. All inhibitors were also characterized on their inhibition of M^Pro^ in 293T cells, which revealed their *in cellulo* potency that is drastically different from their *in vitro* enzyme inhibition potency. Inhibitors that showed high *in cellulo* potency all contain *O*-*tert*-butyl-threonine at the P3 site. Based on the current and a previous study, we conclude that *O*-*tert*-butyl-threonine at the P3 site is a key component to achieve high cellular and antiviral potency for peptidyl aldehyde inhibitors of M^Pro^. This finding will be critical to the development of novel antivirals to address the current global emergency of concerning the COVID-19 pandemic.

## INTRODUCTION

COVID-19 is the prevailing pandemic that has ravaged much of the world. As the COVID-19 pathogen, SARS-CoV-2 uses its membrane Spike protein to recognize the human receptor ACE2 for infection.^1-2^ Current COVID-19 vaccines all target this process for the neutralization of the virus. However, the continuous emergence of new viral strains that evade vaccines demands other antivirals to be developed as well. SARS-CoV-2 has a very large open reading frame ORF1ab that is translated to two large polypeptides pp1a and pp1b in the human cell host.^3-4^ The processing of pp1a and pp1b to 16 nonstructural proteins (nsps) that are functionally critical to the viral replication relies on proteolytic activities of two internal nsp fragments, nsp3 and nsp5.^5^ Nsp5 is also called 3C-like protease and more recently main protease (M^Pro^). Since M^Pro^ hydrolyzes 13 out of the total of 16 nsps, it has been considered as a viable target for the development of antivirals. In the past year, a number of papers have been published on the development of peptidyl aldehydes that contain β-(S-2-oxopyrrolidin-3-yl)-alaninal (opal) at the P1 position as potent M^Pro^ inhibitors.^6-14^ Other inhibitors were developed as well.^15-28^ However, systematic structure-activity relationship (SAR) studies of opal-based M^Pro^ inhibitors are needed. In the current study, we explore variations at the P2 and P3 sites and the *N*-terminal protection group in opal-based peptidyl M^Pro^ inhibitors. These inhibitors were characterized in their *in vitro* and *in cellulo* inhibition of M^Pro^ and their interactions with M^Pro^ by X-ray protein crystallography. Our study reveals that *O*-*tert*-butyl-threonine at the P3 site in an opal-based M^Pro^ inhibitor is critical to achieve high cellular and antiviral potency.

## RESULTS

### The Design and Synthesis of MPI11-28

In previous studies, we developed MPI1-10 and characterized both their M^Pro^ inhibition potency and their antiviral potency.^13-14^ As a tripeptidyl aldehyde, MPI8 shows the highest antiviral potency with an EC_50_ value of 30 nM to neutralize SARS-CoV-2 (USA-WA1/2020) in Vero E6 cells.^29^ MPI3, another tripeptidyl aldehyde doesn’t have high antiviral potency. However, it has the highest *in vitro* M^Pro^ inhibition potency with an IC_50_ value as 8.5 nM.^13^ To explore how substituents at different positions in a tripeptidyl opal-based M^Pro^ inhibitor (shown on the top of Figure 1) influence its potency, we decided to carry out a systematic SAR study. We maintained opal at the P1 position due to its established preferential binding to the M^Pro^ P1 binding pocket and its covalent adduct formation with C145, the M^Pro^ catalytic cysteine. M^Pro^ tolerates leucine and phenylalanine at the P2 site of a substrate. Therefore, we chose to vary this site in our inhibitor design with β-alkyl alanines with a size between or around leucine and phenylalanine. Chosen alkyl substituents are isopropyl, phenyl, cyclohexyl, t-butyl, isopropenyl, cycopropyl, 2-furyl, and 2-thienyl. M^Pro^ doesn’t have a binding pocket for the P3 residue in a substrate. However, native M^Pro^ substrates have valine or a similar size residue at this position. Based on known opal inhibitors and substrates of M^Pro^, we varied this site with amino acids including valine, *O*-*tert*-butyl-threonine, *L*-cyclopropylglycine, *L*-*tert*-butyl-glycine, *L*-*α*-methyl-valine, dimethylglycine, and 1-aminocyclopropane-1-carboxylate. The *N*-terminal protection group was chosen between CBZ, *m*-chloro CBZ, and indole-2-carboxylate due to their demonstrated contributions to antiviral potency.^6, 10^ A total of 18 new M^Pro^ inhibitors, designated as MPI11-28 shown in Figure 1, were designed. We synthesized all inhibitors according to the synthetic route shown in Scheme 1. In this synthesis, a P2 amino acid ester was conjugated with a *N*-protected P3 amino acid and the afforded product was then hydrolyzed to free its *C*-terminal carboxylate for reaction with β-(S-2-oxopyrrolidin-3-yl)-alanine ester. An obtained tripeptidyl ester was reduced to afford a *C*-terminal alcohol that was then oxidized via Dess-Martin oxidation in a mild condition to make a final product.

**Scheme 1.**
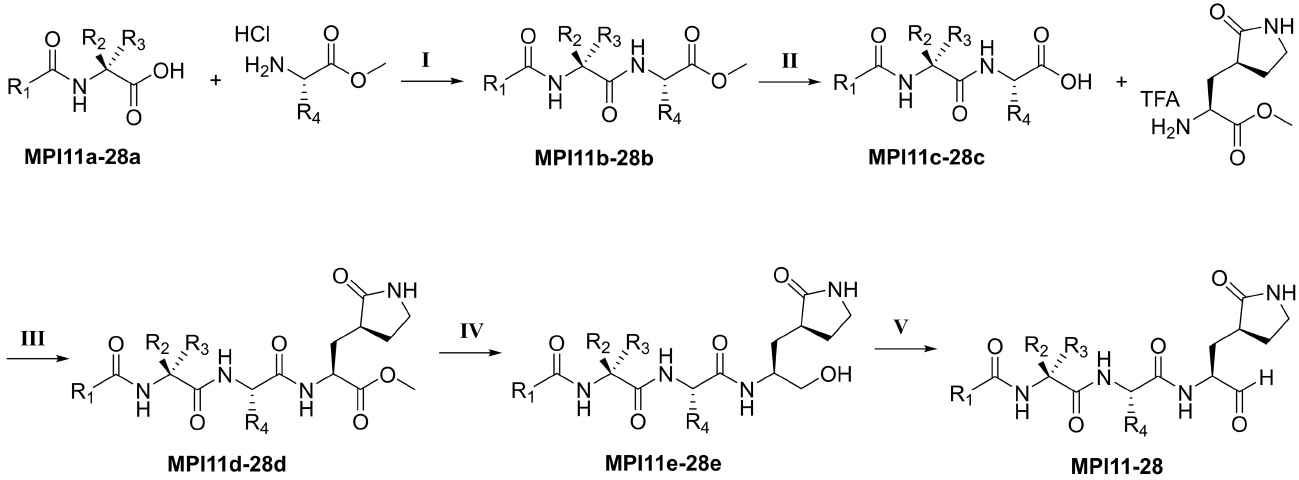
Reagents and conditions: (I) HATU, DIPEA, DMF; (II) LiOH•H_2_O, THF/H_2_O; (III) HATU, DIPEA, DMP; (IV) LiBH_4_, THF; (V) DMP, DCM.

**Figure 1.**
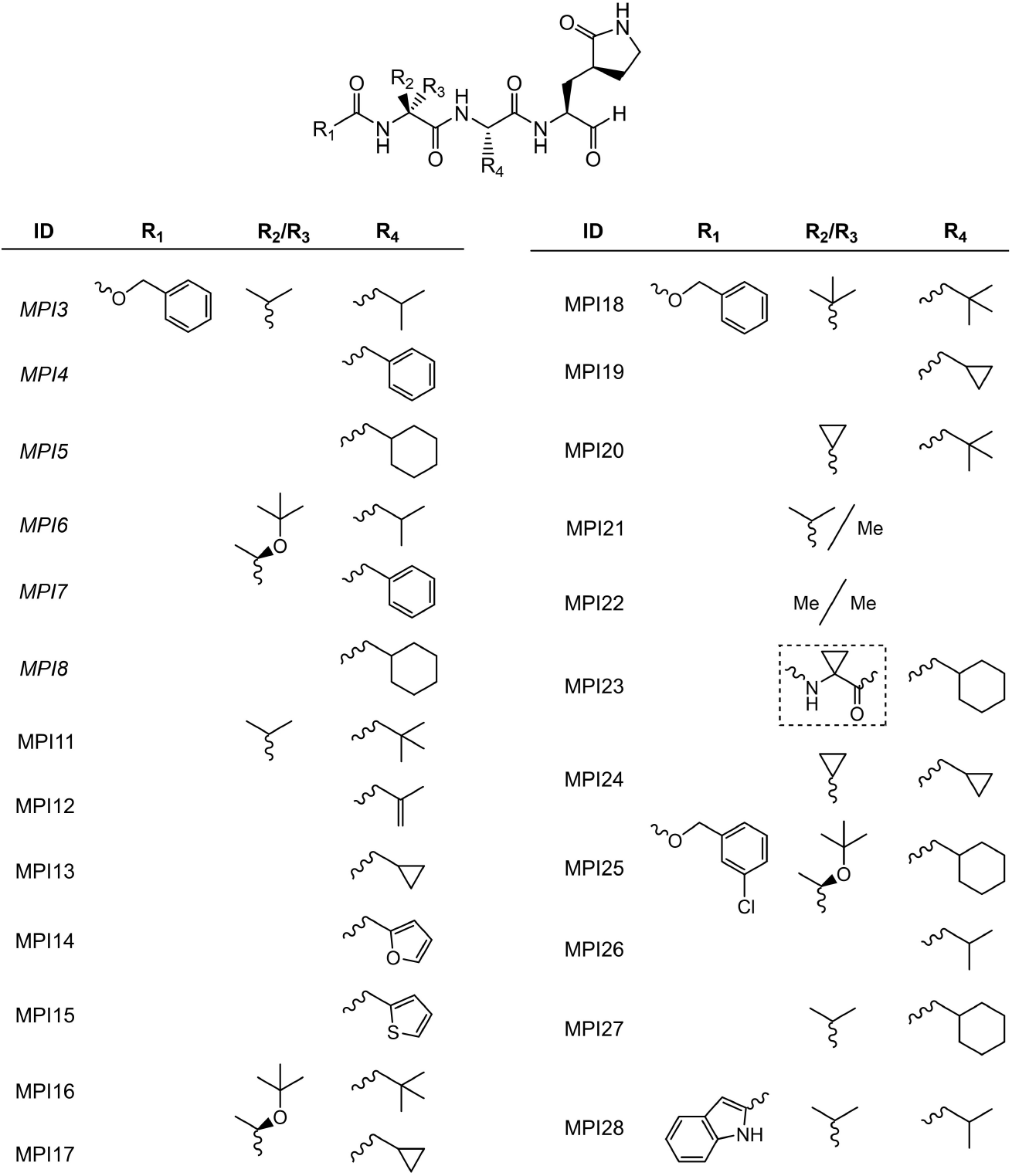
Structures of tripeptidyl M^Pro^ inhibitors MPI11-28. Inhibitors that have just a single component at R_2_/R_3_ shown contain a hydrogen at the R_3_ position. **MPI3-8** were previously developed and are shown for comparison.

### Kinetic characterizations of MPI11-28 on their enzymatic inhibition of M^Pro^

We followed a previously established protocol that uses Sub3, a fluorogenic peptide substrate of M^Pro^ to determine IC_50_ values for MPI11-28.^28^ In this assay, we preincubated M^Pro^ with an inhibitor for 30 min before Sub3 was added and the fluorescent product formation was recorded in a fluorescence plate reader. Except MPI15 that is insoluble in DMSO and therefore was not characterized, all other inhibitors displayed well traceable inhibition curves as shown in Figure 2. We fit all data to a four-parameter variable slope inhibition equation in GraphPad 9.0 to obtain IC_50_ values for all inhibitors. As shown in Table 1, MPI11-28 display a large variation of IC_50_ values that ranges from 4.8 to 650 nM. MPI11-14 all contain valine at the P3 site. They have the lowest IC_50_ values among all newly developed M^Pro^ inhibitors. MPI13-14 have determined IC_50_ values around 5 nM. Since 10 nM M^Pro^ is the lowest enzyme concentration we can use to do the inhibition analysis, 5 nM is technically the lowest IC_50_ value we can detect. Therefore, M^Pro^ inhibition potency for MPI13-14 is likely higher than what the numbers shown in Table 1 indicate. In comparison to MPI13-14, MPI11-12 have slightly higher IC_50_ values that are around 10 nM. In previous work, we developed another three M^Pro^ inhibitors MPI3-5 that also contain a valine at the P3 site. All three have low IC_50_ values.

**Table 1:** Determined enzymatic IC_50_ and cellular EC_50_ values of M^Pro^ inhibitors

| ID | Enzymatic IC<sub>50</sub> (nM) | Cellular EC<sub>50</sub> (μM) | Antiviral EC<sub>50</sub> (μM) | PDB Entry | ID | Enzymatic IC<sub>50</sub> (nM) | Cellular EC<sub>50</sub> (μM) | Antiviral EC<sub>50</sub> (μM) | PDB Entry |
| --- | --- | --- | --- | --- | --- | --- | --- | --- | --- |
| MPI3 | 8.5<sup>b</sup> | > 2<sup>c</sup> |  | 7JQ0 | MPI17 | 60 | 0.097 | 1.2<sup>e</sup>/1.8<sup>f</sup>/1.1<sup>g</sup> | - |
| MPI4 | 15<sup>b</sup> | > 2<sup>c</sup> |  | 7JQ1 | MPI18 | 36 | 0.86 | > 5<sup>e</sup>/0.71<sup>f</sup>/> 5<sup>g</sup> | 7RVR |
| MPI5 | 33<sup>b</sup> | 0.66<sup>c</sup> |  | 7JQ2 | MPI19 | 13 | 1.5 |  | 7RVS |
| MPI6 | 60<sup>b</sup> | 0.12<sup>c</sup> |  | 7JQ3 | MPI20 | 16 | > 2 |  | 7RVT |
| MPI7 | 47<sup>b</sup> | 0.19<sup>c</sup> |  | 7JQ4 | MPI21 | 77 | > 2 |  | 7RVU |
| MPI8 | 105<sup>b</sup> | 0.03<sup>c</sup> |  | 7JQ5 | MPI22 | 49 | > 10 |  | 7RVV |
| MPI11 | 8.8 | > 2 |  | 7RVM | MPI23 | 350 | > 10 |  | 7RVW |
| MPI12 | 11 | > 2 |  | 7RVN | MPI24 | 8.4 | > 2 |  | 7RVX |
| MPI13 | 4.8<sup>d</sup> | > 2 |  | 7RVO | MPI25 | 650 | 0.14 | 2.2<sup>e</sup>/1.6<sup>f</sup>/0.87<sup>g</sup> | 7RVY |
| MPI14 | 5.2<sup>d</sup> | > 2 |  | 7RVP | MPI26 | 530 | 0.23 | 1.9<sup>e</sup>/0.65<sup>f</sup>/1.6<sup>g</sup> | 7RVZ |
| MPI15<sup>a</sup> | n.d. | n.d. |  | - | MPI27 | 160 | 0.63 | > 5<sup>e</sup>/> 5<sup>f</sup>/3.2<sup>g</sup> | 7RW0 |
| MPI16 | 150 | 0.056 | 1.2<sup>e</sup>/1.2<sup>f</sup>/0.58<sup>g</sup> | 7RVQ | MPI28 | 52 | > 10 |  | 7RW1 |

<sup>a</sup>Its IC<sub>50</sub> and EC<sub>50</sub> values were not determined (n.d.) due to insolubility.
<sup>b</sup>Data were taken from ref. <sup>13</sup>.
<sup>c</sup>Data were taken from ref. <sup>29</sup>.
<sup>d</sup>Reaching the detection limit.
<sup>e</sup>Antiviral EC<sub>50</sub> value for the USA-WA1/2020 strain.
<sup>f</sup>Antiviral EC<sub>50</sub> value for the Beta strain.
<sup>g</sup>Antiviral EC<sub>50</sub> value for the Delta strain.

**Figure 2.**
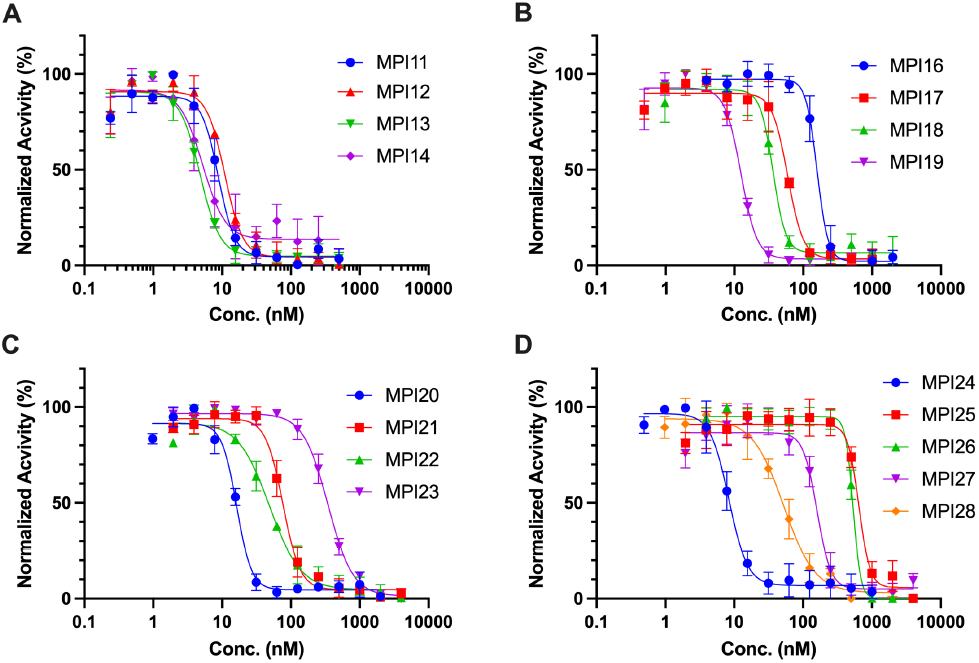
Inhibition curves of MPI11-28 on M^Pro^. Triplicate experiments were performed for each compound. For all experiments, 20 or 10 nM M^Pro^ was incubated with an inhibitor for 30 min before 10 μM Sub3 was added. The M^Pro^-catalyzed Sub3 hydrolysis rate was determined by measuring linear increase of product fluorescence (Ex: 336 nm /Em: 455 nm) for 5 min.

All 7 inhibitors that have valine at the P3 site and a *N*-terminal CBZ protection group are among the most potent M^Pro^ inhibitors *in vitro*. A P3 valine apparently favors the enzyme inhibition kinetics. A comparison of all seven inhibitors also reveals that a residue at the P2 site with a size close to leucine leads to better M^Pro^ inhibition. β–Cy-βclopropyl alanine and β-(furan-2yl) alanine that have a more rigid side chain than leucine favor the enzyme inhibition kinetics more. MPI6-8 that have a P3 *O*-*tert*-butyl-threonine were developed previously and showed high cellular and antiviral potency. We synthesized two more M^Pro^ inhibitors with a P3 *O*-*tert*-butyl-threonine, MPI16-17. Both have an IC_50_ value similar to MPI6-8. Varying the P2 position does not seem to significantly change *in vitro* M^Pro^ inhibition potency among this group of inhibitors. To explore further on whether variations at the P3 site lead to different inhibition potency, we developed MPI18-24. Variants include two dialkyl glycines and 1-aminocyclopropane-1-carboxylate that are not standard *L*-amino acids. Except MPI23 that contains a P3 1-aminocyclopropane-1-carboxylate, all other inhibitors have an IC_50_ value below 100 nM. The more rigid P3 residue in MPI23 likely contributes to its less favorable M^Pro^ inhibition kinetics. MPI24 is structurally similar to MPI13. Its enzyme inhibition potency is similar to MPI13 as well. Three inhibitors MPI25-27 that have a *N*-terminal *m*-chloro CBZ group were developed and characterized as well. In comparison to their regular CBZ-containing counterpart inhibitors MPI8, MPI6, and MPI5, they display more than 5-fold higher IC_50_ values. Apparently adding a *m*-chloro substituent to the terminal CBZ group doesn’t improve *in vitro* enzyme inhibition potency. MPI28 has a *N*-terminal indole-2-carboxylate. In comparison to its CBZ counterpart inhibitor MPI3, MPI28 has a 6-folder higher IC_50_ value. In comparison to indole-2-carboxylate, it is evident that CBZ serves as a better *N*-terminal group for improved *in vitro* enzyme inhibition potency.

### X-Ray Crystallography analysis of M^Pro^ bound with 16 different inhibitors

To characterize our developed inhibitors in their interactions with M^Pro^, we crystallized M^Pro^ in its apo form, soaked apo-M^Pro^ crystals with different inhibitors and then determined structures of M^Pro^ bound with different inhibitors using X-ray crystallography. Among 17 inhibitors that we used to soak crystals, 16 were observable in the active site of M^Pro^. All structures were determined in high resolutions (Table S1). 2F_o_-F_c_ electron density map around each inhibitor clearly shows a covalent interaction with C145 of M^Pro^ to generate a hemithioacetal (Figure 3A). In all formed hemithioacetals, the hemithioacetal alcohol takes an *S* configuration. In most structures, electron density around inhibitors is well shaped for modeling the P1 opal, P2, and P3 residues. For MPI12, the isopropenyl group shows weak electron density around its 1- and 3-carbon atoms indicating that the group is probably freely rotating around its 2-carbon atom in the active site of M^Pro^. Although a few M^Pro^-inhibitor complexes have electron density that can be used to model the *N*-terminal group, the majority have relatively weak electron density around the *N*-terminal group indicating weak binding of the *N*-terminal group at the active site. Superpositioning all M^Pro^-inhibitor complexes shows a similar binding mode for all inhibitors to the M^Pro^ active site (Figures 3B and 3C). MPI28 displays the most difference from other inhibitors due to its unique *N*-terminal indole-2-carboxylate. All other inhibitors show very similar binding to M^Pro^ except that different orientations are adopted at the relatively flexible *N*-terminal group. In all M^Pro^-inhibitor complexes, the protein shows a similar structure except at two residues N142 and Q189 and the M49-containing helix. Both N142 and Q189 adopt varied orientations in different M^Pro^-inhibitor structures and the M49-containing helix is structurally rearranged and not visible in most M^Pro^-inhibitor complexes. Due to a large number of crystal structures that are presented in this work, we will focus on only four representative M^Pro^-inhibitor complexes to describe structure variations of M^Pro^ at the active site. In M^Pro^-MPI11 as shown in Figure 3D, MPI11 forms extensive hydrogen bonds with M^Pro^. The hemithioacetal hydroxyl group forms two hydrogen bonds with two backbone amide nitrogen atoms explaining its exclusive *S* configuration. The opal side-chain amide forms three hydrogen bonds with E166, N142, and F140 and its backbone nitrogen forms a hydrogen bond with H164. The side chain of the P2 β-*tert*-butyl-alanine (4-methyl-leucine) involves van der Waals interactions with M^Pro^ residues surrounding the P2 binding pocket including M49, H41, M165, D187, and Q189. Its α-nitrogen interacts with the side chain of Q189 indirectly through a water molecule. The P3 residue of MPI11 forms two hydrogen bonds with backbone nitrogen and oxygen atoms of E166. The *N*-terminal CBZ of MPI11 interacts hydrophobically with the backbone of the Q189-containing loop and the side chain of P168. In M^Pro^-MPI11, the side chain of N142 flips away from the opal side chain of MPI11. MPI16 interacts with M^Pro^ similar to MPI11. Notable differences are at N142 and Q189. N142 forms van der Waals interactions with the opal oxopyrrolidine ring and Q189 forms a hydrogen bond directly with the backbone nitrogen of the P2 residue. In M^Pro^-MPI21, N142 adopts a conformation away from the opal side chain and Q189 forms a hydrogen bond with a CBZ carbamate oxygen atom. MPI28 has a different *N*-terminal group.

**Figure 3.**
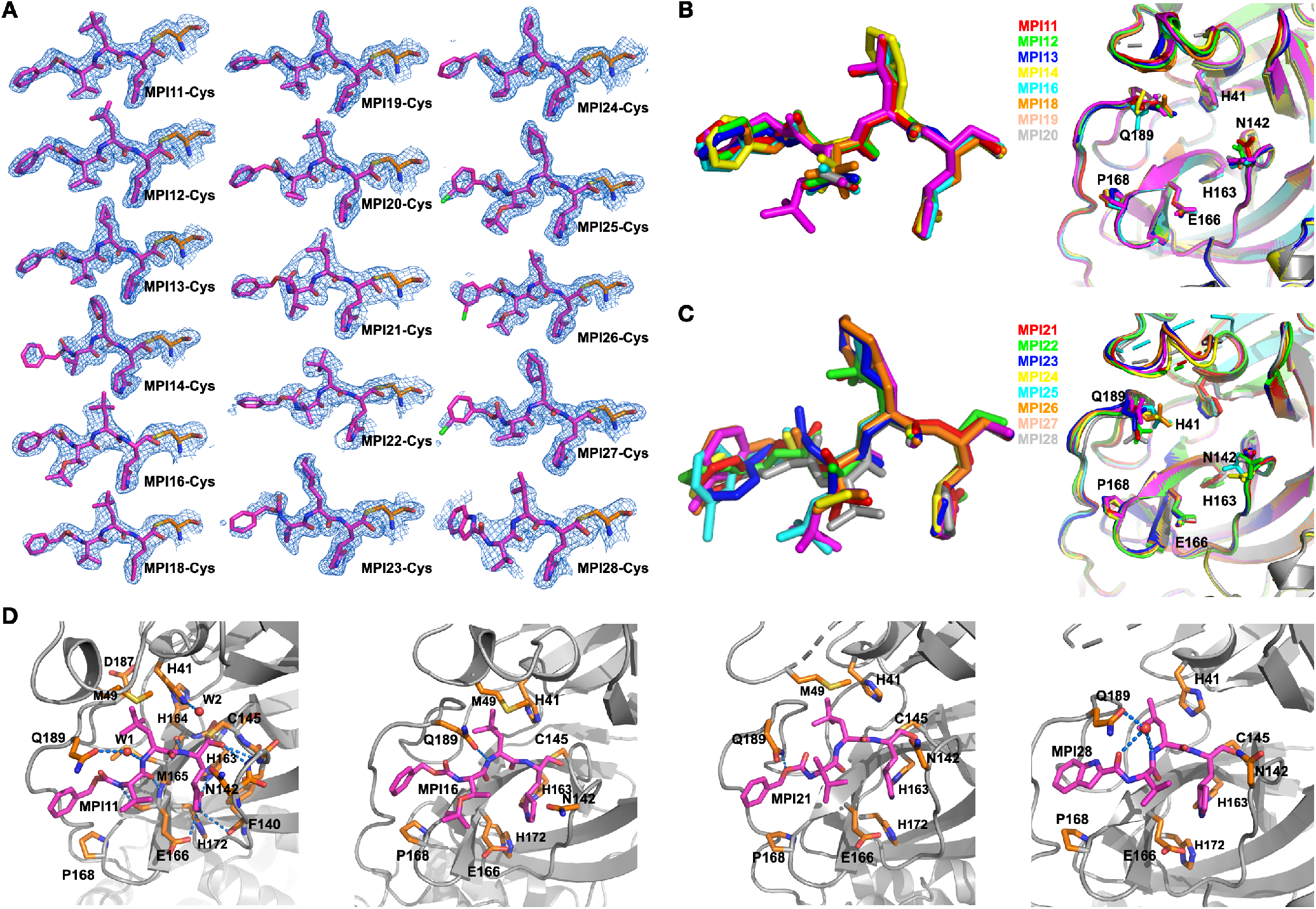
Crystal structures of M^Pro^ bound with 16 MPIs. (A) Contoured 2F_*o*_-F_*c*_ maps at the 1*σ* level around 16 MPIs and C145 in the active site of M^Pro^. (B) Superposition of M^Pro^ complexes with 8 MPIs including MPI11-14, MPI16, and MPI18-20. Inhibitors on the left and proteins on the right are presented separately for clarity. (C) Superposition of M^Pro^ complexes with MPI21-28. Inhibitors and proteins are shown separately. (D) Interactions between the inhibitor and active site residues in M^Pro^ complexes with MPI11, MPI16, MPI21, and MPI28. Some of hydrogen bonds between ligands and M^Pro^ are shown as blue dotted lines.

Interactions involving its *N*-terminal group are different from other inhibitors. The indole-2-carboxyl oxygen forms a hydrogen bond with a water molecule that also forms hydrogen bonds with Q189 and the backbone nitrogen of the P2 residue.

### Characterizations of Cellular M^Pro^ inhibition potency of MPI11-28

M^Pro^ is acutely toxic to a human cell host. Using this unique characteristic, we previously developed a cell-based analysis to characterize cellular M^Pro^ inhibition potency for M^Pro^ inhibitors.^29^ In this assay, an inhibitor with *in cellulo* potency suppresses cellular toxicity from the expression of an M^Pro^-eGFP fusion protein, which leads to enhanced overall expression of M^Pro^-eGFP that can be characterized by flow cytometry. We consider this assay more advantageous over a direct antiviral assay in the characterization of M^Pro^ inhibitors since a compound may inhibit host proteases such as TMPRSS2, furin, and cathepsin L that are critical for SARS-CoV-2 infection to provide false positive *in cellulo* potency results of M^Pro^ inhibition. *In cellulo* potency determined by this method for MPI5-8 agreed well with their antiviral potency that were characterized using SARS-CoV-2 (USA-WA1/2020) in Vero E6 cells. We adopted this assay to characterize MPI11-28 as well. All inhibitors were tested up to 10 μM in their inhibition of M^Pro^-eGFP in human 293T cells. Overall cellular eGFP fluorescence were plotted against the inhibition concentration to obtain their EC_50_ values. Results are presented in Figure 4 and Table 1. As shown in Figure 4A, MPI11-14 exhibit minimal *in cellulo* potency to inhibit M^Pro^, which is in significant contrast to their very high *in vitro* enzyme inhibition potency. Since their inhibition curves do not reach a plateau, their EC_50_ values are estimated as higher than 2 μM. MPI16-17 are the two most potent inhibitors among MPI11-28 on *in cellulo* potency with determined EC_50_ values as 56 nM and 97 nM, respectively (Figure 4B). In comparison to MPI16-17, MPI18-19 have weaker *in cellulo* potency with determined EC_50_ values as 860 nM and 1,500 nM, respectively. MPI20-23 all have weak *in cellulo* potency (Figure 4C). MPI22-23 show very low inhibition of M^Pro^-eGFP in 293T cells. At 10 μM, their driven M^Pro^-eGFP expression is lower than half of the plateau level of M^Pro^-eGFP observed for MPI16-19. For these four inhibitors, their EC_50_ values are estimated as higher than 2 μM for MPI20-21 and higher than 10 μM for MPI22-23. Among MPI24-28 (Figure 4D), MPI25-27 exhibit high *in cellulo* potency with EC_50_ values as 140 nM, 230 nM, and 630 nM respectively. On contrary to their high *in vitro* enzyme inhibition potency, MPI24 and MPI28 have very weak *in cellulo* potency with estimated EC_50_ values as higher than 2 μM and higher than 10 μM, respectively.

**Figure 4.**
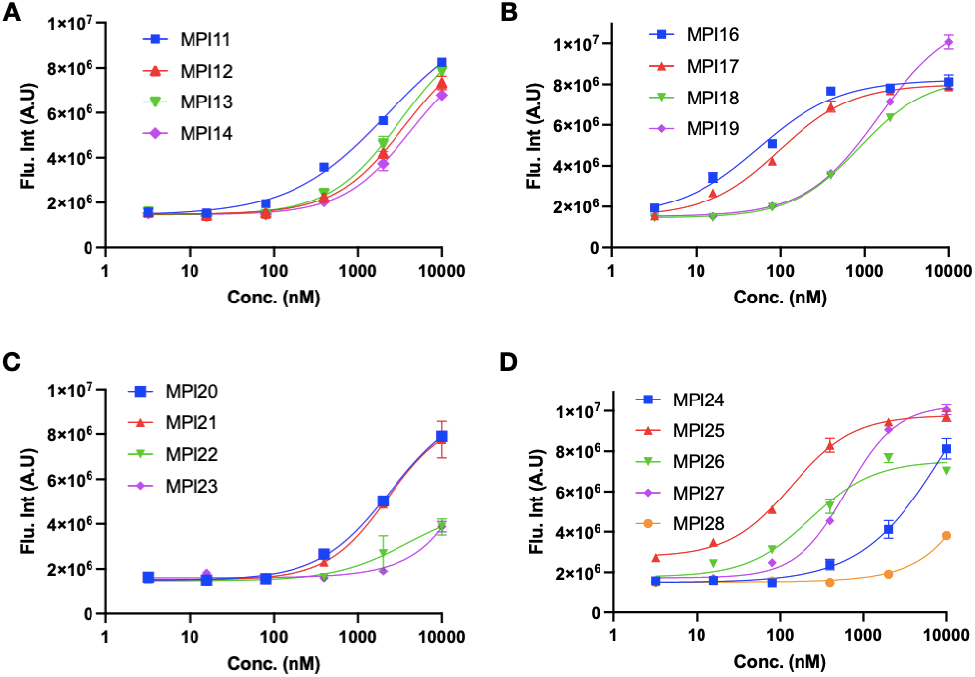
Cellular potency of MPI11-28 in their inhibition of M^Pro^ to drive host 293T cell survival and overall M^Pro^-eGFP expression.

**Figure 5.**
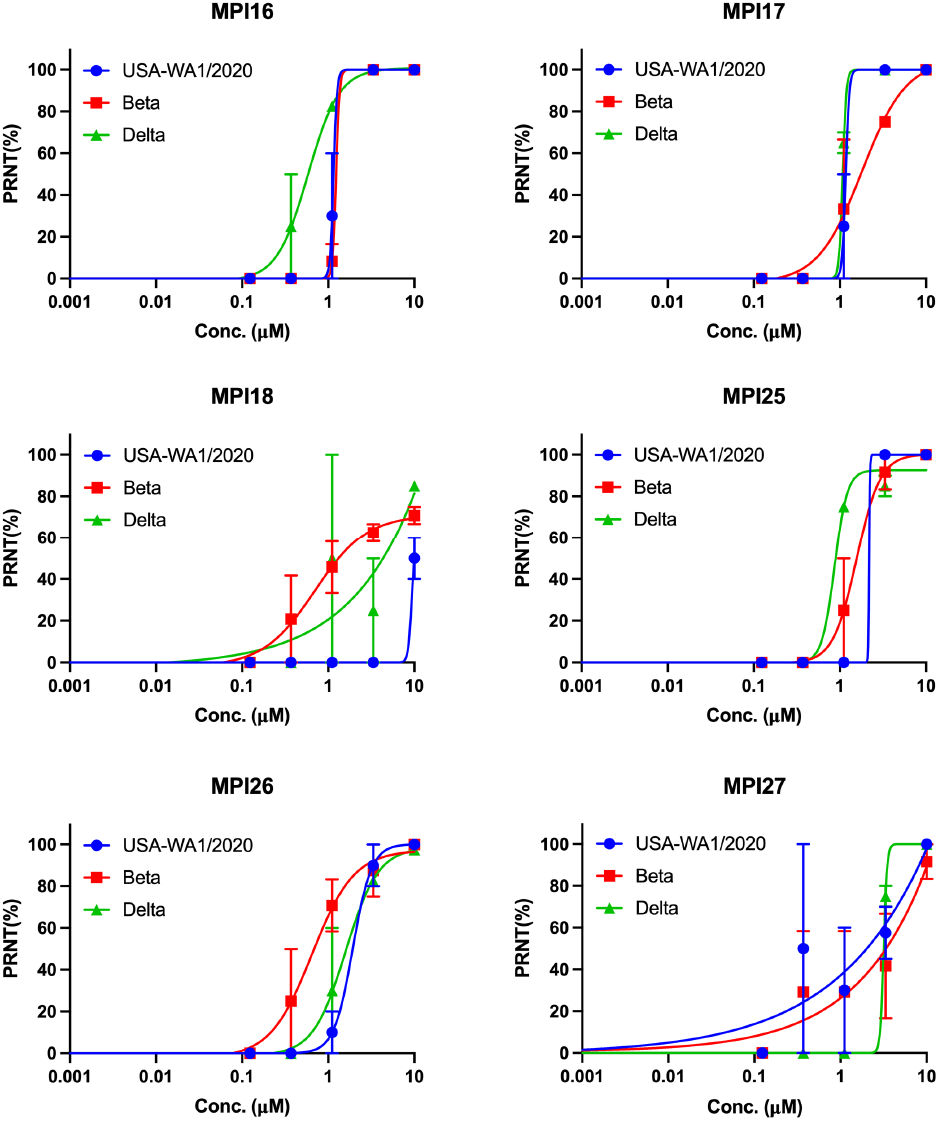
Plaque reduction neutralization tests (PRNTs) of MPI16-18 and MPI25-27 on their inhibition of three SARS-CoV-2 strains USA-WA1/2020, Beta and Delta in Vero E6 cells. Two repeats were conducted for each concentration.

### Characterizations of antiviral potency of six selected inhibitors on three SARS-CoV-2 variants

Four most cellularly potent inhibitors MPI16, MPI17, MPI25, and MPI26 that contain a P3 *O-tert-*butyl-threonine and two mildly cellularly potent inhibitors MPI18 and MPI27 that don’t contain a P3 *O-tert-*butyl-threonine were analyzed on their antiviral potency against three SARS-CoV-2 variants including USA-WA1/2020, Beta and Delta. We conducted plaque reduction neutralization tests in Vero E6 cells for all inhibitors. We infected Vero E6 cells by virus in the presence of an inhibitor at various concentrations for three days and then quantified viral plaque reduction. As presented in the attached figure and table, all four inhibitors that contain a P3 *O-tert-*butyl-threonine showed high antiviral potency with determined EC_50_ values around 1 μM. Except MPI8 on the Beta strain, MPI18 and MPI27 that don’t have a P3 *O-tert-*butyl-threonine showed only mild antiviral potency. These antiviral results support that P3 *O-tert-*butyl-threonine in a peptidyl aldehyde inhibitor leads to high antiviral potency.

## DISCUSSION

There are 11 unique proteolytic sites in pp1a and pp1b that are hydrolyzed by M^Pro^. Glutamine is a strictly required residue at the P1 site for all 11 sites. Due to this strict requirement and medicinal chemistry information learnt from the same function enzyme in SARS-CoV, most peptidyl inhibitors that have been developed for M^Pro^ have maintained a β-(S-2-oxopyrrolidin-3-yl)-alanine analog at the P1 site for improved potency. In our design, we keep opal at this site for all inhibitors. Our crystallography results show a well-shaped opal side chain in all M^Pro^-inhibitor complexes that fits neatly in the P1 binding pocket of the enzyme. Since opal is optimized for this site, there is small chemical space to manipulate its side chain for improved binding. In all determined structures, the side chain of N142 adopts different conformations that do not necessarily interact with the opal oxopyrrolidine ring. It may be possible to modify the opal oxopyrrolidine ring to introduce interactions such as hydrogen bonding with the side chain of N142 for improved binding. Among all 11 M^Pro^-targeted proteolytic sites, 9 have a P2 leucine, 1 has a P2 phenylalanine, and 1 has a P2 valine. The enzyme has an apparent preference for a P2 leucine in its substrates. A comparison of *in vitro* enzyme inhibition potency for all MPIs that we have developed also reveals that M^Pro^ prefers a P2 residue with a similar size as leucine in a peptidyl inhibitor for favorable binding. Converting the isopropyl group at the β-carbon of the P2 leucine to cyclopropyl and 2-furanyl improves the enzyme inhibition kinetics. Although changing the P2 leucine in an inhibitor to a larger residue such as phenylalanine and β-cyclohexylalanine diminishes *in vitro* M^Pro^ inhibition potency, the level is not significant.

M^Pro^ shows almost no preference to the P3 residue in its targeted proteolytic sites in pp1a and pp1b. We explored a variety of chemical variants at this site including dialkyl glycines and 1-aminocyclopropane-1-carboxylate that have no α-proton. Although M^Pro^ displays a clear preference for a P3 valine, other substituents at this site are well tolerated. Given the diverse structures of tolerable chemical compositions at this site, D-amino acids might be introduced at this site for the development of novel M^Pro^ inhibitors. This will need to be explored further. All our crystal structures show weak electron density around the *N*-terminal group indicating its weak binding to the enzyme. Adding a *m*-chloride to the *N*-terminal CBZ group also leads to significant decrease in *in vitro* enzyme inhibition potency. Our study indicates that both CBZ and indole-2-carboxyl are not optimal chemical groups for interactions with M^Pro^. Some smaller groups need to be explored at this site.

Although most inhibitors we have developed in this study exhibit very high *in vitro* enzyme inhibition potency and two inhibitors have IC_50_ values reaching the characterization limit, just a few inhibitors show *in cellulo* potency to inhibit M^Pro^ in 293T cells. Inhibitors that have a P3 *O*-*tert*-butyl-threonine all show high *in cellulo* potency. All other P3 substituents lead to low *in cellulo* potency. Many factors may contribute to this phenomenon. Since all MPIs are peptidyl inhibitors, a small or native P3 residue might be prone to proteolytic degradation by host proteases leading to low *in cellulo* potency. This is partially supported by characterizable *in cellulo* potency for MPI18-19 that have a P3 β-*tert*-butyl-alanine. It is also possible that a P3 *O*-*tert*-butyl-threonine introduces more favorable cellular permeability than other P3 substituents into an M^Pro^ inhibitor. MPI21-23 that have a P3 dialkylglycine are expected to be more resistant to proteolytic digestion by human proteases than other MPIs. Their low *in cellulo* potency might be due to low cellular permeability. Based on *in cellulo* potency of MPIs, we can also derive that an *N*-terminal CBZ group works better than the other two groups that have been tested at this site and a P2 -cyclohexylalanine favors high *in cellulo* potency.

## CONCLUSIONS

In combination with our previous studies,^13,29^ we can conclude that a P3 *O*-*tert*-butyl-threonine in peptidyl aldehyde inhibitors for M^Pro^ is optimal for high *in cellulo* and antiviral potency. This new and critical finding will be highly useful for the development of novel antiviral lead series and drug candidates with potential to treat COVID-19. However, chemical compositions at the P2 site and the *N*-terminal group in peptidyl aldehyde inhibitors for optimal *in cellulo* and antiviral potency need to be explored further.

## EXPERIMENTAL SECTION

## ASSOCIATED CONTENT

Supporting Information.

The Supporting Information is available free of charge on the ACS Publications website.

Supplementary information for the synthesis of MPI11-28, NMR spectrascopies and HPLC chromatographies of MPI11-28, X-ray data collection and processing parameters, and all original flow cytometry graphs. All compounds are >95% pure by HPLC analysis.

## ACKNOWLEDGMENT

This work was supported by Welch Foundation (grant A-1715), DHHS-NIH-National Institute of Allergy and Infectious Diseases (R21AI164088), TAMU COS Strategic Transformative Research Program, and Texas A&M X Grants. The ALS-ENABLE beam-lines are supported in part by the National Institutes of Health, National Institute of General Medical Sciences, grant P30 GM124169-01 and the Howard Hughes Medical Institute. The Advanced Light Source is a Department of Energy Office of Science User Facility under Contract No. DE-AC02-05CH11231.

---

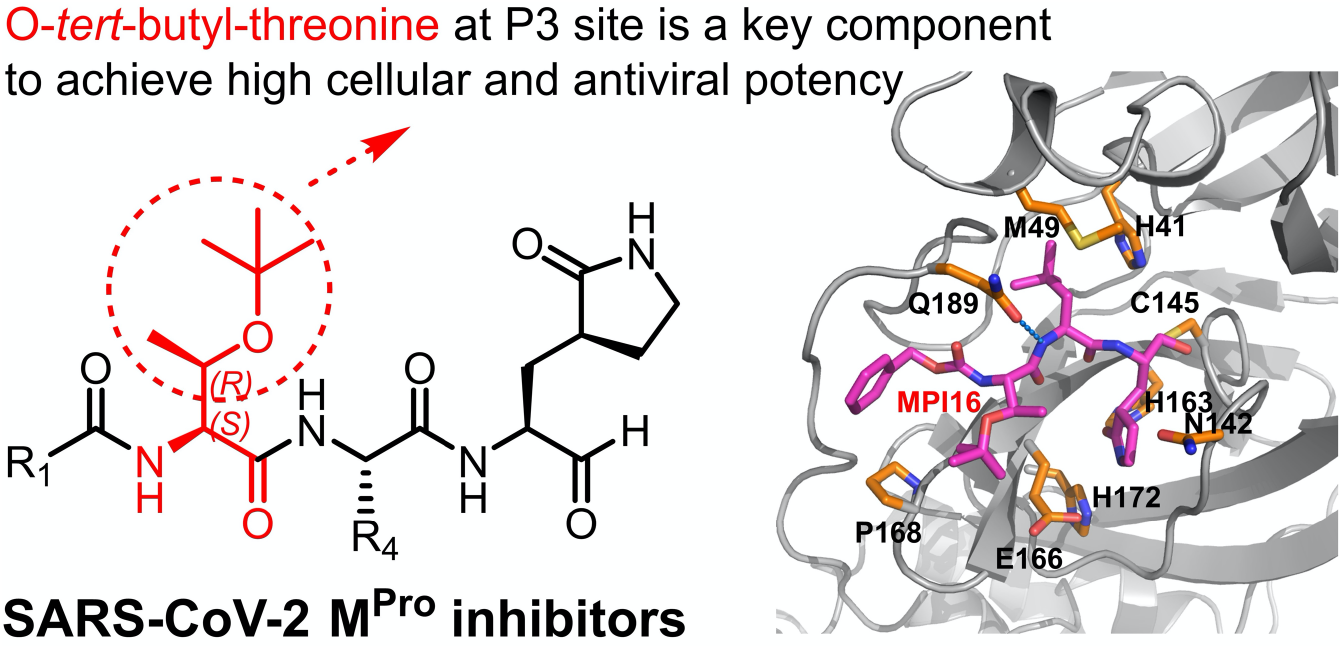

