## Supplementary Material for "The P3 *O*-*Tert*-Butyl-Threonine is Key to High Cellular and Antiviral Potency for Aldehyde-Based SARS-CoV-2 Main Protease Inhibitors"

for

**Materials.** We purchased yeast extract from Thermo Fisher Scientific, tryptone from Gibco, Sub3 from Bachem, HEK 293T/17 cells from ATCC, DMEM with GlutaMax from Gibco, FBS from Gibco, polyethyleneimine from Polysciences, the trypsin-EDTA solution from Gibco. Chemicals used in this work were acquired from Sigma Aldrich, Chem Impex, Ambeed, A2B, etc.

**$M^{Pro}$ Expression and Purification.** The expression plasmid pET28a-His-SUMO-$M^{Pro}$ was constructed in a previous study. We used this construct to transform *E. coli* BL21(DE3) cells. A single colony grown on a LB plate containing 50 μg/mL kanamycin was picked and grown in 5 mL LB media supplemented with 50 μg/mL kanamycin overnight. We inoculated this overnight culture to 6 L 2YT media with 50 μg/mL kanamycin. Cells were grown to $OD_{600}$ as 0.8. At this point, we added 1 mM IPTG to induce the expression of His-SUMO-$M^{Pro}$. Induced cells were let grown for 3 h and then harvested by centrifugation at 12,000 rpm, 4 °C for 30 min. We resuspended cell pellets in 150 mL lysis buffer (20 mM Tris-HCl, 100 mM NaCl, 10 mM imidazole, pH 8.0) and lysed the cells by sonication on ice. We clarified the lysate by centrifugation at 16,000 rpm, 4 °C for 30 min. We decanted the supernatant and mixed with Ni-NTA resins (GenScript). We loaded the resins to a column, washed the resins with 10 volumes of lysis buffer, and eluted the bound protein using elution buffer (20 mM Tris-HCl, 100 mM NaCl, 250 mM imidazole, pH 8.0). We exchanged buffer of the elute to another buffer (20 mM Tris-HCl, 100 mM NaCl, 10 mM imidazole, 1 mM DTT, pH 8.0) using a HiPrep 26/10 desalting column (Cytiva) and digested the elute using 10 units SUMO protease overnight at 4 °C. The digested elute was subjected to Ni-NTA resins in a column to remove His-tagged SUMO protease, His-tagged SUMO tag, and undigested His-SUMO-$M^{Pro}$. We loaded the flow-through onto a Q-Sepharose column and purified $M^{Pro}$ using FPLC by running a linear gradient from 0 to 500 mM NaCl in a buffer (20 mM Tris-HCl, 1 mM DTT, pH 8.0). Fractions eluted from the Q-Sepharose column was concentrated and loaded onto a HiPrep 16/60 Sephacryl S-100 HR column and purified using a buffer containing 20 mM Tris-HCl, 100 mM NaCl, 1 mM DTT, and 1 mM EDTA at pH 7.8. The final purified was concentrated and stored in a -80 °C freezer.

***In Vitro* $M^{Pro}$ Inhibition Potency Characterizations of MPIs.** For most MPIs, we conducted the assay using 20 nM $M^{Pro}$ and 10 □M Sub3. For MPI13-14, 10 nM $M^{Pro}$ was used. We dissolved all inhibitors in DMSO as 10 mM stock solutions. Sub3 was dissolved in DMSO as a 1 mM stock solution and diluted 100 times in the final assay buffer containing 10 mM $Na_xH_yPO_4$, 10 mM NaCl, 0.5 mM EDTA, and 1.25% DMSO at pH 7.6. We incubated $M^{Pro}$ and an inhibitor in the final assay buffer for 30 min before adding the substrate to initiate the reaction catalyzed by $M^{Pro}$. The production format was monitored in a fluorescence plate reader with excitation at 336 nm and emission at 455 nm. More assay details can be found in a previous study.[1]

**X-Ray Crystallography Analysis of $M^{Pro}$-Inhibitor Complexes.** The production of crystals of $M^{pro}$-inhibitor complexes wasfollowing the previous protocols.[1] The data of $M^{pro}$ with MPI11, MPI12 and MPI24 were collected on a Rigaku R-AXIS IV++ image plate detector. The data of $M^{pro}$ with MPI16, MPI18, MPI19, MPI22, MPI23, MPI25 and MPI27 were collected on a Bruker Photon II detector. The data of $M^{pro}$ with MPI14, MPI20, MPI21, MPI26 and MPI28 were collected at the Advanced Light Source (ALS) beamline 5.0.2 using a Pilatus3 6M detector. The diffraction data were indexed, integrated and scaled with iMosflm or PROTEUM3.[2] All the structures were determined by molecular replacement using the structure model of the free enzyme of the SARS-CoV-2 $M^{pro}$ [Protein Data Bank (PDB) ID code 7JPY] as the search model using Phaser in the Phenix package.[1,3] *JLigand* and *Sketcher* from the CCP4 suite were employed for the generation of PDB and geometric restraints for the inhibitors. The inhibitors were built into the $F_O$-$F_C$ density by using *Coot*.[4] Refinement of all the structures was performed with Real-space Refinement in Phenix.[3] Details of data quality and structure refinement are summarized in Table S1. All structural figures were generated with PyMOL (https://www.pymol.org).

***In cellulo* $M^{Pro}$ Inhibition Potency Characterizations of MPIs.** We grew HEK 293T/17 cells in high-glucose DMEM with GlutaMAX supplement and 10% FBS in 10 cm culture plates under 37 °C and 5% $CO_2$ to 80-90% confluency and then transfected cells with the pLVX-$M^{Pro}$-eGFP-2 plasmid. 30 mg/mL polyethyleneimine and the total of 8 μg of the plasmid in 500 μL opti-MEM media were used for transfection. We incubated transfected cells overnight. On the second day, we collected cells using 0.05% trypsin-EDTA to detach them from plates, resuspended collected cells in the original growth media, adjusted the cell density to $5 \times 10^5$ cells/mL, added 500 μL adjusted cells to each well of a 48-well plate, and then added 100 μL of a drug solution in DMEM. We incubated treated cells under 37 °C and 5% $CO_2$ for 72 h. After 72 h incubation, cells were collected using trypsinization and centrifugation. We resuspended collected cells in 200 μL PBS and analyzed cells with fluorescence using a Cytoflex Research Flow Cytometer based on the size scatters (SSC-A and SSC-H) and forward scatter (FSC-A). We gated cells based on SSC-A and FSC-A then with SSC-A and SSC-H. Fluorescence was detected with excitation at 488 nm and emission at 525 nm. All collected data were converted to csv files and analyzed using a self-prepared MATLAB script for massive data processing. We sorted the FITC-A column from lowest to highest. A $10^6$ cutoff was set to separate the column to two groups with higher than $10^6$ as positive and lower than $10^6$ as negative. We integrated the positive group and divided the total integrated fluorescence intensity by the total cell positive cell counts as Flu. Int. shown in all graphs. The standard deviation of positive fluorescence was calculated as well. All processed data were plotted and fitted to a four-parameter Hill equation in GraphPad 9.0 to obtain determined $EC_{50}$ values.

**The Synthesis of MPIs.** All reagents and solvents for the synthesis were purchased from commercial sources and used without purification. All glassware was flame-dried prior to use. Thin-layer chromatography (TLC) was carried out on aluminum plates coated with 60 F254 silica gel. TLC plates were visualized under UV light (254 or 365 nm) or stained with 5% phosphomolybdic acid. Normal phase column chromatography was carried out using a Yamazen Small Flash AKROS system. Analytical reverse phase HPLC was carried out on a Shimazu LC20 HPLC system with an analytical C18 column. Semipreparative HPLC was carried out the same system with a semipreparative C18 column. The mobile phases were $H_2O$ with 0.1% formic acid (A) and acetone with 0.1% formic acid (B). NMR spectra were recorded on a Bruker AVANCE Neo 400 MHz or Varian INOVA 300 MHz spectrometer in specified deuterated solvents. High-resolution electrospray mass spectrometry was carried out on a Thermo Scientific Q Exactive Focus system. The purity of all compounds were confirmed by NMR and analytic HPLC-UV as ≥ 95%.

**HPLC analysis of MPIs.** All compounds were determined by using Thermo Scientific ultimate 3000 HPLC with binary pumps, using Acclaim 120 C18 column (2.1x150 mm, 5 μL). All compounds were analyzed using (MeOH/$H_2O$ 0.1% Formic acid)(v/v)(0.3mL/min) and calculated the peak areas at 254 or 214 nm.

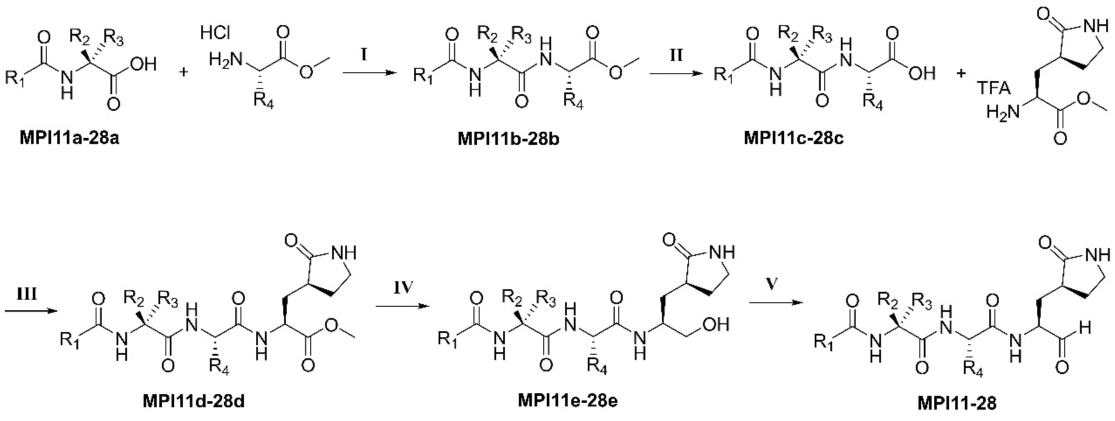


**Scheme S1.** General synthetic route to MPI11-28. Reagents and conditions: (I) HATU, DIPEA, DMF; (II) LiOH•$H_2O$, THF/$H_2O$; (III) HATU, DIPEA, DMP; (IV) $LiBH_4$, THF; (V) DMP, DCM

**(S)-methyl 2-((S)-2-(((benzyloxy)carbonyl)amino)-3-methylbutanamido)-4,4-dimethylpentanoate (MPI11b).** Methyl (S)-2-amino-4,4-dimethylpentanoate hydrochloride (0.4 g, 2.27 mmol, 1.1 equiv) and *N*-Cbz-L-valine (0.52 g, 2.07 mmol, 1.0 equiv) were dissolved in dry DMF (20 mL) and the reaction was cooled to 0 °C. HATU (1.01 g, 2.69 mmol, 1.3 equiv) and DIPEA (1.46 mL, 8.28 mmol, 4.0 equiv) were added, and the reaction mixture was allowed warm up to room temperature and stirred for 12 h. The mixture was then poured into water (50 mL) and extracted with ethyl acetate (4×20 mL). The organic layer was washed with aqueous hydrochloric acid 10% v/v (2×20 mL), saturated aqueous $NaHCO_3$ (2×20 mL), brine (2×20 mL) and dried over $Na_2SO_4$. The organic phase was evaporated to dryness and the crude material purified by silica gel column chromatography (15-50% EtOAc in n-hexane as the eluent) to afford pure **MPI11b** as a white solid (0.6 g, 73%). $^1$H NMR (400 MHz, Chloroform-*d*) δ 7.36 (m, 5H), 6.39 (m, 1H), 5.43 (t, *J* = 10.6 Hz, 1H), 5.12 (s, 2H), 4.65 (td, *J* = 8.5, 3.6 Hz, 1H), 4.04 (q, *J* = 7.2 Hz, 1H), 3.73 (s, 3H), 2.24 – 2.01 (m, 1H), 1.80 (dd, *J* = 14.2, 3.5 Hz, 1H), 1.50 (dd, *J* = 14.3, 8.7 Hz, 1H), 1.07 – 0.90 (m, 15H). $^{13}$C NMR (100 MHz, Chloroform-*d*): δ 173.5, 170.9, 156.4, 136.26, 128.5, 128.2, 128.0, 77.4, 77.0, 76.7, 67.0, 60.3, 52.3, 49.8, 46.0, 31.1, 30.7, 29.5, 19.1, 17.9.

**(S)-2-((S)-2-(((benzyloxy)carbonyl)amino)-3-methylbutanamido)-4,4-dimethylpentanoic acid (MPI11c).** The **MPI11b** (600 mg, 1.53 mmol, 1.0 equiv) was dissolved in THF/$H_2O$ (1:1, 10 mL). LiOH•$H_2O$ (153 mg, 3.82 mmol, 2.5 equiv) was added at 0 °C. The mixture was stirred at room temperature overnight. Then THF was removed on vacuum and the aqueous layer was acidified with 1 M HCl and extracted with dichloromethane (3 x 10 mL). The organic layer was dried over anhydrous $Na_2SO_4$ and concentrated to yield **MPI11c** as a white solid (415 mg, yield 70%). $^1$H NMR (400 MHz, Chloroform-*d*) δ 9.42 (s, 1H), 7.26 (tdd, *J* = 9.1, 5.5, 2.5 Hz, 5H), 6.69 (d, *J* = 8.2 Hz, 1H), 5.82 (d, *J* = 9.3 Hz, 1H), 5.10 – 4.96 (m, 2H), 4.54 (td, *J* = 8.6, 3.2 Hz, 1H), 4.13 – 3.88 (m, 1H), 1.83 – 1.73 (m, 1H), 1.43 (dd, *J* = 14.7, 8.9 Hz, 1H), 1.18 (dt, *J* = 12.4, 7.1 Hz, 1H), 0.91 (s, 3H), 0.85 (m, 12H).

**(5S,8S,11S)-methyl 5-isopropyl-8-neopentyl-3,6,9-trioxo-11-(((S)-2-oxopyrrolidin-3-yl)methyl)-1-phenyl-2-oxa-4,7,10-triazadodecan-12-oate (MPI11d).** The methyl (S)-2-amino-3-((S)-2-oxopyrrolidin-3-yl)propanoate hydrochloride (110 mg, 0.494 mmol, 1.1 equiv) and the **MPI11c** (170 mg, 0.45 mmol, 1.0 equiv) were dissolved in anhydrous DMF (10 mL) and the reaction was cooled to 0 °C. HATU (222 mg, 0.585 mmol, 1.3 equiv) and DIPEA (0.32 mL, 1.81 mmol, 4.0 equiv) were added, and the reaction mixture was allowed to warm up to room temperature and stirred for 12 h. The mixture was then poured into water (20 mL) and extracted with ethyl acetate (4×20 mL). The organic layer was washed with aqueous hydrochloric acid 10% v/v (2×20 mL), saturated aqueous $NaHCO_3$ (2×20 mL), brine (2×20 mL) and dried over $Na_2SO_4$. The organic phase was evaporated to dryness and the crude material purified by silica gel column chromatography to afford pure **MPI11d** as a white solid (200 mg, yield 81%). $^1$H NMR (400 MHz, Chloroform-*d*): δ 7.72 (d, *J* = 7.7 Hz, 1H), 7.32 – 7.21 (m, 5H), 6.88 – 6.60 (m, 2H), 5.47 (d, *J* = 9.2 Hz, 1H), 5.00 (s, 2H), 4.58 - 4.56 (m, 1H), 4.46 – 4.43 (m, 1H), 3.91 – 3.88 (m, 1H), 3.61 (s, 3H), 3.27 – 3.24 (m, 2H), 2.34 – 2.24 (m, 2H), 2.15 – 2.12 (m, 1H), 2.01 – 1.97 (m, 1H), 1.78 –1.75 (m, 3H), 1.19 – 1.11 (m, 1H), 0.91 – 0.81 (m, 16H).

**Benzyl ((S)-1-(((S)-1-(((S)-1-hydroxy-3-((S)-2-oxopyrrolidin-3-yl)propan-2-yl)amino)-4,4-dimethyl-1-oxopentan-2-yl)amino)-3-methyl-1-oxobutan-2-yl)carbamate (MPI11e).** To a stirred solution of **MPI11d** (150 mg, 0.27 mmol, 1.0 equiv) in anhydrous THF (8 mL) was added $LiBH_4$ (2.0 M in THF, 0.412 mL, 0.54 mmol, 2.0 equiv) in several portions at 0 °C. The reaction mixture was then allowed to warm up to room temperature and stirred for an additional 2 h. The reaction was quenched by the dropwise addition of 1.0 M HCl (aq) (1.2 mL) with cooling in an ice bath. Removed THF in vacuo, and the mixture was diluted with $H_2O$ and extracted with EtOAc, washed with sat. NaCl, dried over $Na_2SO_4$ and concentrated. The residue was purified by column chromatography (6% MeOH in $CH_2Cl_2$ as the eluent) to afford the pure product **MPI11e** as a white solid (102 mg, yield 71%). $^1$H NMR (400 MHz, Chloroform-*d*): δ 7.84 (d, *J* = 9.3 Hz, 1H), 7.71 (d, *J* = 7.5 Hz, 1H), 7.28 – 7.24 (m, 5H), 6.86 (d, *J* = 4.4 Hz, 1H), 5.66 (d, *J* = 8.7 Hz, 1H), 5.07 – 4.94 (m, 2H), 4.60 – 4.58 (m, 1H), 4.08 (t, *J* = 8.2 Hz, 1H), 3.89 (dq, *J* = 11.9, 5.6 Hz, 1H), 3.77 (s, 1H), 3.59 (d, *J* = 10.9 Hz, 1H), 3.32 – 3.06 (m, 2H), 2.46 – 2.14 (m, 3H), 1.99 – 1.96 (m, 2H), 1.72 – 1.69 (m, 2H), 1.64 – 1.40 (m, 2H), 0.94 – 0.77 (m, 15H). $^{13}$C NMR (100 MHz, $CDCl_3$):

δ 180.8, 173.6, 171.0, 156.6, 136.2, 128.5, 128.2, 128.0, 67.1, 65.3, 60.3, 51.1, 50.4, 46.6, 40.6, 38.2, 32.2, 31.4, 30.6, 29.6, 28.2, 19.1, 18.3.

**Benzyl ((S)-1-(((S)-4,4-dimethyl-1-oxo-1-(((S)-1-oxo-3-((S)-2-oxopyrrolidin-3-yl)propan-2-yl)amino)pentan-2-yl)amino)-3-methyl-1-oxobutan-2-yl)carbamate (MPI11).** To a solution of **MPI11e** (90 mg, 0.17 mmol, 1.0 equiv) in anhydrous $CH_2Cl_2$ (6 mL) was added $NaHCO_3$ (60 mg, 0.68 mmol, 4 equiv) and the Dess-Martin reagent (225 mg, 0.51 mmol, 3 equiv). The resulting mixture was stirred at RT for 12 h. Then the reaction was quenched with a saturated $NaHCO_3$ solution containing 10 % $Na_2S_2O_3$. The layers were separated. The organic layer was then washed with saturated brine solution, dried over anhydrous $Na_2SO_4$ and concentrated *on vacuum*. The residue was purified by column chromatography (6% MeOH in $CH_2Cl_2$ as the eluent) to afford the pure product **MPI11** as a white solid (58 mg, yield 65%). $^1$H NMR (400 MHz, Chloroform-*d*): δ 9.42 (s, 1H), 7.32 – 7.23 (m, 5H), 6.64 – 6.41 (m, 1H), 5.91 (d, *J* = 26.3 Hz, 1H), 5.43 (d, *J* = 8.9 Hz, 1H), 5.31 (d, *J* = 8.6 Hz, 1H), 5.03 (d, *J* = 7.8 Hz, 2H), 4.51 (dt, *J* = 8.6, 4.4 Hz, 1H), 4.26 (d, *J* = 7.0 Hz, 1H), 4.18 – 4.01 (m, 1H), 3.92 (dd, *J* = 8.4, 6.3 Hz, 1H), 3.29-3.26 (m, 2H), 2.44 – 2.25 (m, 2H), 2.16 – 2.05 (m, 1H), 1.91 – 1.87 (m, 2H), 1.80 – 1.73 (m, 1H), 1.58 – 1.54 (m, 2H), 1.02 – 0.73 (m, 15H). $^{13}$C NMR (100 MHz, Chloroform-*d*): δ 199.5, 180.0, 173.8, 171.1, 156.6, 136.2, 128.5, 128.2, 128.0, 67.1, 60.6, 57.2, 50.9, 46.4, 40.6, 37.9, 31.0, 30.6, 29.9, 29.6, 28.2, 19.2, 18.1.

**(S)-methyl 2-((S)-2-(((benzyloxy)carbonyl)amino)-3-methylbutanamido)-4-methylpent-4-enoate (MPI12b). MPI112b** was prepared with methyl (S)-2-amino-4-methylpent-4-enoate hydrochloride and *N*-Cbz-L-valine as a white solid following a similar procedure to **MPI11b** (yield 40%). $^1$H NMR (400 MHz, Chloroform-*d*): δ 7.41-7.28 (m, 5H), 6.19 (t, *J* = 5.8 Hz, 1H), 5.33 (d, *J* = 9.2 Hz, 1H), 5.11 (s, 2H), 4.83 (s, 1H), 4.73 (s, 1H), 4.67 (td, *J* = 8.0, 5.6 Hz, 1H), 4.08 – 3.95 (m, 1H), 3.72 (s, 3H), 2.55 (dd, *J* = 14.1, 5.4 Hz, 1H), 2.40 (dd, *J* = 14.0, 8.4 Hz, 1H), 2.14 (dt, *J* = 13.4, 7.0 Hz, 1H), 1.71 (s, 3H), 1.69 -1.66 (m, 1H), 0.95 (dd, *J* = 18.8, 6.7 Hz, 6H). $^{13}$C NMR (100 MHz, Chloroform-*d*): δ 172.3, 170.9, 156.3, 140.2, 136.2, 128.6, 128.2, 128.1, 114.9, 67.1, 60.2, 52.3, 50.5, 40.4, 31.2, 21.8, 21.1, 19.1, 17.7.

**(S)-2-((S)-2-(((benzyloxy)carbonyl)amino)-3-methylbutanamido)-4-methylpent-4-enoic acid (MPI12c). MPI12c** was prepared as a white solid following a similar procedure to **MPI11c** (yield 83%). $^1$H NMR (400 MHz, Chloroform-*d*): δ 7.41 – 7.28 (m, 5H), 6.41 (d, *J* = 7.3 Hz, 1H), 5.44 (d, *J* = 9.1 Hz, 1H), 5.11 (s, 2H), 4.83 (s, 1H), 4.75 (s, 1H), 4.73 – 4.63 (m, 1H), 2.62 (dd, *J* = 14.2, 5.2 Hz, 1H), 2.43 (dd, *J* = 14.2, 8.7 Hz, 1H), 1.94 – 1.84 (m, 1H), 1.72 (s, 3H), 0.94 (dd, *J* = 16.9, 6.8 Hz, 6H). $^{13}$C NMR (100 MHz, Chloroform-*d*): δ 174.8, 171.7, 156.6, 140.2, 136.1, 128.6, 128.3, 128.0, 127.0, 114.9, 68.0, 67.2, 60.3, 50.5, 40.0, 31.0, 25.6, 21.8, 19.1, 17.9.

**(5S,8S,11S)-methyl 5-isopropyl-8-(2-methylallyl)-3,6,9-trioxo-11-(((S)-2-oxopyrrolidin-3-yl)methyl)-1-phenyl-2-oxa-4,7,10-triazadodecan-12-oate (MPI12d). MPI12d** was prepared as a white solid following a similar procedure to **MPI11d** (yield 63%). $^1$H NMR (400 MHz, Chloroform-*d*) δ 7.82 (d, *J* = 7.7 Hz, 1H), 7.41 – 7.28 (m, 5H), 7.06 (d, *J* = 8.5 Hz, 1H), 6.67 (s, 1H), 5.45 (d, *J* = 8.8 Hz, 1H),

5.09 (s, 2H), 4.75 (s, 1H), 4.71 (s, 1H), 4.54 (ddd, *J* = 11.6, 7.8, 3.4 Hz, 1H), 4.05 – 3.95 (m, 1H), 3.70 (s, 3H), 3.31 (p, *J* = 8.1, 6.7 Hz, 2H), 2.58 (dd, *J* = 14.1, 4.8 Hz, 1H), 2.45 – 2.28 (m, 3H), 2.25 – 2.05 (m, 2H), 1.90 – 1.73 (m, 2H), 1.72 (s, 3H), 0.91 (dd, *J* = 18.8, 6.8 Hz, 6H). $^{13}$C NMR (100 MHz, Chloroform-*d*): δ 179.8, 172.0, 171.8, 171.2, 156.5, 140.9, 136.2, 128.6, 128.2, 128.0, 114.3, 67.1, 60.6, 52.4, 51.3, 41.1, 40.5, 38.3, 33.3, 30.9, 28.1, 21.9, 19.2, 18.6, 17.3.

**Benzyl ((S)-1-(((S)-1-(((S)-1-hydroxy-3-((S)-2-oxopyrrolidin-3-yl)propan-2-yl)amino)-4-methyl-1-oxopent-4-en-2-yl)amino)-3-methyl-1-oxobutan-2-yl)carbamate (MPI12e).** **MPI12e** was prepared as a white solid following a similar procedure to **MPI11e** (yield 70%). $^{1}$H NMR (400 MHz, Methanol-$d_4$) δ 7.36 – 7.15 (m, 5H), 5.01 (q, *J* = 12.5 Hz, 2H), 4.69 (d, *J* = 8.8 Hz, 2H), 4.41 (dd, *J* = 9.5, 5.6 Hz, 1H), 3.84 (dd, *J* = 16.3, 7.9 Hz, 2H), 3.44 (dd, *J* = 11.2, 5.4 Hz, 1H), 3.36 (dd, *J* = 11.0, 6.4 Hz, 1H), 3.20 – 3.12 (m, 2H), 2.50 – 2.16 (m, 4H), 2.00 – 1.80 (m, 2H), 1.65 (s, 4H), 1.45 (ddd, *J* = 14.2, 11.1, 3.1 Hz, 1H), 0.84 (dd, *J* = 11.7, 6.8 Hz, 6H). $^{13}$C NMR (100 MHz, Methanol-$d_4$) δ 181.2, 172.8, 172.6, 157.5, 141.0, 136.8, 128.1, 127.6, 127.5, 112.9, 66.5, 64.1, 61.0, 52.0, 49.3, 40.1, 39.4, 38.1, 32.2, 30.4, 27.6, 20.8, 18.3, 17.1.

**Benzyl ((S)-3-methyl-1-(((S)-4-methyl-1-oxo-1-(((S)-1-oxo-3-((S)-2-oxopyrrolidin-3-yl)propan-2-yl)amino)pent-4-en-2-yl)amino)-1-oxobutan-2-yl)carbamate (MPI12).** **MPI12** was prepared as a white solid following a similar procedure to **MPI11** (yield 60%). $^{1}$H NMR (400 MHz, Chloroform-*d*) δ 9.48 (s, 1H), 8.33 – 8.20 (m, 1H), 7.42 – 7.29 (m, 5H), 6.63 (d, *J* = 7.8 Hz, 1H), 6.05 (s, 1H), 5.34 (d, *J* = 8.2 Hz, 1H), 5.10 (s, 2H), 4.82 (t, *J* = 1.7 Hz, 1H), 4.75 (s, 1H), 4.68 (td, *J* = 8.5, 5.7 Hz, 1H), 4.33 (d, *J* = 8.5 Hz, 1H), 4.02 (dd, *J* = 8.2, 5.8 Hz, 1H), 3.33 (dt, *J* = 8.9, 4.3 Hz, 2H), 2.62 (dd, *J* = 14.1, 5.5 Hz, 1H), 2.53 – 2.32 (m, 3H), 2.21 – 2.09 (m, 1H), 2.05 – 1.88 (m, 2H), 1.87 – 1.82 (m, 1H), 1.75 (s, 3H), 0.93 (dd, *J* = 23.8, 6.8 Hz, 6H). $^{13}$C NMR (100 MHz, Chloroform-*d*): δ 199.7, 180.0, 172.4, 171.4, 156.6, 140.8, 136.2, 128.5, 128.2, 128.0, 114.4, 67.1, 60.6, 57.5, 51.4, 41.0, 40.6, 38.0, 30.9, 29.9, 28.3, 21.9, 19.2, 17.7.

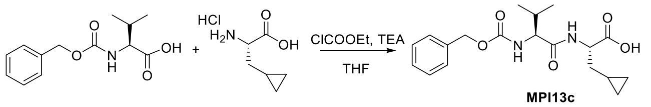


**(S)-2-((S)-2-(((benzyloxy)carbonyl)amino)-3-methylbutanamido)-3-cyclopropylpropanoic acid (MPI13c).** *N*-Cbz-L-valine (500 mg, 2.0 mmol, 1.0 equiv), ethyl chloroformate (268 μL, 2.8 mmol, 1.4 equiv) and TEA (836 μL, 6.0 mmol, 3.0 equiv) were dissolved in anhydrous THF (30 mL). After stirring at 0 °C for 0.5 h, L-cyclopropylalanine (386 mg, 4.0 mmol, 1.5 equiv) in water was added. The reaction mixture was stirred for 0.5 h at 0 °C. The THF was removed in vacuo and the aqueous layer was acidified with 1.0 M HCl. The aqueous layer was extracted with ethyl acetate. The combined organic layers were dried over anhydrous $Na_2SO_4$, filtered and concentrated. The residue was recrystallized from $CHCl_3$ and hexane to yield the pure product **MPI13c** as a white solid (380 mg, yield 52%).

**Methyl (5S,8S,11S)-8-(cyclopropylmethyl)-5-isopropyl-3,6,9-trioxo-11-(((S)-2-oxopyrrolidin-3-yl)methyl)-1-phenyl-2-oxa-4,7,10-triazadodecan-12-oate (MPI13d). MPI13d** was prepared as a white solid following a similar procedure to **MPI11d** (yield 51%).

**Benzyl ((S)-1-(((S)-3-cyclopropyl-1-(((S)-1-hydroxy-3-((S)-2-oxopyrrolidin-3-yl)propan-2-yl)amino)-1-oxopropan-2-yl)amino)-3-methyl-1-oxobutan-2-yl)carbamate (MPI13e). MPI13e** was prepared as a white solid following a similar procedure to **MPI11e** (yield 75%). $^1$H NMR (400 MHz, Methanol-$d_4$) δ 7.3 – 7.1 (m, 5H), 5.0 (d, $J$ = 4.1 Hz, 2H), 4.3 (t, $J$ = 7.3 Hz, 1H), 3.9 (d, $J$ = 7.3 Hz, 2H), 3.5 – 3.3 (m, 2H), 3.1 (dt, $J$ = 16.3, 5.5 Hz, 2H), 2.4 (dd, $J$ = 14.7, 6.6 Hz, 1H), 2.2 (dt, $J$ = 15.7, 7.8 Hz, 1H), 2.0 – 1.8 (m, 2H), 1.6 (dq, $J$ = 20.5, 7.7, 7.0 Hz, 2H), 1.5 – 1.3 (m, 2H), 0.8 (dd, $J$ = 9.4, 6.7 Hz, 6H), 0.7 (p, $J$ = 6.9 Hz, 1H), 0.3 (t, $J$ = 10.5 Hz, 2H), 0.0 (dq, $J$ = 7.5, 4.0 Hz, 2H). $^{13}$C NMR (100 MHz, Methanol-$d_4$) δ 181.3, 172.9, 172.7, 157.4, 136.8, 128.1, 127.6, 127.5, 66.4, 64.1, 60.7, 54.3, 49.1, 40.1, 38.1, 36.6, 32.2, 30.6, 27.6, 18.4, 17.3, 7.3, 3.8, 3.6.

**Benzyl ((S)-1-(((S)-3-cyclopropyl-1-oxo-1-(((S)-1-oxo-3-((S)-2-oxopyrrolidin-3-yl)propan-2-yl)amino)propan-2-yl)amino)-3-methyl-1-oxobutan-2-yl)carbamate (MPI13). MPI13** was prepared as a white solid following a similar procedure to **MPI11** (yield 50%). $^1$H NMR (400 MHz, Chloroform-$d$) δ 9.4 (s, 1H), 8.3 (d, $J$ = 6.5 Hz, 1H), 7.2 (s, 5H), 7.2 (s, 1H), 6.6 (d, $J$ = 23.9 Hz, 1H), 5.7 – 5.5 (m, 1H), 5.1 – 4.9 (m, 2H), 4.6 (dq, $J$ = 18.1, 9.9, 8.6 Hz, 1H), 4.4 – 4.2 (m, 1H), 4.0 (h, $J$ = 6.9, 5.7 Hz, 1H), 3.3 (s, 1H), 3.2 (d, $J$ = 10.8 Hz, 1H), 2.4 (t, $J$ = 8.4 Hz, 1H), 2.3 (s, 1H), 2.1 (s, 1H), 2.0 (dp, $J$ = 25.6, 9.2, 7.8 Hz, 2H), 1.9 – 1.6 (m, 2H), 1.5 – 1.4 (m, 1H), 0.8 (dd, $J$ = 12.2, 6.6 Hz, 6H), 0.7 – 0.5 (m, 1H), 0.3 (t, $J$ = 9.5 Hz, 2H). $^{13}$C NMR (100 MHz, Chloroform-$d$) δ 199.5, 180.0, 172.7, 171.3, 156.6, 136.2, 128.5, 128.2, 128.0, 67.1, 60.5, 53.7, 50.7, 40.6, 38.0, 37.6, 31.1, 29.8, 28.4, 19.2, 17.9, 7.3, 4.5, 4.4.

**Methyl (S)-2-((S)-2-(((benzyloxy)carbonyl)amino)-3-methylbutanamido)-3-(furan-2-yl)propanoate (MPI14b). MPI14b** was prepared with methyl (S)-2-amino-3-(furan-2-yl)propanoate hydrochloride and *N*-Cbz-L-valine as a white solid following a similar procedure to **MPI11b** (yield 80%). $^1$H NMR (400 MHz, Methanol-$d_4$) δ 7.32 – 7.09 (m, 6H), 6.15 (dd, $J$ = 3.2, 1.9 Hz, 1H), 6.02 (d, $J$ = 3.2 Hz, 1H), 4.99 (s, 2H), 4.59 (dd, $J$ = 8.1, 5.5 Hz, 1H), 3.84 (d, $J$ = 7.1 Hz, 1H), 3.59 (s, 3H), 3.14 – 2.90 (m, 2H), 2.02 – 1.77 (m, 1H), 0.82 (dd, $J$ = 11.5, 6.8 Hz, 6H). $^{13}$C NMR (100 MHz, Methanol-$d_4$) δ 172.73, 171.40, 150.58, 141.76, 128.09, 127.46, 109.92, 107.20, 48.25, 48.04, 47.82, 47.61, 47.40, 47.19, 46.97, 30.61, 29.49, 18.25.

**(S)-2-((S)-2-(((benzyloxy)carbonyl)amino)-3-methylbutanamido)-3-(furan-2-yl)propanoic acid (MPI14c). MPI14c** was prepared as a white solid following a similar procedure to **MPI11c** (yield 88%). $^1$H NMR (400 MHz, Methanol-$d_4$) δ 7.30 – 7.14 (m, 6H), 6.19 – 6.08 (m, 1H), 6.02 (d, $J$ = 3.2 Hz, 1H), 4.98 (s, 2H), 4.58 (dd, $J$ = 8.2, 5.0 Hz, 1H), 3.92 – 3.76 (m, 1H), 3.11 (dd, $J$ = 15.3, 5.1 Hz, 1H), 2.97 (dd, $J$ = 15.3, 8.2 Hz, 1H), 1.99 – 1.82 (m, 1H), 0.82 (dd, $J$ = 14.3, 6.8 Hz, 6H). $^{13}$C NMR (100 MHz, Methanol-$d_4$) δ 172.61, 172.49, 150.87, 141.63, 128.10, 127.63, 127.45, 109.89, 107.09, 66.31, 60.59, 48.27, 48.06, 47.84, 47.63, 47.42, 47.21, 46.99, 30.63, 18.32.

**Methyl (5S,8S,11S)-8-(furan-2-ylmethyl)-5-isopropyl-3,6,9-trioxo-11-(((S)-2-oxopyrrolidin-3-yl)methyl)-1-phenyl-2-oxa-4,7,10-triazadodecan-12-oate (MPI14d).** **MPI14d** was prepared as a white solid following a similar procedure to **MPI11d** (yield 80%). $^1$H NMR (400 MHz, Chloroform-*d*) δ 7.67 (d, *J* = 7.2 Hz, 1H), 7.38 – 7.22 (m, 5H), 6.90 (d, *J* = 7.9 Hz, 1H), 6.23 – 6.11 (m, 1H), 6.03 (d, *J* = 3.2 Hz, 1H), 5.86 (s, 1H), 5.26 (d, *J* = 8.0 Hz, 1H), 5.04 (d, *J* = 5.1 Hz, 2H), 4.74 (d, *J* = 7.3 Hz, 1H), 4.45 (s, 1H), 3.95 (t, *J* = 7.1 Hz, 1H), 3.64 (s, 3H), 3.24 (d, *J* = 8.8 Hz, 1H), 3.17 – 3.01 (m, 2H), 2.31 (s, 1H), 2.19 (s, 1H), 2.07 (dd, *J* = 13.3, 6.9 Hz, 2H), 1.86 – 1.68 (m, 2H), 1.60 (s, 3H), 1.41 (d, *J* = 22.6 Hz, 4H), 0.96 – 0.74 (m, 6H).

**Benzyl ((S)-1-(((S)-3-(furan-2-yl)-1-(((S)-1-hydroxy-3-((S)-2-oxopyrrolidin-3-yl) propan-2-yl)amino)-1-oxopropan-2-yl)amino)-3-methyl-1-oxobutan-2-yl)carbamate (MPI14e).** **MPI14e** was prepared as a white solid following a similar procedure to **MPI11e** (yield 37%). $^1$H NMR (400 MHz, Methanol-$d_4$) δ 7.25 (s, 6H), 6.24 – 6.14 (m, 1H), 6.05 (d, *J* = 3.1 Hz, 1H), 5.09 – 4.86 (m, 2H), 4.52 (d, *J* = 7.9 Hz, 2H), 3.80 (t, *J* = 9.6 Hz, 2H), 3.48 – 3.25 (m, 2H), 3.11 – 2.88 (m, 2H), 2.39 – 2.02 (m, 2H), 2.03 – 1.73 (m, 2H), 1.73 – 1.55 (m, 1H), 1.51 – 1.32 (m, 1H), 1.26 (dd, *J* = 6.8, 2.3 Hz, 1H), 0.80 (dd, *J* = 9.1, 6.5 Hz, 6H). $^{13}$C NMR (100 MHz, Methanol-$d_4$) δ 181.23, 172.74, 171.62, 157.61, 151.02, 141.73, 136.74, 128.10, 127.66, 127.57, 109.96, 107.24, 66.57, 64.01, 61.10, 52.89, 48.26, 48.05, 47.83, 47.62, 47.41, 47.20, 46.98, 40.11, 38.05, 32.13, 29.99 (d, *J* = 65.0 Hz), 27.59, 18.23, 17.08.

**Benzyl ((S)-1-(((S)-3-(furan-2-yl)-1-oxo-1-(((S)-1-oxo-3-((S)-2-oxopyrrolidin-3-yl)propan-2-yl)amino)propan-2-yl)amino)-3-methyl-1-oxobutan-2-yl)carbamate (MPI14).** **MPI14** was prepared as a white solid following a similar procedure to **MPI11** (yield 80%). $^1$H NMR (400 MHz, Chloroform-*d*) δ 9.34 (s, 1H), 8.22 (d, *J* = 6.8 Hz, 1H), 7.27 (q, *J* = 7.8, 5.8 Hz, 5H), 7.02 (d, *J* = 8.1 Hz, 1H), 6.16 (d, *J* = 2.5 Hz, 2H), 6.03 (d, *J* = 3.1 Hz, 1H), 5.40 (t, *J* = 11.0 Hz, 1H), 5.03 (d, *J* = 6.8 Hz, 2H), 4.79 (q, *J* = 6.7 Hz, 1H), 4.22 (d, *J* = 10.4 Hz, 1H), 4.00 – 3.88 (m, 1H), 2.25 (s, 2H), 2.17 – 2.02 (m, 1H), 1.76 (d, *J* = 35.2 Hz, 5H), 0.84 (dd, *J* = 25.3, 6.8 Hz, 6H). $^{13}$C NMR (100 MHz, Chloroform-*d*) δ 199.96, 179.97, 171.32, 171.18, 150.71, 142.04, 136.11, 128.60, 128.29, 128.11, 110.41, 108.07, 67.26, 52.41, 50.86, 40.56, 30.70, 28.57, 19.26, 17.59.

**Methyl (S)-2-((S)-2-(((benzyloxy)carbonyl)amino)-3-methylbutanamido)-3-(thiophen-2-yl)propanoate (MPI15b).** **MPI15b** was prepared with methyl (S)-2-amino-3-(thiophen-2-yl)propanoate and *N*-Cbz-L-valine as a white solid following a similar procedure to **MPI11b** (yield 72%). $^1$H NMR (400 MHz, Methanol-$d_4$) δ 7.37 – 7.13 (m, 5H), 7.11 – 7.03 (m, 1H), 6.78 (dd, *J* = 4.6, 2.9 Hz, 2H), 4.99 (d, *J* = 2.9 Hz, 2H), 4.58 (dd, *J* = 7.9, 5.3 Hz, 1H), 3.90 – 3.74 (m, 1H), 3.60 (s, 3H), 3.27 (dd, *J* = 14.9, 5.3 Hz, 1H), 3.21 – 3.12 (m, 3H), 1.91 (s, 1H), 0.82 (dd, *J* = 8.4, 6.7 Hz, 6H). $^{13}$C NMR (100 MHz, Methanol-$d_4$) δ 172.79, 171.25, 138.12, 136.86, 128.10, 127.63, 127.47, 66.30, 60.62, 53.75, 51.40, 48.26, 48.04, 47.83, 47.62, 47.41, 47.19, 46.98, 31.05, 30.62, 18.28, 17.13.

**(S)-2-((S)-2-(((benzyloxy)carbonyl)amino)-3-methylbutanamido)-3-(thiophen-2-yl)propanoic acid (MPI15c).** **MPI15c** was prepared as a white solid following a similar procedure to **MPI11c** (yield 60%). $^1$H NMR (400 MHz, Methanol-$d_4$) δ 7.44 –

7.29 (m, 5H), 7.19 (dd, $J$ = 4.9, 1.5 Hz, 1H), 6.97 – 6.83 (m, 2H), 5.12 (d, $J$ = 2.6 Hz, 2H), 4.69 (dd, $J$ = 7.9, 4.9 Hz, 1H), 3.99 (d, $J$ = 7.1 Hz, 1H), 3.43 (dd, $J$ = 15.0, 4.8 Hz, 1H), 3.31 – 3.22 (m, 1H), 2.14 – 1.93 (m, 1H), 1.08 – 0.86 (m, 6H).

**Methyl (5S,8S,11S)-5-isopropyl-3,6,9-trioxo-11-(((S)-2-oxopyrrolidin-3-yl)methyl)-1-phenyl-8-(thiophen-2-ylmethyl)-2-oxa-4,7,10-triazadodecan-12-oate (MPI15d). MPI15d** was prepared as a white solid following a similar procedure to **MPI11d** (yield 90%). $^1$H NMR (400 MHz, Methanol-$d_4$) δ 7.35 – 7.18 (m, 5H), 7.08 (dd, $J$ = 5.0, 1.3 Hz, 1H), 6.84 – 6.70 (m, 1H), 5.03 – 4.95 (m, 2H), 4.58 – 4.48 (m, 1H), 4.44 – 4.32 (m, 1H), 3.88 – 3.73 (m, 1H), 3.60 (d, $J$ = 6.3 Hz, 3H), 3.14 (d, $J$ = 3.4 Hz, 2H), 2.39 – 2.25 (m, 1H), 2.24 – 2.09 (m, 1H), 2.10 – 2.01 (m, 1H), 1.99 – 1.83 (m, 1H), 1.67 (s, 2H), 1.19 (d, $J$ = 3.6 Hz, 2H), 0.89 (dd, $J$ = 9.1, 6.8 Hz, 2H), 0.78 (dd, $J$ = 8.4, 6.7 Hz, 4H). $^{13}$C NMR (100 MHz, Methanol-$d_4$) δ 180.21, 172.65, 172.01, 171.69, 157.27, 138.44, 136.84, 128.14, 127.66, 127.51, 126.54, 126.36, 124.01, 66.40, 51.58, 50.54, 48.32, 48.11, 47.90, 47.69, 47.47, 47.26, 47.05, 32.51, 31.41, 27.30, 18.40, 17.21.

**Benzyl ((S)-1-(((S)-1-(((S)-1-hydroxy-3-((S)-2-oxopyrrolidin-3-yl)propan-2-yl)amino)-1-oxo-3-(thiophen-2-yl)propan-2-yl)amino)-3-methyl-1-oxobutan-2-yl)carbamate (MPI15e). MPI15e** was prepared as a white solid following a similar procedure to **MPI11e** (yield 53%). $^1$H NMR (400 MHz, Methanol-$d_4$) δ 7.32 – 7.15 (m, 5H), 7.10 (d, $J$ = 4.8 Hz, 1H), 6.80 (d, $J$ = 4.9 Hz, 2H), 5.08 – 4.88 (m, 2H), 4.48 (s, 1H), 3.80 (d, $J$ = 6.7 Hz, 2H), 3.30 (d, $J$ = 23.7 Hz, 2H), 3.17 – 3.00 (m, 3H), 2.24 (dd, $J$ = 24.9, 14.9 Hz, 2H), 2.02 – 1.69 (m, 2H), 1.70 – 1.54 (m, 1H), 1.50 – 1.24 (m, 1H), 1.19 (s, 1H), 0.78 (t, $J$ = 7.2 Hz, 6H).

**Benzyl ((S)-3-methyl-1-oxo-1-(((S)-1-oxo-1-(((S)-1-oxo-3-((S)-2-oxopyrrolidin-3-yl)propan-2-yl)amino)-3-(thiophen-2-yl)propan-2-yl)amino)butan-2-yl)carbamate (MPI15). MPI15** was prepared as a white solid following a similar procedure to **MPI11** (yield 60%). $^1$H NMR (400 MHz, Chloroform-$d$) δ 9.30 (s, 1H), 8.24 (s, 1H), 7.38 – 7.22 (m, 4H), 7.04 (d, $J$ = 5.0 Hz, 1H), 6.91 (d, $J$ = 8.4 Hz, 1H), 6.87 – 6.65 (m, 2H), 5.97 (s, 1H), 5.33 (d, $J$ = 8.2 Hz, 1H), 5.03 (s, 2H), 4.80 (d, $J$ = 7.7 Hz, 1H), 4.20 (s, 1H), 3.97 (t, $J$ = 7.2 Hz, 1H), 3.49 – 3.03 (m, 4H), 2.24 (s, 2H), 2.14 – 1.93 (m, 2H), 1.76 (d, $J$ = 33.7 Hz, 3H), 0.83 (dd, $J$ = 25.9, 6.7 Hz, 6H).

**Methyl (S)-2-((2S,3R)-2-(((benzyloxy)carbonyl)amino)-3-(tert-butoxy)butanamido)-4,4-dimethylpentanoate (MPI16b). MPI16b** was prepared with Z-Thr(tBu)-OH and methyl (S)-2-amino-4,4-dimethylpentanoate hydrochloride as a white solid following a similar procedure to **MPI11b** (yield 83%).$^1$H NMR (400 MHz, Chloroform-$d$) δ 7.5 (d, $J$ = 8.1 Hz, 1H), 7.3 (d, $J$ = 4.2 Hz, 5H), 5.9 (d, $J$ = 4.3 Hz, 1H), 5.1 – 4.9 (m, 2H), 4.5 (td, $J$ = 9.0, 3.2 Hz, 1H), 4.1 (d, $J$ = 5.0 Hz, 2H), 3.6 (s, 3H), 1.7 (dd, $J$ = 14.5, 3.2 Hz, 1H), 1.4 (dd, $J$ = 14.5, 9.4 Hz, 1H), 1.1 (d, $J$ = 74.2 Hz, 12H), 0.9 (s, 9H). $^{13}$C NMR (100 MHz, Chloroform-$d$) δ 173.2, 169.2, 156.1, 136.3, 128.5, 128.1, 127.9, 75.6, 66.9, 66.8, 58.4, 52.2, 50.3, 45.7, 30.7, 29.7, 28.2, 16.2.

**(S)-2-((2S,3R)-2-(((benzyloxy)carbonyl)amino)-3-(tert-butoxy)butanamido)-4,4-dimethylpentanoic acid (MPI16c). MPI16c** was prepared as a white solid following a similar procedure to **MPI11c** (yield 82%).

**Methyl (5S,8S,11S)-5-((R)-1-(tert-butoxy)ethyl)-8-(cyclopropylmethyl)-3,6,9-trioxo-11-(((S)-2-oxopyrrolidin-3-yl)methyl)-1-phenyl-2-oxa-4,7,10-triazadodecan-12-oate (MPI16d).** **MPI16d** was prepared as a white solid following a similar procedure to **MPI11d** (yield 66%). $^1$H NMR (400 MHz, Chloroform-*d*) δ 7.9 (d, *J* = 7.5 Hz, 1H), 7.5 (d, *J* = 8.4 Hz, 1H), 7.2 (q, *J* = 8.0, 6.1 Hz, 5H), 7.2 (s, 1H), 6.0 (d, *J* = 5.5 Hz, 1H), 5.0 (q, *J* = 12.3 Hz, 2H), 4.5 (td, *J* = 8.4, 4.1 Hz, 1H), 4.4 (ddd, *J* = 11.9, 7.4, 3.5 Hz, 1H), 4.1 (p, *J* = 4.2 Hz, 2H), 3.6 (s, 3H), 3.2 (dd, *J* = 20.8, 9.1 Hz, 2H), 2.3 (qd, *J* = 9.9, 3.9 Hz, 1H), 2.2 – 2.1 (m, 2H), 1.8 – 1.6 (m, 3H), 1.4 (dd, *J* = 14.5, 8.6 Hz, 1H), 1.3 – 0.9 (m, 12H), 0.9 (s, 9H). $^{13}$C NMR (100 MHz, Chloroform-*d*) δ 179.6, 172.5, 172.0, 168.9, 155.9, 136.0, 128.3, 128.0, 127.8, 75.3, 66.7, 66.7, 58.6, 52.0, 50.9, 50.7, 45.8, 40.3, 38.0, 32.8, 30.2, 29.6, 28.0, 27.6, 16.8.

**Benzyl ((2S,3R)-3-(tert-butoxy)-1-(((S)-1-(((S)-1-hydroxy-3-((S)-2-oxopyrrolidin-3-yl)propan-2-yl)amino)-4,4-dimethyl-1-oxopentan-2-yl)amino)-1-oxobutan-2-yl)carbamate (MPI16e).** **MPI16e** was prepared as a white solid following a similar procedure to **MPI11e** (yield 66%). $^1$H NMR (400 MHz, Chloroform-*d*) δ 8.1 (d, *J* = 8.7 Hz, 1H), 7.5 (d, *J* = 8.0 Hz, 1H), 7.3 – 7.1 (m, 6H), 6.8 (s, 1H), 5.1 – 4.9 (m, 2H), 4.4 (q, *J* = 6.9 Hz, 2H), 4.1 (dd, *J* = 6.3, 3.8 Hz, 1H), 4.0 – 3.9 (m, 1H), 3.9 (q, *J* = 10.1, 9.2 Hz, 1H), 3.6 – 3.4 (m, 2H), 3.1 (dtd, *J* = 17.2, 9.3, 8.7, 5.0 Hz, 2H), 2.4 – 2.1 (m, 3H), 1.8 – 1.6 (m, 2H), 1.3 (dd, *J* = 14.3, 6.9 Hz, 2H), 1.2 (s, 9H), 0.9 (d, *J* = 6.3 Hz, 3H), 0.8 (s, 9H). $^{13}$C NMR (100 MHz, Chloroform-*d*) δ 181.7, 172.7, 169.0, 156.3, 136.3, 128.4, 128.0, 127.9, 75.6, 67.2, 66.7, 65.2, 58.6, 53.4, 51.2, 49.1, 45.4, 40.5, 38.5, 33.9, 30.2, 29.5, 28.1, 16.5.

**Benzyl ((2S,3R)-3-(tert-butoxy)-1-(((S)-4,4-dimethyl-1-oxo-1-(((S)-1-oxo-3-((S)-2-oxopyrrolidin-3-yl)propan-2-yl)amino)pentan-2-yl)amino)-1-oxobutan-2-yl)carbamate (MPI16).** **MPI16** was prepared as a white solid following a similar procedure to **MPI11** (yield 40%). $^1$H NMR (400 MHz, Chloroform-*d*) δ 9.4 (s, 1H), 8.1 (d, *J* = 6.6 Hz, 1H), 7.5 (d, *J* = 8.3 Hz, 1H), 7.3 – 7.2 (m, 5H), 6.8 (s, 1H), 5.9 (d, *J* = 5.1 Hz, 1H), 5.0 (q, *J* = 12.2 Hz, 2H), 4.5 (td, *J* = 8.5, 4.1 Hz, 1H), 4.3 (dq, *J* = 10.8, 5.4, 4.8 Hz, 1H), 4.1 (d, *J* = 5.8 Hz, 2H), 3.3 – 3.1 (m, 2H), 2.3 (p, *J* = 8.3 Hz, 1H), 2.3 – 2.2 (m, 1H), 1.9 (td, *J* = 12.4, 10.2, 5.7 Hz, 1H), 1.8 (ddt, *J* = 11.8, 8.3, 3.9 Hz, 2H), 1.7 (tt, *J* = 12.3, 6.2 Hz, 1H), 1.4 (dd, *J* = 14.4, 8.6 Hz, 1H), 1.1 (d, *J* = 81.8 Hz, 12H), 0.9 (d, *J* = 7.3 Hz, 9H). $^{13}$C NMR (100 MHz, Chloroform-*d*) δ 199.6, 180.0, 173.3, 169.3, 156.2, 136.2, 128.6, 128.2, 128.1, 75.6, 67.0, 66.8, 58.9, 57.3, 51.2, 46.1, 40.5, 37.8, 30.5, 29.9, 29.8, 29.7, 28.2, 17.0.

**Methyl (S)-2-((2S,3R)-2-(((benzyloxy)carbonyl)amino)-3-(tert-butoxy)butanamido)-3-cyclopropylpropanoate (MPI17b).** **MPI17b** was prepared with Z-Thr(tBu)-OH and methyl (S)-2-amino-3-cyclopropylpropanoate hydrochloride as a white solid following a similar procedure to **MPI11b** (yield 79%). $^1$H NMR (400 MHz, Chloroform-*d*) δ 7.7 (d, *J* = 7.5 Hz, 1H), 7.3 (d, *J* = 4.3 Hz, 5H), 5.9 (d, *J* = 5.3 Hz, 1H), 5.1 – 5.0 (m, 2H), 4.6 (q, *J* = 6.3 Hz, 1H), 4.2 – 4.1 (m, 2H), 3.7 (s, 3H), 1.6 (ddt, *J* = 27.6, 13.9, 7.1 Hz, 2H), 1.2 (s, 8H), 1.0 (d, *J* = 6.4 Hz, 4H), 0.6 (p, *J* = 6.3 Hz, 1H), 0.4 (d, *J* = 8.1 Hz, 2H), -0.0 (q, *J* = 8.7, 7.3 Hz, 2H). $^{13}$C NMR (100 MHz,

Chloroform-*d*) δ 172.2, 169.0, 156.0, 136.2, 128.4, 128.0, 127.9, 75.4, 66.8, 66.7, 58.5, 52.7, 52.0, 36.9, 28.1, 16.5, 6.9, 4.2, 4.0.

**(S)-2-((2S,3R)-2-(((benzyloxy)carbonyl)amino)-3-(tert-butoxy)butanamido)-3-cyclopropylpropanoic acid (MPI17c).** **MPI17c** was prepared as a white solid following a similar procedure to **MPI11c** (yield 90%).

**Methyl (5S,8S,11S)-5-((R)-1-(tert-butoxy)ethyl)-8-(cyclopropylmethyl)-3,6,9-trioxo-11-(((S)-2-oxopyrrolidin-3-yl)methyl)-1-phenyl-2-oxa-4,7,10-triazadodecan-12-oate (MPI17d).** **MPI17d** was prepared as a white solid following a similar procedure to **MPI11d** (yield 60%). $^{1}$H NMR (400 MHz, Chloroform-*d*) δ 7.8 (d, *J* = 7.4 Hz, 1H), 7.6 (d, *J* = 7.9 Hz, 1H), 7.2 (d, *J* = 12.4 Hz, 5H), 7.0 (s, 1H), 5.9 (d, *J* = 6.0 Hz, 1H), 5.0 (t, *J* = 10.8 Hz, 2H), 4.5 (q, *J* = 7.1 Hz, 1H), 4.4 (ddd, *J* = 11.4, 7.7, 3.8 Hz, 1H), 4.1 – 4.0 (m, 2H), 3.6 (s, 3H), 3.2 (dt, *J* = 16.5, 9.1 Hz, 2H), 2.3 (ddq, *J* = 28.6, 13.9, 7.6 Hz, 2H), 2.1 (ddd, *J* = 15.6, 11.7, 4.2 Hz, 1H), 1.8 – 1.6 (m, 2H), 1.6 (q, *J* = 7.7 Hz, 2H), 1.2 (s, 8H), 1.0 (d, *J* = 6.2 Hz, 4H), 0.7 (td, *J* = 8.0, 4.2 Hz, 1H), 0.4 (dd, *J* = 8.1, 5.0 Hz, 2H), 0.0 (d, *J* = 4.9 Hz, 2H). $^{13}$C NMR (100 MHz, Chloroform-*d*) δ 179.8, 172.1, 171.7, 169.2, 156.1, 136.2, 128.5, 128.1, 128.0, 75.2, 66.8, 58.9, 53.6, 53.5, 52.2, 50.9, 40.4, 38.2, 37.6, 33.0, 28.2, 27.9, 17.3, 7.1, 4.4, 4.2.

**Benzyl ((2S,3R)-3-(tert-butoxy)-1-(((S)-3-cyclopropyl-1-(((S)-1-hydroxy-3-((S)-2-oxopyrrolidin-3-yl)propan-2-yl)amino)-1-oxopropan-2-yl)amino)-1-oxobutan-2-yl)carbamate (MPI17e).** **MPI17e** was prepared as a white solid following a similar procedure to **MPI11e** (yield 85%). $^{1}$H NMR (400 MHz, Chloroform-*d*) δ 7.6 (t, *J* = 7.0 Hz, 2H), 7.3 (d, *J* = 4.3 Hz, 5H), 7.0 (s, 1H), 6.1 (d, *J* = 6.3 Hz, 1H), 5.1 – 4.9 (m, 2H), 4.4 (t, *J* = 7.2 Hz, 1H), 4.2 (d, *J* = 6.6 Hz, 1H), 4.2 – 4.0 (m, 2H), 4.0 (ddt, *J* = 13.4, 9.3, 4.5 Hz, 1H), 3.6 – 3.4 (m, 2H), 3.2 (s, 2H), 2.4 – 2.2 (m, 2H), 2.0 (d, *J* = 12.4 Hz, 1H), 1.8 – 1.4 (m, 4H), 1.1 (d, *J* = 66.6 Hz, 12H), 0.6 (t, *J* = 7.0 Hz, 1H), 0.4 (q, *J* = 8.3, 7.4 Hz, 2H), 0.0 (s, 2H). $^{13}$C NMR (100 MHz, Chloroform-*d*) δ 181.0, 171.8, 169.4, 156.2, 136.1, 128.4, 128.1, 128.0, 75.1, 67.8, 66.8, 65.1, 59.1, 54.0, 49.7, 40.5, 38.2, 37.3, 32.4, 28.1, 28.0, 17.6, 7.2, 4.3, 4.3.

**Benzyl ((2S,3R)-3-(tert-butoxy)-1-(((S)-3-cyclopropyl-1-oxo-1-(((S)-1-oxo-3-((S)-2-oxopyrrolidin-3-yl)propan-2-yl)amino)propan-2-yl)amino)-1-oxobutan-2-yl)carbamate (MPI17).** **MPI17** was prepared as a white solid following a similar procedure to **MPI11** (yield 54%). $^{1}$H NMR (400 MHz, Chloroform-*d*) δ 9.4 (s, 1H), 8.1 (d, *J* = 6.3 Hz, 1H), 7.5 (t, *J* = 8.6 Hz, 1H), 7.3 (s, 5H), 6.4 – 6.1 (m, 1H), 5.9 (d, *J* = 5.4 Hz, 1H), 5.0 (q, *J* = 12.3 Hz, 2H), 4.4 (dt, *J* = 34.9, 6.6 Hz, 1H), 4.3 (p, *J* = 5.6, 4.6 Hz, 1H), 4.1 (t, *J* = 5.9 Hz, 2H), 3.3 (d, *J* = 8.6 Hz, 1H), 3.2 – 3.2 (m, 1H), 2.3 (d, *J* = 14.0 Hz, 1H), 2.2 (s, 2H), 2.0 – 1.9 (m, 1H), 1.9 (t, *J* = 6.8 Hz, 1H), 1.7 (dt, *J* = 15.4, 8.5 Hz, 1H), 1.6 (dd, *J* = 12.9, 6.3 Hz, 1H), 1.1 (d, *J* = 72.7 Hz, 12H), 0.7 (d, *J* = 9.1 Hz, 1H), 0.4 (q, *J* = 8.5 Hz, 2H), 0.0 (d, *J* = 5.1 Hz, 2H). $^{13}$C NMR (100 MHz, Chloroform-*d*) δ 199.7, 180.1, 172.4, 169.6, 156.3, 136.2, 128.7, 128.4, 128.2, 75.5, 67.1, 66.9, 59.2, 57.7, 55.0, 40.6, 38.0, 37.5, 29.9, 28.6, 28.3, 17.7, 7.4, 4.7, 4.5.

**(S)-methyl 2-((S)-2-(((benzyloxy)carbonyl)amino)-3,3-dimethylbutanamido)-4,4-dimethylpentanoate (MPI18b).** **MPI18b** was prepared with Cbz-L-tert-leucine and methyl (S)-2-amino-4,4-dimethylpentanoate hydrochloride as a white solid following a similar procedure to **MPI11b** (yield 81%). $^1$H NMR (400 MHz, Chloroform-*d*) δ 7.40 – 7.19 (m, 5H), 6.51 – 6.17 (m, 1H), 5.64 – 5.46 (m, 1H), 5.09 (s, 2H), 4.74 – 4.54 (m, 1H), 4.06 – 3.96 (m, 1H), 3.70 (d, *J* = 2.5 Hz, 3H), 1.83 – 1.74 (m, 1H), 1.52 – 1.38 (m, 1H), 1.00 (d, *J* = 2.4 Hz, 9H), 0.91 (d, *J* = 2.6 Hz, 9H). $^{13}$C NMR (100 MHz, Chloroform-*d*): δ 173.5, 170.2, 156.4, 136.3, 128.5, 128.1, 127.92, 127.90, 66.9, 60.4, 52.3, 49.7, 46.1, 34.8, 30.7, 29.4, 26.4, 21.1.

**(S)-2-((S)-2-(((benzyloxy)carbonyl)amino)-3,3-dimethylbutanamido)-4,4-dimethylpentanoic acid (MPI18c). MPI18c** was prepared as a white solid following a similar procedure to **MPI11c** (yield 79%). $^1$H NMR (400 MHz, Chloroform-*d*): δ 7.40 – 7.27 (m, 5H), 7.03 – 6.56 (m, 1H), 5.85 (s, 1H), 5.08 (q, *J* = 12.5 Hz, 2H), 4.64 – 4.42 (m, 1H), 4.14 – 3.94 (m, 1H), 1.85 – 1.72 (m, 1H), 1.59 – 1.43 (m, 1H), 0.94 (s, 9H), 0.89 (s, 9H).

**(5S,8S,11S)-methyl 5-(tert-butyl)-8-neopentyl-3,6,9-trioxo-11-(((S)-2-oxopyrrolidin-3-yl)methyl)-1-phenyl-2-oxa-4,7,10-triazadodecan-12-oate (MPI18d). MPI18d** was prepared as a white solid following a similar procedure to **MPI11d** (yield 60%). $^1$H NMR (400 MHz, Chloroform-*d*): δ 7.73 (d, *J* = 8.2 Hz, 1H), 7.60 (d, *J* = 9.0 Hz, 1H), 7.38 – 7.27 (m, 5H), 7.18 – 7.08 (m, 1H), 5.62 (d, *J* = 9.8 Hz, 1H), 5.06 (d, *J* = 2.4 Hz, 2H), 4.75 – 4.61 (m, 1H), 4.61 – 4.50 (m, 1H), 4.00 (dd, *J* = 9.6, 5.3 Hz, 1H), 3.67 (s, 3H), 3.41 – 3.21 (m, 2H), 2.51 – 2.32 (m, 2H), 2.29 – 2.16 (m, 1H), 1.94 – 1.74 (m, 3H), 1.49 – 1.44 (m, 1H), 0.96 (s, 9H), 0.89 (s, 9H). $^{13}$C NMR (100 MHz, Chloroform-*d*): δ 179.9, 173.0, 171.9, 170.2, 156.6, 136.3, 128.5, 128.5, 128.2, 128.1, 127.9, 67.0, 62.7, 55.5, 52.3, 50.8, 50.5, 46.8, 40.6, 38.2, 34.6, 30.6, 29.5, 27.8, 26.5, 26.4.

**Benzyl ((S)-1-(((S)-1-(((S)-1-hydroxy-3-((S)-2-oxopyrrolidin-3-yl)propan-2-yl)amino)-4,4-dimethyl-1-oxopentan-2-yl)amino)-3,3-dimethyl-1-oxobutan-2-yl)carbamate (MPI18e). MPI18e** was prepared as a white solid following a similar procedure to **MPI11e** (yield 67%). $^1$H NMR (400 MHz, Chloroform-*d*): δ 8.73 – 8.45 (m, 1H), 8.15 – 7.94 (m, 1H), 7.62 – 7.43 (m, 1H), 7.42 – 7.25 (H), 5.83 – 5.63 (m, 1H), 5.07 (d, *J* = 4.2 Hz, 2H), 4.93 – 4.75 (m, 1H), 4.49 – 4.33 (m, 1H), 4.01 – 3.85 (m, 1H), 3.85 – 3.67 (m, 1H), 3.51 – 3.22 (m, 3H), 3.22 – 3.03 (m, 1H), 2.35 – 2.23 (m, 1H), 22.20 – 2.10 (m, 1H), 1.90 – 1.55 (m, 4H), 1.55 – 1.37 (m, 1H), 0.93 (s, 9H), 0.90 (s, 9H).$^{13}$C NMR (100 MHz, Chloroform-*d*): δ 180.3, 173.5, 169.8, 156.5, 136.2, 128.6, 128.5, 128.2, 128.1, 128.0, 67.2, 65.0, 61.7, 53.4, 50.5, 47.2, 40.5, 38.2, 35.3, 34.8, 30.6, 29.6, 26.5.

**Benzyl ((S)-1-(((S)-4,4-dimethyl-1-oxo-1-(((S)-1-oxo-3-((S)-2-oxopyrrolidin-3-yl)propan-2-yl)amino)pentan-2-yl)amino)-3,3-dimethyl-1-oxobutan-2-yl)carbamate (MPI18). MPI18** was prepared as a white solid following a similar procedure to **MPI11** (yield 67%). $^1$H NMR (400 MHz, Chloroform-*d*) δ 9.50 (s, 1H), 8.16 – 7.99 (m, 1H), 7.42 – 7.28 (m,, 5H), 6.90 – 6.74(m, 1H), 5.55 (dd, *J* = 22.5, 9.6 Hz, 1H), 5.08 (d, *J* = 2.7 Hz, 2H), 4.72 – 4.58 (m, 1H), 4.38 (dd, *J* = 10.6, 5.9 Hz, 1H), 3.96 (dd, *J* = 13.1, 9.4 Hz, 1H), 3.47 – 3.25 (m, 2H), 2.51 – 2.29 (m, 2H), 2.09 – 1.77 (m, 4H), 1.53 – 1.43 (m, 1H), 1.13 – 0.71 (m, 18H). $^{13}$C NMR (100 MHz, Chloroform-

*d*): δ 199.3, 18.0, 173.7, 170.3, 156.5, 136.3, 128.5, 128.2, 127.9, 67.1, 62.6, 57.0, 50.8, 46.6, 40.5, 37.8, 36.5, 34.7, 30.6, 29.9, 29.5, 28.0, 26.5.

**Methyl (S)-2-((S)-2-(((benzyloxy)carbonyl)amino)-3,3-dimethylbutanamido)-3-cyclopropylpropanoate (MPI19b). MPI19b** was prepared with Cbz-L-tert-leucine and methyl (S)-2-amino-3-cyclopropylpropanoate hydrochloride as a colorless oil following a similar procedure to **MPI11b** (yield 73%).

**(S)-2-((S)-2-(((benzyloxy)carbonyl)amino)-3,3-dimethylbutanamido)-3-cyclopropylpropanoic acid (MPI19c). MPI19c** was prepared as a white solid following a similar procedure to **MPI11c** (yield 82%). $^1$H NMR (400 MHz, Chloroform-*d*) δ 7.41 – 7.28 (m, 5H), 6.89 (d, *J* = 7.6 Hz, 1H), 5.85 (d, *J* = 9.7 Hz, 1H), 5.10 (s, 2H), 4.67 (q, *J* = 7.7, 6.3 Hz, 1H), 4.16 (d, *J* = 9.7 Hz, 1H), 1.72 (dtd, *J* = 27.3, 13.9, 6.4 Hz, 2H), 0.99 (s, 9H), 0.67 (pd, *J* = 7.5, 3.7 Hz, 1H), 0.42 (td, *J* = 8.1, 5.0 Hz, 2H), 0.07 (td, *J* = 4.7, 2.3 Hz, 2H).

**Methyl (5S,8S,11S)-5-(tert-butyl)-8-(cyclopropylmethyl)-3,6,9-trioxo-11-(((S)-2-oxopyrrolidin-3-yl)methyl)-1-phenyl-2-oxa-4,7,10-triazadodecan-12-oate (MPI19d). MPI19d** was prepared as a white solid following a similar procedure to **MPI11d** (yield 64%). $^1$H NMR (400 MHz, Chloroform-*d*) δ 7.80 (d, *J* = 7.4 Hz, 1H), 7.41 – 7.28 (m, 5H), 6.96 (d, *J* = 8.3 Hz, 1H), 6.33 (d, *J* = 9.9 Hz, 1H), 5.55 (t, *J* = 10.4 Hz, 1H), 5.09 (s, 2H), 4.67 – 4.48 (m, 2H), 3.96 (t, *J* = 9.2 Hz, 1H), 3.71 (s, 3H), 3.43 – 3.27 (m, 2H), 2.53 – 2.32 (m, 2H), 2.17 (ddd, J = 16.5, 12.2, 4.8 Hz, 1H), 1.87 (dddd, J = 17.6, 11.4, 8.6, 2.9 Hz, 1H), 1.80 – 1.68 (m, 2H), 1.60 (dt, *J* = 13.9, 7.0 Hz, 1H), 0.99 (s, 9H), 0.73 (td, *J* = 7.5, 3.9 Hz, 1H), 0.42 (ddq, *J* = 11.4, 7.7, 3.8 Hz, 2H), 0.16 – 0.01 (m, 2H).

**Benzyl ((S)-1-(((S)-3-cyclopropyl-1-(((S)-1-hydroxy-3-((S)-2-oxopyrrolidin-3-yl)propan-2-yl)amino)-1-oxopropan-2-yl)amino)-3,3-dimethyl-1-oxobutan-2-yl)carbamate (MPI19e). MPI19e** was prepared as a white solid following a similar procedure to **MPI11e**. It was used in the next step without further purification.

**Benzyl ((S)-1-(((S)-3-cyclopropyl-1-oxo-1-(((S)-1-oxo-3-((S)-2-oxopyrrolidin-3-yl)propan-2-yl)amino)propan-2-yl)amino)-3,3-dimethyl-1-oxobutan-2-yl)carbamate (MPI19). MPI19** was prepared as a white solid following a similar procedure to **MPI11** (yield 48%). $^1$H NMR (400 MHz, Chloroform-*d*) δ 9.51 (s, 1H), 8.20 (d, *J* = 6.7 Hz, 1H), 7.32 (d, *J* = 4.2 Hz, 5H), 7.17 (d, *J* = 8.3 Hz, 1H), 6.67 (s, 1H), 5.68 (d, J = 9.4 Hz, 1H), 5.08 (s, 2H), 4.67 (q, J = 7.3 Hz, 1H), 4.42 (p, *J* = 4.9 Hz, 1H), 4.02 (d, *J* = 9.4 Hz, 1H), 3.31 (dq, *J* = 17.4, 9.5 Hz, 2H), 2.44 (p, *J* = 8.1 Hz, 1H), 2.34 (dt, *J* = 14.9, 8.5 Hz, 1H), 2.07 – 1.86 (m, 2H), 1.86 – 1.74 (m, 1H), 1.65 (ddt, J = 20.5, 13.6, 6.9 Hz, 2H), 0.98 (s, 9H), 0.70 (h, *J* = 7.1, 6.6 Hz, 1H), 0.43 (d, *J* = 8.0 Hz, 2H), 0.08 (d, *J* = 5.1 Hz, 2H). $^{13}$C NMR (100 MHz, Chloroform-*d*) δ 199.3, 180.0, 172.6, 170.4, 156.6, 136.3, 128.5, 128.2, 128.0, 77.3, 67.1, 62.9, 57.4, 53.7, 40.6, 37.9, 37.7, 34.7, 29.9, 28.4, 26.5, 7.3, 4.5, 4.3.

**(S)-methyl 2-((S)-2-(((benzyloxy)carbonyl)amino)-2-cyclopropylacetamido)-4,4-dimethylpentanoate (MPI20b). MPI20b** was obtained from methyl (S)-2-amino-4,4-dimethylpentanoate hydrochloride and (S)-2-(((benzyloxy)carbonyl)amino)-2-

cyclopropylacetic acid following a similar procedure to **MPI11b.** The crude product was purified by silica gel column chromatography (15-50% EtOAc in n-hexane) to afford the **MPI20b** as a white solid (yield 80%). $^{1}$H NMR (400 MHz, Chloroform-*d*): δ 7.42 – 7.28 (m, 5H), 6.53 – 6.37 (m, 1H), 5.63 – 5.44 (m, 1H), 5.09 (s, 2H), 4.64 (td, *J* = 8.7, 3.5 Hz, 1H), 3.71 (s, 3H), 3.68 – 3.60 (m, 1H), 1.77 (dd, *J* = 14.4, 3.6 Hz, 1H), 1.48 (dd, *J* = 14.4, 8.8 Hz, 1H), 1.22 – 1.07 (m, 1H), 0.93 (s, 9H), 0.72 – 0.41 (m, 4H). $^{13}$C NMR (100 MHz, Chloroform-*d*): δ 173.59, 170.61, 156.26, 136.20, 128.54, 128.17, 128.03, 67.06, 60.43, 52.39, 49.90, 46.06, 30.72, 29.50, 14.21, 3.60, 2.99.

**(S)-2-((S)-2-(((benzyloxy)carbonyl)amino)-2-cyclopropylacetamido)-4,4-dimethylpentanoic acid (MPI20c). MPI20c** was prepared as a white solid following a similar procedure to **MPI11c** (yield 83%). $^{1}$H NMR (400 MHz, Chloroform-*d*) δ 7.41 – 7.29 (m, 5H), 6.61 (d, *J* = 8.2 Hz, 1H), 5.72 (d, *J* = 7.6 Hz, 1H), 5.09 (s, 2H), 4.62 (td, *J* = 8.7, 3.0 Hz, 1H), 3.72 – 3.55 (m, 1H), 1.91 – 1.81 (m, 1H), 1.51 (dd, *J* = 14.5, 9.1 Hz, 1H), 1.20 – 1.06 (m, 1H), 0.94 (s, 9H), 0.69 – 0.36 (m, 4H). $^{13}$C NMR (100 MHz, Chloroform-*d*): δ 176.37, 171.31, 156.49, 136.06, 128.56, 128.24, 128.06, 67.97, 67.24, 50.01, 45.68, 30.77, 29.49, 25.61, 13.90, 3.66, 3.15.

**(5S,8S,11S)-methyl 5-cyclopropyl-8-neopentyl-3,6,9-trioxo-11-(((S)-2-oxopyrrolidin-3-yl)methyl)-1-phenyl-2-oxa-4,7,10-triazadodecan-12-oate (MPI20d). MPI20d** was prepared as a white solid following a similar procedure to **MPI11d** (yield 49%). $^{1}$H NMR (400 MHz, Chloroform-*d*): δ 7.89 – 7.74 (m, 1H), 7.30 – 7.19 (m, 5H), 6.75 (s, 1H), 5.85 – 5.67 (m, 1H), 5.00 (d, *J* = 5.2 Hz, 2H), 4.55 (td, *J* = 8.7, 3.5 Hz, 1H), 4.49 – 4.34 (m, 1H), 3.61 (s, 3H), 3.60 – 3.52 (m, 1H), 3.28 – 3.14 (m, 2H), 2.39 – 2.19 (m, 2H), 2.11 (td, *J* = 12.7, 11.7, 4.1 Hz, 1H), 1.87 – 1.61 (m, 3H), 1.12 – 0.98 (m, 1H), 0.85 (s, 9H), 0.57 – 0.23 (m, 4H). $^{13}$C NMR (100 MHz, Chloroform-*d*): δ 179.90, 173.11, 172.07, 170.88, 156.30, 136.24, 128.52, 128.15, 128.00, 67.02, 58.86, 55.27, 52.36, 50.89, 45.93, 40.57, 38.33, 30.55, 29.60, 28.01, 12.47, 3.30, 3.18.

**Benzyl ((S)-1-cyclopropyl-2-(((S)-1-(((S)-1-hydroxy-3-((S)-2-oxopyrrolidin-3-yl)propan-2-yl)amino)-4,4-dimethyl-1-oxopentan-2-yl)amino)-2-oxoethyl)carbamate (MPI20e). MPI20e** was prepared as a white solid following a similar procedure to **MPI11e** (yield 79%). $^{1}$H NMR (400 MHz, Chloroform-*d*): δ 7.70 – 7.42 (m, 2H), 7.38 – 7.21 (m, 5H), 6.79 (d, *J* = 5.1 Hz, 1H), 6.16 – 5.91 (m, 1H), 5.05 (q, *J* = 12.4 Hz, 2H), 4.62 – 4.43 (m, 1H), 4.06 – 3.87 (m, 1H), 3.63 – 3.54 (m, 2H), 3.24 (t, *J* = 9.2 Hz, 2H), 2.44 – 2.22 (m, 2H), 1.91 – 1.68 (m, 2H), 1.62 – 1.44 (m, 2H), 1.17 – 1.02 (m, 1H), 0.90 (s, 9H), 0.57 – 0.42 (m, 3H), 0.42 – 0.30 (m, 1H).

**Benzyl ((S)-1-cyclopropyl-2-(((S)-4,4-dimethyl-1-oxo-1-(((S)-1-oxo-3-((S)-2-oxopyrrolidin-3-yl)propan-2-yl)amino)pentan-2-yl)amino)-2-oxoethyl)carbamate (MPI20). MPI20** was prepared as a white solid following a similar procedure to **MPI11** (yield 62%). $^{1}$H NMR (400 MHz, Chloroform-*d*): δ 9.48 (s, 1H), 8.22 (s, 1H), 7.40 – 7.27 (m, 5H), 7.06 – 6.87 (m, 1H), 6.51 – 6.29 (m, 1H), 5.78 (s, 1H), 5.07 (s, 2H), 4.68 – 4.52 (m, 1H), 4.36 – 4.24 (m, 1H), 3.56 (ddd, *J* = 9.2, 7.0, 2.4 Hz, 1H), 3.36 – 3.19 (m, 2H), 2.50 – 2.28 (m, 2H), 2.07 – 1.84 (m, 4H), 1.84 – 1.72 (m, 1H), 1.21 – 1.07 (m, 1H), 0.95 (s, 9H), 0.66 – 0.35 (m, 4H). $^{13}$C NMR (100 MHz, Chloroform-*d*): δ 199.77,

180.13, 173.80, 171.01, 156.44, 136.16, 128.55, 128.21, 128.04, 67.10, 59.12, 57.47, 51.03, 45.90, 40.59, 38.03, 30.62, 30.01, 29.64, 28.29, 13.85, 3.29, 3.22.

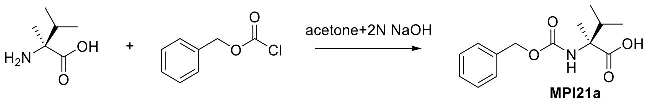


**(S)-2-(((benzyloxy)carbonyl)amino)-2,3-dimethylbutanoic acid (MPI21a).** H-L-(αMe)Val-OH (1.0 g, 7.63 mmol, 1.0 equiv) was dissolved in a 1:1 mixture of 1N NaOH (20 mL) : acetone (20 mL) and kept at 0 °C . Z-Cl (1.25 mL, 0.877 mmol, 0.1 equiv), diluted in acetone (5 mL), was added drop wise in 40 min and the pH was adjusted to 10.8-11.0 with 2N NaOH. After stirring the reaction mixture at room temperature for 5 h, Acetone was evaporated under reduce pressure and NaOH 2 N was added (20 mL). The unreacted Z-Cl was extracted with $Et_2O$. The aqueous solution was acidified with $KHSO_4$, the product was extracted with EtOAc (4x30 mL). The organic phase was washed with water (3x30 mL), and the solvent was evaporated at reduced pressure. The solid colorless product was obtained in 76% yield. $^1$H NMR (400 MHz, Chloroform-*d*): δ 10.38, 7.38, 7.37, 7.36, 7.36, 7.35, 7.34, 7.33, 5.46, 5.12, 1.61, 1.03, 1.01, 0.99, 0.98.

**(S)-methyl 2-((S)-2-(((benzyloxy)carbonyl)amino)-2,3-dimethylbutanamido)-4,4-dimethylpentanoate (MPI21b). MPI21b** was obtained from methyl (S)-2-amino-4,4-dimethylpentanoate hydrochloride and **MPI21a** following a similar procedure to **MPI11b.** The crude product was purified by silica gel column chromatography (15-50% EtOAc in n-hexane) to afford the **MPI21b** as a white solid (yield 78%). $^1$H NMR (400 MHz, Chloroform-*d*) δ 7.53 – 7.30 (m, 5H), 6.93 (s, 1H), 5.24 (s, 1H), 5.13 – 5.01 (m, 2H), 4.68 – 4.51 (m, 1H), 3.68 (s, 3H), 1.77 (dd, *J* = 14.4, 3.6 Hz, 1H), 1.50 (dd, *J* = 14.5, 8.9 Hz, 1H), 1.45 (s, 3H), 1.08 – 0.67 (m, 15H).

**(S)-2-((S)-2-(((benzyloxy)carbonyl)amino)-2,3-dimethylbutanamido)-4,4-dimethylpentanoic acid (MPI21c). MPI21c** was prepared as a white solid following a similar procedure to **MPI11c**. The crude **MPI21c** was used in the next step without further purification.

**(5S,8S,11S)-methyl 5-isopropyl-5-methyl-8-neopentyl-3,6,9-trioxo-11-(((S)-2-oxopyrrolidin-3-yl)methyl)-1-phenyl-2-oxa-4,7,10-triazadodecan-12-oate (MPI21d). MPI21d** was prepared as a white solid following a similar procedure to **MPI11d** (yield 63%). $^1$H NMR (400 MHz, Methanol-$d_4$) δ 8.35 (d, *J* = 8.1 Hz, 1H), 7.67 (d, *J* = 8.0 Hz, 1H), 7.37 – 7.13 (m, 5H), 6.93 (s, 1H), 5.14 – 4.93 (m, 2H), 4.43 – 4.19 (m, 2H), 3.60 (s, 3H), 3.20 – 3.10 (m, 2H), 2.45 – 2.33 (m, 1H), 2.28 – 2.18 (m, 1H), 1.97 – 1.87 (m, 1H), 1.77 – 1.61 (m, 3H), 1.50 (dd, *J* = 14.4, 8.3 Hz, 1H), 1.32 (s, 3H), 0.93 – 0.62 (m, 15H). $^{13}$C NMR (100 MHz, Methanol-$d_4$): δ 180.36, 174.44, 174.38, 172.08, 156.83, 136.80, 128.18, 128.09, 127.65, 127.62, 66.51, 62.93, 54.47, 51.65, 51.37, 50.73, 44.60, 42.43, 40.04, 38.06, 30.08, 28.71, 17.34, 17.12, 16.55, 16.16, 15.90.

**Benzyl ((S)-1-(((S)-1-(((S)-1-hydroxy-3-((S)-2-oxopyrrolidin-3-yl)propan-2-yl)amino)-4,4-dimethyl-1-oxopentan-2-yl)amino)-2,3-dimethyl-1-oxobutan-2-yl)carbamate (MPI21e). MPI21e** was prepared as a white solid following a similar procedure to **MPI11e** (yield 70%). $^1$H NMR (400 MHz, Chloroform-*d*) δ 7.41 – 7.27 (m, 5H), 6.56 (d, *J* = 6.3 Hz, 1H), 5.99 (s, 1H), 5.47 (s, 1H), 5.22 – 5.07 (m, 2H), 4.23 (ddd, *J* = 9.1, 6.2, 2.4 Hz, 1H), 4.03 (s, 1H), 3.74 – 3.64 (m, 1H), 3.63 – 3.54 (m, 1H), 3.36 – 3.19 (m, 2H), 2.46 – 2.29 (m, 2H), 2.22 – 1.96 (m, 2H), 2.01 – 1.88 (m, 2H), 1.76 (dd, *J* = 12.2, 9.0 Hz, 1H), 1.41 (s, 3H), 1.29 (dd, *J* = 14.7, 9.6 Hz, 1H), 1.02 – 0.77 (m, 15H). $^{13}$C NMR (100 MHz, Chloroform-*d*): δ 180.44, 173.62, 173.41, 156.25, 135.91, 128.67, 128.46, 128.37, 67.62, 65.70, 63.37, 53.46, 53.04, 50.63, 45.80, 40.33, 35.36, 32.36, 30.69, 29.69, 28.38, 17.45, 17.33, 17.11, 17.00.

**Benzyl ((S)-1-(((S)-4,4-dimethyl-1-oxo-1-(((S)-1-oxo-3-((S)-2-oxopyrrolidin-3-yl)propan-2-yl)amino)pentan-2-yl)amino)-2,3-dimethyl-1-oxobutan-2-yl)carbamate (MPI21). MPI21** was prepared as a white solid following a similar procedure to **MPI11** (yield 78%). $^1$H NMR (400 MHz, Chloroform-*d*) δ 9.52 (s, 1H), 8.07 (d, *J* = 7.7 Hz, 1H), 7.50 – 7.26 (m, 5H), 6.77 (d, *J* = 7.4 Hz, 1H), 6.39 – 6.20 (m, 1H), 5.47 (s, 1H), 5.07 (d, *J* = 3.2 Hz, 2H), 4.57 – 4.37 (m, 1H), 4.33 – 4.16 (m, 1H), 3.35 – 3.18 (m, 2H), 2.44 – 2.29 (m, 2H), 2.14 – 2.05 (m, 2H), 1.98 – 1.92 (m, 1H), 1.90 – 1.81 (m, 1H), 1.74 (t, *J* = 9.8 Hz, 1H), 1.40 (s, 3H), 0.97 – 0.85 (m, 15H). $^{13}$C NMR (100 MHz, Chloroform-*d*): δ 200.59, 173.83, 173.43, 173.17, 156.23, 136.06, 128.70, 128.63, 128.35, 128.16, 67.48, 67.28, 63.36, 57.12, 51.83, 45.78, 40.35, 37.64, 34.88, 30.65, 30.24, 29.67, 28.12, 17.44, 17.36, 17.09, 17.04.

**Methyl (S)-2-(2-(((benzyloxy)carbonyl)amino)-2-methylpropanamido)-4,4-dimethylpentanoate (MPI22b). MPI22b** was obtained from methyl (S)-2-amino-4,4-dimethylpentanoate hydrochloride and Z-Aib-OH following a similar procedure to **MPI11b.** The crude product was purified by silica gel column chromatography (15-50% EtOAc in hexanes as the eluent) to afford the **MPI22b** as a colorless oil (yield 79%). $^1$H NMR (400 MHz, Chloroform-*d*) δ 7.34 (s, 5H), 6.68 (s, 1H), 5.31 (s, 1H), 5.08 (s, 2H), 4.57 (t, *J* = 8.4 Hz, 1H), 3.69 (s, 3H), 1.91 – 1.64 (m, 2H), 1.53 (d, *J* = 6.0 Hz, 6H), 0.94 (s, 9H).

**(S)-2-(2-(((benzyloxy)carbonyl)amino)-2-methylpropanamido)-4,4-dimethylpentanoic acid (MPI22c). MPI22c** was prepared as a white solid following a similar procedure to **MPI11c** (yield 85%).

**Methyl (8S,11S)-5,5-dimethyl-8-neopentyl-3,6,9-trioxo-11-(((S)-2-oxopyrrolidin-3-yl)methyl)-1-phenyl-2-oxa-4,7,10-triazadodecan-12-oate (MPI22d). MPI22d** was prepared as a white solid following a similar procedure to **MPI11d** (yield 66%). $^1$H NMR (400 MHz, Chloroform-*d*) δ 7.81 (d, *J* = 7.8 Hz, 1H), 7.41 – 7.29 (m, 5H), 6.67 (d, *J* = 8.2 Hz, 1H), 5.77 (s, 1H), 5.43 (s, 1H), 5.17 – 5.01 (m, 2H), 4.63 – 4.43 (m, 2H), 3.71 (s, 3H), 3.40 – 3.16 (m, 2H), 2.48 – 2.32 (m, 2H), 2.30 – 2.16 (m, 1H), 2.15 – 1.95 (m, 1H), 1.95 – 1.82 (m, 2H), 1.52 (d, *J* = 11.7 Hz, 6H), 1.49 – 1.36 (m, 1H), 0.95 (s, 9H).

**Benzyl (1-(((S)-1-(((S)-1-hydroxy-3-((S)-2-oxopyrrolidin-3-yl)propan-2-yl)amino)-4,4-dimethyl-1-oxopentan-2-yl)amino)-2-methyl-1-oxopropan-2-yl)carbamate (MPI22e). MPI22e** was prepared as a white solid following a similar procedure to **MPI11e**. The crude **MPI22e** was used in the next step without further purification.

**Benzyl (1-(((S)-4,4-dimethyl-1-oxo-1-(((S)-1-oxo-3-((S)-2-oxopyrrolidin-3-yl)propan-2-yl)amino)pentan-2-yl)amino)-2-methyl-1-oxopropan-2-yl)carbamate (MPI22).** **MPI22** was prepared as a white solid following a similar procedure to **MPI11** (yield 50%). $^{1}$H NMR (400 MHz, Chloroform-*d*) δ 9.52 (s, 1H), 8.15 (d, *J* = 7.6 Hz, 1H), 7.33 (s, 5H), 6.83 (d, *J* = 8.0 Hz, 1H), 6.12 (s, 1H), 5.71 (s, 1H), 5.05 (q, *J* = 12.2 Hz, 2H), 4.58 – 4.42 (m, 1H), 4.23 (ddd, *J* = 11.3, 7.6, 3.9 Hz, 1H), 3.33 – 3.16 (m, 2H), 2.49 – 2.29 (m, 2H), 2.13 – 1.97 (m, 2H), 1.96 – 1.82 (m, 2H), 1.82 – 1.68 (m, 1H), 1.47 (d, *J* = 6.8 Hz, 6H), 0.94 (s, 9H). $^{13}$C NMR (100 MHz, Chloroform-*d*) δ 200.5, 180.0, 173.8, 156.0, 136.0, 128.6, 128.3, 128.1, 67.2, 57.3, 57.1, 51.6, 45.5, 40.4, 37.7, 30.7, 30.2, 29.6, 28.3, 26.0.

**Methyl (S)-2-(1-(((benzyloxy)carbonyl)amino)cyclopropane-1-carboxamido)-3-cyclohexylpropanoate (MPI23b).** **MPI23b** was obtained from H-Cha-OMe hydrochloride and 1-(((benzyloxy)carbonyl)amino)cyclopropane-1-carboxylic acid following a similar procedure to **MPI11b.** The crude product was purified by silica gel column chromatography (50-100% EtOAc in hexanes as the eluent) to afford the **MPI23b**. $^{1}$H NMR (400 MHz, Chloroform-*d*) δ 7.29 (d, *J* = 3.9 Hz, 5H), 6.72 (s, 1H), 5.05 (d, *J* = 12.0 Hz, 2H), 4.60 – 4.33 (m, 1H), 3.63 (s, 3H), 1.78 – 0.66 (m, 18H). $^{13}$C NMR (100 MHz, Chloroform-*d*) δ 173.61, 171.53, 135.97, 128.63, 52.27, 50.58, 40.06, 34.09, 33.38, 32.57, 26.35, 26.15, 25.98.

**(S)-2-(1-(((benzyloxy)carbonyl)amino)cyclopropane-1-carboxamido)-3-cyclohexylpropanoic acid (MPI23c).** **MPI23c** was prepared as a white solid following a similar procedure to **MPI11c**. $^{1}$H NMR (400 MHz, Chloroform-*d*) δ 7.26 (s, 5H), 6.84 (d, *J* = 11.3 Hz, 1H), 5.80 (s, 1H), 5.25 – 4.94 (m, 2H), 4.49 (s, 1H), 1.59 (dd, *J* = 13.3, 6.7 Hz, 6H), 1.51 – 1.24 (m, 3H), 1.20 (s, 2H), 1.06 – 0.91 (m, 4H), 0.81 (s, 3H). $^{13}$C NMR (100 MHz, Chloroform-*d*) δ 176.01, 172.64, 128.63, 128.35, 67.96, 50.83, 34.07, 33.40, 32.43, 29.72, 26.34, 26.13, 25.95, 25.60.

**Methyl (S)-2-((S)-2-(1-(((benzyloxy)carbonyl)amino)cyclopropane-1-carboxamido)-3-cyclohexylpropanamido)-3-((S)-2-oxopyrrolidin-3-yl)propanoate (MPI23d).** **MPI23d** was prepared as a white solid following a similar procedure to **MPI11d** (yield 62%). $^{1}$H NMR (400 MHz, Methanol-*d*$_4$) δ 7.32 – 7.13 (m, 5H), 5.14 – 4.94 (m, 2H), 4.47 – 4.26 (m, 2H), 3.62 (s, 3H), 3.13 (d, *J* = 7.5 Hz, 2H), 2.39 (d, *J* = 10.3 Hz, 1H), 2.22 – 2.02 (m, 2H), 1.82 – 1.43 (m, 9H), 1.43 – 1.30 (m, 2H), 1.17 (d, *J* = 12.4 Hz, 2H), 1.11 (q, *J* = 8.4, 7.6 Hz, 2H), 0.96 (d, *J* = 8.3 Hz, 2H), 0.89 – 0.70 (m, 2H).

**Benzyl (1-(((S)-3-cyclohexyl-1-(((S)-1-hydroxy-3-((S)-2-oxopyrrolidin-3-yl)propan-2-yl)amino)-1-oxopropan-2-yl)carbamoyl)cyclopropyl)carbamate (MPI23e).** **MPI23e** was prepared as a white solid following a similar procedure to **MPI11e** (yield 61%). $^{1}$H NMR (400 MHz, Methanol-*d*$_4$) δ 7.32 – 7.09 (m, 5H), 5.09 (d, *J* = 12.4 Hz, 1H), 4.98 (d, *J* = 12.4 Hz, 1H), 4.35 (dd, *J* = 10.4, 4.9 Hz, 1H), 3.96 – 3.74 (m, 1H), 3.42 (dd, *J* = 11.5, 5.9 Hz, 2H), 2.37 – 2.14 (m, 2H), 1.87 (t, *J* = 12.5 Hz, 1H),

1.70 – 1.41 (m, 9H), 1.42 – 1.28 (m, 2H), 1.28 – 1.19 (m, 3H), 1.19 – 0.97 (m, 4H), 0.97 – 0.72 (m, 3H). $^{13}$C NMR (100 MHz, Methanol-*d*$_4$) δ 181.30, 173.80, 173.46, 136.60, 128.14, 127.59, 66.73, 64.06, 48.26, 48.05, 47.84, 47.63, 47.41, 47.20, 46.99, 40.16, 38.24, 35.21, 34.08, 33.57, 27.55, 26.19, 26.04, 25.80.

**Benzyl (1-(((S)-3-cyclohexyl-1-oxo-1-(((S)-1-oxo-3-((S)-2-oxopyrrolidin-3-yl)propan-2-yl)amino)propan-2-yl)carbamoyl)cyclopropyl)carbamate (MPI23).** **MPI23** was prepared as a white solid following a similar procedure to **MPI11** (yield 41%). $^1$H NMR (400 MHz, Chloroform-*d*) δ 9.43 (d, *J* = 24.1 Hz, 1H), 8.30 (s, 1H), 7.27 (d, *J* = 4.2 Hz, 5H), 6.86 (d, *J* = 56.9 Hz, 1H), 6.25 (s, 1H), 5.88 (d, *J* = 34.9 Hz, 1H), 5.18 – 4.90 (m, 2H), 4.56 (q, *J* = 4.1 Hz, 1H), 4.51 – 4.35 (m, 1H), 4.20 (s, 1H), 3.24 (d, *J* = 10.0 Hz, 2H), 2.45 – 2.20 (m, 2H), 2.03 – 1.66 (m, 7H), 1.52 – 1.35 (m, 3H), 1.22 (d, *J* = 9.4 Hz, 1H), 1.14 – 1.00 (m, 4H), 0.98 – 0.67 (m, 4H).

**(S)-methyl 2-((S)-2-(((benzyloxy)carbonyl)amino)-2-cyclopropylacetamido)-3-cyclopropylpropanoate (MPI24b).** **MPI24b** was obtained from methyl (S)-2-amino-3-cyclopropylpropanoate hydrochloride and (S)-2-(((benzyloxy)carbonyl)amino)-2-cyclopropylacetic acid following a similar procedure to **MPI11b.** The crude product was purified by silica gel column chromatography (15-50% EtOAc in hexanes as the eluent) to afford the **MPI24b** (yield 80%). $^1$H NMR (400 MHz, Chloroform-*d*): δ 7.58 – 7.21 (m, 5H), 6.77 (s, 1H), 5.68 – 5.49 (m, 1H), 5.10 (s, 2H), 4.66 (q, *J* = 6.4 Hz, 1H), 3.75 (s, 3H), 3.71 – 3.59 (m, 1H), 1.70 (q, *J* = 6.4 Hz, 2H), 1.20 – 1.04 (m, 1H), 0.68 – 0.41 (m, 6H), 0.20 – -0.02 (m, 2H).$^{13}$C NMR (100 MHz, Chloroform-*d*): δ 172.51, 171.10, 156.27, 136.21, 128.52, 128.15, 127.98, 67.05, 58.58, 52.79, 52.36, 36.91, 14.30, 6.82, 4.20, 4.10, 3.57, 3.16.

**(S)-2-((S)-2-(((benzyloxy)carbonyl)amino)-2-cyclopropylacetamido)-3-cyclopropylpropanoic acid (MPI24c).** **MPI24c** was prepared as a white solid following a similar procedure to **MPI11c** (yield 85%). $^1$H NMR (400 MHz, DMSO-*d*$_6$) δ 7.96 (d, *J* = 7.9 Hz, 1H), 7.43 (d, *J* = 8.2 Hz, 1H), 7.34 – 7.16 (m, 5H), 4.96 (s, 2H), 4.23 (td, *J* = 7.8, 5.3 Hz, 1H), 3.57 (t, *J* = 8.3 Hz, 1H), 1.64 – 1.35 (m, 2H), 1.07 – 0.87 (m, 1H), 0.82 – 0.64 (m, 1H), 0.50 – 0.19 (m, 6H), 0.12 – -0.09 (m, 2H). $^{13}$C NMR (100 MHz, DMSO-*d*$_6$): δ 173.78, 171.30, 156.15, 137.51, 128.78, 128.22, 128.13, 65.83, 58.15, 52.78, 36.62, 14.20, 8.04, 4.85, 4.53, 3.54, 3.00.

**(5S,8S,11S)-methyl 5-cyclopropyl-8-(cyclopropylmethyl)-3,6,9-trioxo-11-(((S)-2-oxopyrrolidin-3-yl)methyl)-1-phenyl-2-oxa-4,7,10-triazadodecan-12-oate (MPI24d).** **MPI24d** was prepared as a white solid following a similar procedure to **MPI11d** (yield 50%). $^1$H NMR (400 MHz, Chloroform-*d*): δ 8.03 (d, *J* = 7.2 Hz, 1H), 7.37 – 7.26 (m, 5H), 7.19 (d, *J* = 8.0 Hz, 1H), 6.53 (s, 1H), 5.83 (d, *J* = 7.4 Hz, 1H), 5.08 (d, *J* = 3.0 Hz, 2H), 4.63 (q, *J* = 7.1 Hz, 1H), 4.57 – 4.43 (m, 1H), 3.70 (s, 4H), 3.40 – 3.18 (m, 2H), 2.51 – 2.28 (m, 2H), 2.26 – 2.09 (m, 1H), 1.95 – 1.76 (m, 2H), 1.76 – 1.55 (m, 2H), 1.52 – 1.38 (m, 1H), 1.20 – 1.03 (m, 1H), 0.79 – 0.66 (m, 1H), 0.65 – 0.33 (m, 6H), 0.17 – -0.03 (m, 2H). $^{13}$C NMR (100 MHz, Chloroform-*d*): δ 179.99, 172.09, 171.94, 170.97, 156.32, 136.26, 128.53, 128.15, 128.03, 67.01, 54.81, 53.65, 52.43, 51.53, 40.60, 38.51, 37.45, 33.14, 28.31, 14.36, 7.10, 4.49, 4.22, 3.49, 3.20.

**Benzyl ((S)-1-cyclopropyl-2-(((S)-3-cyclopropyl-1-(((S)-1-hydroxy-3-((S)-2-oxopyrrolidin-3-yl)propan-2-yl)amino)-1-oxopropan-2-yl)amino)-2-oxoethyl)carbamate (MPI24e). MPI24e** was prepared as a white solid following a similar procedure to **MPI11e** (yield 79%). $^1$H NMR (400 MHz, Methanol-$d_4$): δ 7.34 – 6.98 (m, 5H), 5.06 – 4.87 (m, 2H), 4.23 (d, $J$ = 7.3 Hz, 1H), 3.85 (d, $J$ = 10.5 Hz, 1H), 3.48 – 3.27 (m, 3H), 3.16 – 3.09 (m, 1H), 2.41 – 2.25 (m, 1H), 2.27 – 2.13 (m, 1H), 1.88 – 1.77 (m, 1H), 1.70 – 1.31 (m, 4H), 1.05 – 0.90 (m, 1H), 0.73 – 0.59 (m, 1H), 0.53 – 0.20 (m, 6H), 0.15 – -0.11 (m, 2H).

**Benzyl ((S)-1-cyclopropyl-2-(((S)-3-cyclopropyl-1-oxo-1-(((S)-1-oxo-3-((S)-2-oxopyrrolidin-3-yl)propan-2-yl)amino)propan-2-yl)amino)-2-oxoethyl)carbamate (MPI24). MPI24** was prepared as a white solid following a similar procedure to **MPI11** (yield 66%). $^1$H NMR (400 MHz, DMSO-$d_6$): δ 9.35 (s, 1H), 8.38 (d, $J$ = 7.7 Hz, 1H), 7.91 (dd, $J$ = 22.3, 7.8 Hz, 1H), 7.57 (s, 1H), 7.52 – 7.42 (m, 1H), 7.35 – 7.16 (m, 5H), 4.94 (s, 2H), 4.34 – 4.20 (m, 1H), 4.20 – 4.07 (m, 1H), 3.57 – 3.41 (m, 1H), 3.15 – 2.91 (m, 2H), 2.30 – 2.14 (m, 1H), 2.14 – 1.99 (m, 1H), 1.81 (q, $J$ = 16.5, 14.7 Hz, 1H), 1.64 – 1.27 (m, 4H), 1.04 – 0.85 (m, 1H), 0.74 – 0.55 (m, 1H), 0.48 – 0.16 (m, 6H), 0.08 – -0.11 (m, 2H). $^{13}$C NMR (100 MHz, DMSO-$d_6$): δ 201.20, 178.70, 172.56, 171.22, 156.23, 137.50, 128.79, 128.24, 128.15, 65.84, 58.23, 56.67, 53.57, 37.63, 29.81, 27.74, 14.07, 8.05, 4.88, 4.69, 3.59, 2.87.

**(S)-methyl 2-((2S,3R)-2-(((benzyloxy)carbonyl)amino)-3-(tert-butoxy)butanamido)-3-cyclohexylpropanoate (MPI25b). MPI25b** was obtained from methyl (S)-2-amino-3-cyclohexylpropanoate hydrochloride and N-((benzyloxy)carbonyl)-O-(tert-butyl)-L-allothreonine following a similar procedure to **MPI11b.** The crude product was purified by silica gel column chromatography (15-50% EtOAc in hexanes as the eluent) to afford the **MPI25b** (yield 84%).$^1$H NMR (400 MHz, Chloroform-$d$) δ 7.65 (d, $J$ = 7.7 Hz, 1H), 7.43 – 7.29 (m, 5H), 5.94 (d, $J$ = 5.0 Hz, 1H), 5.27 – 4.90 (m, 2H), 4.52 (td, $J$ = 8.4, 5.0 Hz, 1H), 4.27 – 4.16 (m, 2H), 3.72 (s, 3H), 1.79 – 1.61 (m, 7H), 1.60 – 1.47 (m, 1H), 1.30 (s, 9H), 1.22 – 1.14 (m, 3H), 1.11 (d, $J$ = 6.0 Hz, 3H), 1.01 – 0.85 (m, 2H). $^{13}$C NMR (100 MHz, $CDCl_3$) δ 173.07, 169.43, 156.15, 136.28, 128.55, 128.15, 127.96, 75.57, 66.83, 60.42, 58.37, 52.16, 50.51, 39.76, 34.30, 33.51, 32.56, 28.20, 26.32, 26.18, 25.99, 21.07, 16.41.

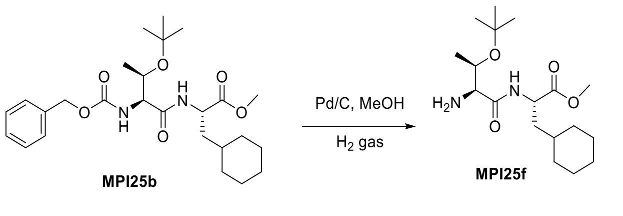


**(S)-methyl 2-((2S,3R)-2-amino-3-(tert-butoxy)butanamido)-3-cyclohexylpropanoate (MPI25f).** To a solution of **MPI25b** (2.73 mmol, 1.3 g) in methanol (20 mL) was added 10 % Pd/C (150 mg). The mixture was then stirred with hydrogen balloon at room temperature for 12 h. The catalyst was then filtered off and the solution was evaporated on vacuum to afford **MPI25f** as white solid, which was used without purification. $^1$H NMR (400 MHz, Methanol-$d_4$) δ 4.49 (dd, $J$ = 9.4, 5.4 Hz, 1H), 4.01 – 3.90 (m, 1H), 3.73 (s, 3H), 3.41 – 3.28 (m, 1H), 3.20 (d, $J$ = 4.5 Hz,

1H), 1.90 – 1.59 (m, 7H), 1.51 – 1.37 (m, 1H), 1.23 (d, $J$ = 9.9 Hz, 15H), 1.11 – 0.89 (m, 2H). $^{13}$C NMR (100 MHz, MeOD): δ 174.31, 173.15, 73.93, 68.77, 59.84, 51.23, 50.10, 38.97, 33.92, 33.35, 32.03, 27.59, 26.10, 25.94, 25.70, 18.68.

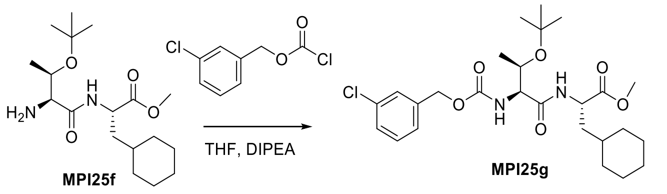


**(S)-methyl 2-((2S,3R)-3-(tert-butoxy)-2-((((3-chlorobenzyl)oxy)carbonyl)amino)butanamido)-3-cyclohexylpropanoate (MPI25g).** To 3,5-dichlorobenzyl alcohol (0.201 g, 1.39 mmol) in THF (5 mL) were added $K_2CO_3$ (193 mg, 1.39 mmol) and Triphosgene (166 mg, 0.56 mmol) and the mixture was stirred at room temperature for 1 h. The mixture was then poured into water (10 mL) and extracted with ethyl acetate (2×20 mL), combine organic layers and dried over $Na_2SO_4$. The organic phase was evaporated to dryness and the crude material was used directly in the next step. 3,5-Dichlorobenzyl Chloroformate in THF (5 mL) was added to drop wise to a mixture of compound **MPI25f** (400 mg,1.39mmol) and DIPEA (0.47 ml, 2.78mmol). The reaction mixture stirred for 12 h. The mixture was then poured into water (30 mL) and extracted with ethyl acetate (4×20 mL). The organic layer was washed with aqueous hydrochloric acid 10% v/v (2×20 mL), saturated aqueous $NaHCO_3$ (2×20 mL), brine (2×20 mL) and dried over $Na_2SO_4$. The organic phase was evaporated to dryness and the crude material purified by silica gel column chromatography (15-50% EtOAc in n-hexane as the eluent) to afford **MPI25g** white solid (410 mg, 70%). $^1$H NMR (400 MHz, Chloroform-*d*) δ 7.69 (d, $J$ = 7.7 Hz, 1H), 7.44 – 7.22 (m, 4H), 6.00 (d, $J$ = 4.9 Hz, 1H), 5.19 – 5.03 (m, 2H), 4.55 (td, $J$ = 8.4, 5.1 Hz, 1H), 4.22 (q, $J$ = 6.9, 5.5 Hz, 2H), 3.75 (s, 3H), 1.88 – 1.60 (m, 7H), 1.46 – 1.11 (m, 16H), 1.04 – 0.89 (m, 2H). $^{13}$C NMR (100 MHz, $CDCl_3$) δ 173.04, 171.18, 169.33, 155.86, 138.37, 134.42, 129.83, 128.24, 127.85, 125.81, 75.62, 66.84, 65.85, 60.42, 58.34, 52.17, 50.54, 39.75, 34.68, 34.31, 33.51, 32.56, 31.61, 28.19, 26.32, 26.18, 16.36.

**(S)-2-((2S,3R)-3-(tert-butoxy)-2-((((3-chlorobenzyl)oxy)carbonyl)amino)butanamido)-3-cyclohexylpropanoic acid (MPI25c). MPI25c** was prepared as a white solid following a similar procedure to **MPI11c** (yield 82%). $^1$H NMR (400 MHz, Chloroform-*d*) δ 7.57 (d, $J$ = 7.3 Hz, 1H), 7.39 – 7.04 (m, 4H), 6.09 (d, $J$ = 5.8 Hz, 1H), 5.16 – 4.83 (m, 2H), 4.36 (td, $J$ = 8.1, 7.6, 4.7 Hz, 1H), 4.20 – 4.01 (m, 2H), 1.72 – 1.46 (m, 7H), 1.33 – 0.95 (m, 16H), 0.95 – 0.76 (m, 2H). $^{13}$C NMR (100 MHz, $CDCl_3$): δ 177.40, 170.06, 156.11, 138.37, 134.39, 129.82, 128.21, 127.85, 125.84, 75.56, 67.03, 65.96, 58.65, 39.41, 34.33, 33.65, 32.35, 28.20, 26.34, 26.20, 25.97, 16.82.

**(5S,8S,11S)-methyl 5-((R)-1-(tert-butoxy)ethyl)-1-(3-chlorophenyl)-8-(cyclohexylmethyl)-3,6,9-trioxo-11-(((S)-2-oxopyrrolidin-3-yl)methyl)-2-oxa-4,7,10-triazadodecan-12-oate (MPI25d). MPI25d** was prepared as a white solid following a similar procedure to **MPI11d** (yield 64%). $^1$H NMR (400 MHz, Chloroform-*d*) δ 7.48 (dd, $J$ = 52.5, 7.6 Hz, 2H), 7.36 – 7.01 (m, 4H), 6.30 (s, 1H), 5.90

(d, *J* = 5.1 Hz, 1H), 5.01 (q, *J* = 12.6 Hz, 2H), 4.54 – 4.33 (m, 2H), 4.11 (d, *J* = 5.7 Hz, 2H), 3.65 (s, 3H), 3.39 – 3.18 (m, 2H), 2.46 – 2.25 (m, 2H), 2.20 – 2.03 (m, 1H), 1.87 – 1.40 (m, 9H), 1.11 (d, *J* = 72.3 Hz, 16H), 0.97 – 0.79 (m, 2H). $^{13}$C NMR (100 MHz, $CDCl_3$) δ 179.92, 172.23, 172.12, 169.47, 155.95, 138.30, 134.42, 129.86, 128.28, 127.93, 125.93, 75.51, 66.73, 65.97, 58.82, 52.43, 51.49, 51.19, 40.55, 39.95, 38.64, 38.24, 34.10, 33.62, 33.13, 32.66, 28.30, 28.23, 26.39, 26.23, 26.07, 17.15.

**3-chlorobenzyl ((2S,3R)-3-(tert-butoxy)-1-(((S)-3-cyclohexyl-1-(((S)-1-hydroxy-3-((S)-2-oxopyrrolidin-3-yl)propan-2-yl)amino)-1-oxopropan-2-yl)amino)-1-oxobutan-2-yl)carbamate (MPI25e). MPI25e** was prepared as a white solid following a similar procedure to **MPI11e** (yield 67%). $^{1}$H NMR (400 MHz, Chloroform-*d*) δ 7.54 (d, *J* = 7.6 Hz, 1H), 7.34 – 7.10 (m, 4H), 6.10 (d, *J* = 5.4 Hz, 1H), 5.89 (s, 1H), 5.01 (q, *J* = 12.6 Hz, 2H), 4.32 (q, *J* = 7.7 Hz, 1H), 4.18 – 4.03 (m, 2H), 3.94 (s, 1H), 3.55 (t, *J* = 4.0 Hz, 2H), 3.36 – 3.12 (m, 2H), 2.44 – 2.23 (m, 4H), 2.05 – 1.90 (m, 1H), 1.78 – 1.41 (m, 8H), 1.11 (d, *J* = 68.1 Hz, 15H), 0.93 – 0.77 (m, 2H). $^{13}$C NMR (100 MHz, $CDCl_3$) δ 180.95, 172.81, 169.73, 156.11, 138.27, 134.42, 129.88, 128.31, 127.97, 125.97, 75.52, 66.73, 66.06, 65.92, 59.13, 51.89, 40.52, 39.74, 38.21, 34.25, 33.62, 32.61, 28.63, 28.24, 26.36, 26.23, 26.03, 17.44.

**3-chlorobenzyl ((2S,3R)-3-(tert-butoxy)-1-(((S)-3-cyclohexyl-1-oxo-1-(((S)-1-oxo-3-((S)-2-oxopyrrolidin-3-yl)propan-2-yl)amino)propan-2-yl)amino)-1-oxobutan-2-yl)carbamate (MPI25). MPI25** was prepared as a white solid following a similar procedure to **MPI11** (yield 72%). $^{1}$H NMR (400 MHz, Chloroform-*d*) δ 9.44 (s, 1H), 7.99 (d, *J* = 6.4 Hz, 1H), 7.41 (d, *J* = 7.8 Hz, 1H), 7.29 (s, 1H), 7.22 (d, *J* = 4.6 Hz, 2H), 7.16 (t, *J* = 4.5 Hz, 1H), 6.11 (s, 1H), 5.89 (d, *J* = 5.1 Hz, 1H), 5.01 (q, *J* = 12.7 Hz, 2H), 4.48 – 4.37 (m, 1H), 4.35 – 4.25 (m, 1H), 4.11 (d, *J* = 5.5 Hz, 2H), 3.36 – 3.19 (m, 2H), 2.44 – 2.26 (m, 2H), 1.98 – 1.86 (m, 2H), 1.78 – 1.70 (m, 2H), 1.60 – 1.47 (m, 3H), 1.30 – 1.15 (m, 12H), 1.16 – 1.08 (m, 3H), 1.02 (d, *J* = 6.0 Hz, 3H), 0.96 – 0.82 (m, 3H). $^{13}$C NMR (100 MHz, $CDCl_3$) δ 199.57, 179.96, 172.90, 169.62, 155.99, 138.26, 134.43, 129.88, 128.32, 127.95, 125.95, 75.51, 66.72, 66.02, 58.93, 57.58, 51.50, 40.54, 40.03, 37.92, 34.24, 33.65, 32.62, 29.81, 29.71, 28.62, 28.24, 26.35, 26.22, 26.02, 17.29.

**(S)-methyl 2-((2S,3R)-2-(((benzyloxy)carbonyl)amino)-3-(tert-butoxy)butanamido)-4-methylpentanoate (MPI26b). MPI26b** was obtained from methyl L-leucinate hydrochloride and N-((benzyloxy)carbonyl)-O-(tert-butyl)-L-allothreonine following a similar procedure to **MPI11b.** The crude product was purified by silica gel column chromatography (15-50% EtOAc in hexanes as the eluent) to afford the **MPI26b** (yield 78%).$^{1}$H NMR (400 MHz, Chloroform-*d*) δ 7.66 (d, *J* = 7.8 Hz, 1H), 7.51 – 7.31 (m, 5H), 5.94 (d, *J* = 4.9 Hz, 1H), 5.26 – 4.95 (m, 2H), 4.57 – 4.42 (m, 1H), 4.28 – 4.16 (m, 2H), 3.72 (s, 3H), 1.75 – 1.52 (m, 3H), 1.30 (s, 9H), 1.11 (d, *J* = 6.2 Hz, 3H), 0.94 (dd, *J* = 6.1, 3.8 Hz, 6H).$^{13}$C NMR (100 MHz, $CDCl_3$): δ 172.95, 169.43, 156.11, 136.28, 128.54, 128.14, 127.95, 75.57, 66.89, 66.82, 60.40, 58.36, 52.15, 51.07, 41.27, 28.18, 25.01, 22.80, 21.95, 21.06, 16.39, 14.21.

**(S)-methyl 2-((2S,3R)-2-amino-3-(tert-butoxy)butanamido)-4-methylpentanoate (MPI26f). MPI26f** was prepared as a white solid following a similar procedure to **MPI125f**. $^1$H NMR (400 MHz, Chloroform-*d*) δ 7.88 (d, *J* = 8.1 Hz, 1H), 4.61 – 4.43 (m, 1H), 4.19 – 4.06 (m, 1H), 3.72 (s, 3H), 3.21 (d, *J* = 3.2 Hz, 1H), 1.75 – 1.51 (m, 3H), 1.18 (s, 12H), 1.04 – 0.86 (m, 6H). $^{13}$C NMR (100 MHz, $CDCl_3$): δ 173.53, 173.48, 74.14, 67.85, 59.58, 52.11, 50.73, 41.23, 28.53, 24.90, 22.85, 21.91, 19.27.

**(S)-methyl 2-((2S,3R)-3-(tert-butoxy)-2-((((3-chlorobenzyl)oxy)carbonyl)amino)butanamido)-4-methylpentanoate (MPI126g). MPI26g** was prepared as a white solid following a similar procedure to **MPI125g** (yield 51%). $^1$H NMR (400 MHz, Chloroform-*d*) δ 7.61 (d, *J* = 7.8 Hz, 1H), 7.36 – 7.09 (m, 4H), 5.91 (d, *J* = 5.0 Hz, 1H), 5.11 – 4.93 (m, 2H), 4.52 – 4.38 (m, 1H), 4.24 – 3.98 (m, 2H), 3.66 (s, 3H), 1.70 – 1.54 (m, 2H), 1.51 (q, *J* = 9.0, 8.4 Hz, 1H), 1.24 (s, 9H), 1.05 (d, *J* = 6.2 Hz, 3H), 0.87 (dd, *J* = 6.1, 3.6 Hz, 6H).$^{13}$C NMR (100 MHz, $CDCl_3$): δ 172.95, 169.36, 155.83, 138.36, 134.41, 129.83, 128.23, 127.85, 125.81, 75.64, 66.83, 65.84, 58.32, 52.19, 51.10, 41.24, 28.16, 25.02, 22.81, 21.95, 16.34.

**(S)-2-((2S,3R)-3-(tert-butoxy)-2-((((3-chlorobenzyl)oxy)carbonyl)amino)butanamido)-4-methylpentanoic acid (MPI126c). MPI26c** was prepared as a white solid following a similar procedure to **MPI11c.** $^1$H NMR (400 MHz, Chloroform-*d*) δ 7.63 (d, *J* = 7.5 Hz, 1H), 7.37 – 7.09 (m, 4H), 5.97 (d, *J* = 5.4 Hz, 1H), 5.01 (q, *J* = 12.7 Hz, 2H), 4.44 (q, *J* = 7.2, 6.0 Hz, 1H), 4.25 – 4.07 (m, 2H), 1.72 – 1.59 (m, 2H), 1.59 – 1.43 (m, 1H), 1.22 (s, 9H), 1.02 (d, *J* = 6.3 Hz, 3H), 0.95 – 0.80 (m, 6H). $^{13}$C NMR (100 MHz, $CDCl_3$): δ 177.19, 169.79, 155.91, 138.31, 134.41, 129.83, 128.25, 127.86, 125.82, 75.70, 67.95, 66.91, 65.92, 58.35, 51.08, 40.96, 28.14, 25.03, 22.84, 21.84, 16.42.

**(5S,8S,11S)-methyl 5-((R)-1-(tert-butoxy)ethyl)-1-(3-chlorophenyl)-8-isobutyl-3,6,9-trioxo-11-(((S)-2-oxopyrrolidin-3-yl)methyl)-2-oxa-4,7,10-triazadodecan-12-oate (MPI126d). MPI26d** was prepared as a white solid following a similar procedure to **MPI11d** (yield 60%). $^1$H NMR (400 MHz, Chloroform-*d*) δ 7.61 (d, *J* = 7.4 Hz, 1H), 7.42 (d, *J* = 7.9 Hz, 1H), 7.33 – 7.12 (m, 4H), 6.58 (s, 1H), 5.88 (d, *J* = 5.3 Hz, 1H), 5.01 (q, *J* = 12.6 Hz, 2H), 4.54 – 4.33 (m, 2H), 4.10 (d, *J* = 5.5 Hz, 2H), 3.65 (s, 3H), 3.36 – 3.18 (m, 2H), 2.52 – 2.25 (m, 2H), 2.17 – 2.05 (m, 1H), 1.87 – 1.71 (m, 2H), 1.67 – 1.59 (m, 2H), 1.49 (q, *J* = 8.9, 8.4 Hz, 1H), 1.20 (s, 9H), 1.00 (d, *J* = 6.0 Hz, 3H), 0.88 (dd, *J* = 11.7, 6.0 Hz, 6H). $^{13}$C NMR (100 MHz, $CDCl_3$): δ 179.96, 172.16, 172.07, 169.48, 155.92, 138.30, 134.41, 129.87, 128.29, 127.93, 125.94, 75.50, 66.73, 65.97, 58.83, 54.58, 52.43, 41.51, 40.82, 38.35, 33.08, 28.21, 24.78, 22.86, 22.12, 18.53, 17.18, 17.14.

**3-chlorobenzyl ((2S,3R)-3-(tert-butoxy)-1-(((S)-1-(((S)-1-hydroxy-3-((S)-2-oxopyrrolidin-3-yl)propan-2-yl)amino)-4-methyl-1-oxopentan-2-yl)amino)-1-oxobutan-2-yl)carbamate (MPI126e). MPI26e** was prepared as a white solid following a similar procedure to **MPI11e** (yield 73%). $^1$H NMR (400 MHz, Chloroform-*d*) δ 7.66 (d, *J* = 7.9 Hz, 1H), 7.40 (d, *J* = 7.8 Hz, 1H), 7.33 – 7.10 (m, 4H), 6.28 – 6.06 (m, 2H), 5.08 – 4.88 (m, 2H), 4.34 (td, *J* = 8.2, 5.2 Hz, 1H), 4.14 – 4.06 (m, 1H), 4.02 – 3.80 (m, 1H), 3.74 (t, *J* = 6.4 Hz, 1H), 3.61 – 3.50 (m, 2H), 3.30 – 3.13 (m, 2H), 2.33 (dt, *J* = 8.6, 5.1 Hz, 2H), 2.15 – 1.96 (m, 1H), 1.82 – 1.68 (m, 1H), 1.64 –

1.32 (m, 4H), 1.20 (s, 9H), 1.00 (d, $J = 6.3$ Hz, 3H), 0.84 (dd, $J = 9.0$, 6.1 Hz, 6H). $^{13}$C NMR (100 MHz, $CDCl_3$): δ 181.06, 172.62, 169.58, 156.06, 138.34, 134.40, 129.86, 128.27, 127.94, 125.94, 75.52, 66.85, 65.97, 65.85, 58.98, 52.37, 50.31, 41.29, 40.55, 38.29, 32.66, 28.47, 28.21, 24.90, 22.77, 22.14, 17.21.

**3-chlorobenzyl ((2S,3R)-3-(tert-butoxy)-1-(((S)-4-methyl-1-oxo-1-(((S)-1-oxo-3-((S)-2-oxopyrrolidin-3-yl)propan-2-yl)amino)pentan-2-yl)amino)-1-oxobutan-2-yl)carbamate (MPI126). MPI26** was prepared as a white solid following a similar procedure to **MPI11** (yield 79%). $^1$H NMR (400 MHz, Chloroform-*d*) δ 9.50 (s, 1H), 8.39 – 7.96 (m, 1H), 7.47 (d, $J = 7.6$ Hz, 1H), 7.35 (s, 1H), 7.29 (d, $J = 4.7$ Hz, 2H), 7.23 (s, 1H), 6.19 – 5.86 (m, 2H), 5.07 (q, $J = 12.6$ Hz, 2H), 4.55 – 4.34 (m, 2H), 4.18 (d, $J = 5.3$ Hz, 2H), 3.37 – 3.26 (m, 2H), 2.56 – 2.26 (m, 3H), 2.01 – 1.92 (m, 1H), 1.87 – 1.77 (m, 1H), 1.77 – 1.64 (m, 2H), 1.59 (t, $J = 9.0$ Hz, 1H), 1.27 (s, 9H), 1.08 (d, $J = 5.9$ Hz, 3H), 0.96 (dd, $J = 10.9$, 5.7 Hz, 6H). $^{13}$C NMR (100 MHz, $CDCl_3$) δ 199.64, 172.82, 169.54, 169.33, 155.98, 138.29, 134.42, 129.88, 128.31, 127.95, 125.96, 75.53, 66.74, 66.00, 58.91, 57.66, 52.11, 41.48, 40.61, 28.60, 28.22, 24.93, 22.92, 22.08, 17.21.

**(((3-chlorobenzyl)oxy)carbonyl)-L-valine (MPI27a). MPI27a** was prepared as a white solid following a similar procedure to **MPI125g** and followed by **MPI111c** protocol. $^1$H NMR (400 MHz, DMSO-$d_6$) δ 12.58 (s, 1H), 7.55 (d, $J = 8.6$ Hz, 1H), 7.47 – 7.25 (m, 4H), 5.18 – 4.98 (m, 2H), 3.87 (dd, $J = 8.6$, 5.8 Hz, 1H), 2.15 – 1.96 (m, 1H), 0.89 (t, $J = 6.5$ Hz, 6H).

**Methyl (S)-2-((S)-2-((((3-chlorobenzyl)oxy)carbonyl)amino)-3-methylbutanamido)-3-cyclohexylpropanoate (MPI127b). MPI27b** was prepared as a white solid following a similar procedure to **MPI11b** (yield 56%). $^1$H NMR (400 MHz, DMSO-$d_6$) δ 8.25 (d, $J = 7.5$ Hz, 1H), 7.48 – 7.22 (m, 5H), 5.11 – 4.95 (m, 2H), 4.33 (q, $J = 7.7$, 7.2 Hz, 1H), 3.88 (t, $J = 8.1$ Hz, 1H), 3.60 (s, 3H), 2.01 – 1.89 (m, 1H), 1.72 – 1.44 (m, 7H), 1.34 (s, 1H), 1.22 – 1.00 (m, 3H), 0.98 – 0.69 (m, 8H).

**(S)-2-((S)-2-((((3-chlorobenzyl)oxy)carbonyl)amino)-3-methylbutanamido)-3-cyclohexylpropanoic acid (MPI127c). MPI27c** was prepared as a white solid following a similar procedure to **MPI11c** (yield 83%).

**Methyl (5S,8S,11S)-1-(3-chlorophenyl)-8-(cyclohexylmethyl)-5-isopropyl-3,6,9-trioxo-11-(((S)-2-oxopyrrolidin-3-yl)methyl)-2-oxa-4,7,10-triazadodecan-12-oate (MPI127d). MPI27d** was prepared as a white solid following a similar procedure to **MPI11d** (yield 56%). $^1$H NMR (400 MHz, Chloroform-*d*) δ 7.81 (d, $J = 7.2$ Hz, 1H), 7.35 (s, 1H), 7.32 – 7.18 (m, 3H), 6.60 (d, $J = 8.4$ Hz, 1H), 6.02 (s, 1H), 5.42 (d, $J = 8.6$ Hz, 1H), 5.15 – 5.00 (m, 2H), 4.58 (td, $J = 8.8$, 5.5 Hz, 1H), 4.49 (d, $J = 6.6$ Hz, 1H), 3.99 (dd, $J = 8.6$, 6.2 Hz, 1H), 3.72 (s, 3H), 3.41 – 3.24 (m, 2H), 2.48 – 2.35 (m, 2H), 2.23 – 2.05 (m, 2H), 2.01 – 1.58 (m, 8H), 1.56 – 1.46 (m, 1H), 1.40 – 1.04 (m, 5H), 1.04 – 0.77 (m, 8H).

**3-chlorobenzyl ((S)-1-(((S)-3-cyclohexyl-1-oxo-1-(((S)-1-oxo-3-((S)-2-oxopyrrolidin-3-yl)propan-2-yl)amino)propan-2-yl)amino)-3-methyl-1-oxobutan-2-yl)carbamate (MPI127). MPI27** was prepared as a white solid following a similar procedure of **MPI11e** followed by **MPI11** procedure (yield 52%). $^1$H NMR (400 MHz,

Chloroform-*d*) δ 9.49 (s, 1H), 8.25 (d, *J* = 6.6 Hz, 1H), 7.39 – 7.11 (m, 4H), 7.02 (d, *J* = 8.3 Hz, 1H), 6.56 (s, 1H), 5.64 (d, *J* = 8.5 Hz, 1H), 5.06 (q, *J* = 12.6 Hz, 2H), 4.74 – 4.61 (m, 1H), 4.37 (s, 1H), 4.03 (t, *J* = 7.6 Hz, 1H), 3.41 – 3.22 (m, 2H), 2.50 – 2.27 (m, 2H), 2.19 – 2.07 (m, 1H), 2.07 – 1.86 (m, 3H), 1.86 – 1.48 (m, 8H), 1.39 – 1.04 (m, 5H), 1.04 – 0.81 (m, 8H). $^{13}$C NMR (100 MHz, $CDCl_3$) δ 199.5, 180.0, 173.3, 171.2, 156.4, 138.2, 134.4, 129.9, 128.3, 128.0, 126.0, 66.2, 60.6, 57.5, 51.1, 40.6, 40.2, 38.0, 34.2, 33.5, 32.5, 31.0, 29.9, 29.7, 28.4, 26.3, 26.2, 26.0, 19.2, 17.8.

**Methyl (S)-2-((S)-2-((S)-2-amino-3-methylbutanamido)-4-methylpentanamido)-3-((S)-2-oxopyrrolidin-3-yl)propanoate (MPI128f). MPI28f** was prepared as a white solid following a similar procedure to **MPI125f** (yield 90%).

**Methyl (S)-2-((S)-2-((S)-2-(1H-indole-2-carboxamido)-3-methylbutanamido)-4-methylpentanamido)-3-((S)-2-oxopyrrolidin-3-yl)propanoate (MPI128d). MPI28d** was prepared as a white solid following a similar procedure to **MPI11d** (yield 53%). $^{1}$H NMR (400 MHz, Methanol-$d_4$) δ 7.62 (d, *J* = 8.0 Hz, 1H), 7.49 – 7.39 (m, 1H), 7.22 (ddd, *J* = 8.2, 7.0, 1.2 Hz, 1H), 7.18 (d, *J* = 0.8 Hz, 1H), 7.11 – 7.01 (m, 1H), 4.53 (dd, *J* = 11.8, 3.9 Hz, 1H), 4.42 (dd, *J* = 13.8, 7.7 Hz, 2H), 3.72 (s, 3H), 2.56 (qd, *J* = 10.4, 4.0 Hz, 1H), 2.35 – 2.14 (m, 3H), 1.87 – 1.69 (m, 3H), 1.63 (t, *J* = 7.2 Hz, 2H), 1.04 (d, *J* = 6.7 Hz, 6H), 0.94 (dd, *J* = 19.4, 6.5 Hz, 6H).

**N-((S)-3-methyl-1-(((S)-4-methyl-1-oxo-1-(((S)-1-oxo-3-((S)-2-oxopyrrolidin-3-yl)propan-2-yl)amino)pentan-2-yl)amino)-1-oxobutan-2-yl)-1H-indole-2-carboxamide (MPI128). MPI28** was prepared as a white solid following a similar procedure of **MPI11e** followed by **MPI11** procedure (yield 60%). $^{1}$H NMR (400 MHz, DMSO-$d_6$) δ 11.59 (d, *J* = 2.2 Hz, 1H), 9.42 (s, 1H), 8.48 (d, *J* = 7.8 Hz, 1H), 8.23 (dd, *J* = 19.8, 8.1 Hz, 2H), 7.71 – 7.57 (m, 2H), 7.50 – 7.39 (m, 1H), 7.29 (d, *J* = 2.1 Hz, 1H), 7.18 (ddd, *J* = 8.2, 6.9, 1.2 Hz, 1H), 7.03 (ddd, *J* = 8.0, 6.9, 1.0 Hz, 1H), 4.44 – 4.32 (m, 2H), 4.24 (ddd, *J* = 11.6, 7.8, 3.9 Hz, 1H), 3.21 – 3.11 (m, 1H), 3.07 (td, *J* = 9.2, 7.0 Hz, 1H), 2.31 (qd, *J* = 10.3, 3.9 Hz, 1H), 2.13 (dt, *J* = 14.3, 7.9 Hz, 2H), 1.96 – 1.85 (m, 1H), 1.73 – 1.58 (m, 3H), 1.58 – 1.43 (m, 2H), 1.02 – 0.79 (m, 12H). $^{13}$C NMR (100 MHz, DMSO-$d_6$) δ 201.2, 178.8, 173.1, 171.5, 161.5, 137.0, 131.8, 127.5, 123.9, 122.0, 120.2, 112.7, 104.2, 58.8, 56.6, 51.6, 41.2, 37.6, 30.9, 29.8, 27.7, 24.7, 23.3, 22.3, 19.6, 19.2

**Table S1. Statistics of crystallographic analysis of M$^{Pro}$ in complexed with different inhibitors.**

| Protein/Ligand (PDI entry) | **MPI11** (7RVM) | **MPI12** (7RVN) | **MPI13** (7RVO) | **MPI14** (7RVP) |
|---|---|---|---|---|
| Data Collection | | | | |
| Space group | C121 | C121 | P1 | P1 |
| cell dimensions | | | | |
| *a*, *b*, *c* (Å) | 98.71, 81.35, 51.98 | 98.92, 81.18, 51.92 | 55.51, 60.54, 63.15 | 55.58, 60.80, 63.45 |
| α, β, γ (°) | 90.00, 115.46, 90.00 | 90.00, 115.15, 90.00 | 79.98, 68.24, 70.25 | 80.06, 68.16, 70.12 |
| Resolution (Å) | 46.93-1.95 (2.00-1.95) | 47.00-1.63 (1.66-1.63) | 44.34-1.80 (1.84-1.80) | 58.81-1.90 (1.94-1.90) |
| $R_{merge}$ | 9.2 (72.0) | 6.6 (77.6) | 20.8 (57.9) | 10.4 (145.9) |
| $I/\sigma I$ | 9.3 (1.5) | 12.5 (1.6) | 3.5 (1.6) | 5.9 (0.9) |
| Completeness (%) | 97.4 (95.8) | 95.8 (90.5) | 89.8 (90.6) | 95.5 (94.5) |
| Redundancy | 6.9 (6.9) | 5.9 (5.5) | 3.7 (3.9) | 3.6 (3.7) |
| Refinement | | | | |
| Resolution (Å) | 46.93-1.95 | 47.00-1.63 | 44.34-1.80 | 43.48-1.90 |
| No. Reflections | 26267 (2594) | 44233 (4238) | 59690 (6029) | 54436 (5351) |
| $R_{work}/R_{free}$ | 0.2055/0.2383 | 0.1972/0.2124 | 0.2444/0.2750 | 0.2368/0.2736 |
| No. atoms | | | | |
| Protein | 2400 | 2426 | 4798 | 4802 |
| Water | 230 | 183 | 246 | 173 |
| *B* factors | | | | |
| Protein | 23.652 | 40.122 | 31.900 | 44.602 |
| Water | 30.233 | 43.562 | 36.757 | 44.891 |
| R.m.s deviations | | | | |
| Bond lengths (Å) | 0.009 | 0.009 | 0.010 | 0.010 |
| Bond angles (°) | 1.67 | 1.38 | 2.38 | 1.41 |

| Protein/Ligand (PDI entry) | **MPI16** (7RVQ) | **MPI18** (7RVR) | **MPI19** (7RVS) | **MPI20** (7RVT) |
|---|---|---|---|---|
| Data Collection | | | | |
| Space group | P1 | P1 | C121 | P 1 |
| cell dimensions | | | | |
| *a*, *b*, *c* (Å) | 56.34, 62.85, 63.57 | 54.70, 61.66, 62.10 | 98.80, 81.80, 52.03 | 55.51, 60.70, 63.34 |
| α, β, γ (°) | 80.34, 68.56, 69.65 | 81.13, 69.31, 69.71 | 90.00, 115.5, 90.00 | 80.42, 68.42, 70.59 |
| Resolution (Å) | 24.91-2.47 (2.58-2.47) | 24.27-2.46 (2.56-2.46) | 24.41-1.85 (1.89-1.85) | 58.83-2.10 (2.16-2.10) |
| $R_{merge}$ | 41.3 (419.6) | 21.7 (231.3) | 6.9 (118.7) | 30.0 (54.2) |
| $I/\sigma I$ | 3.5 (0.3) | 8.0 (0.5) | 17.8 (1.6) | 3.8 (2.6) |
| Completeness (%) | 98.1 (84.8) | 98.4 (86.8) | 99.9 (100.0) | 95.8 (96.3) |
| Redundancy | 8.9 (5.1) | 13.1 (7.8) | 10.1 (7.5) | 2.9 (3.1) |
| Refinement | | | | |
| Resolution (Å) | 24.91-2.49 | 24.27-2.46 | 24.41-1.85 | 32.85 - 2.10 |
| No. Reflections | 13243 (1018) | 12359 (758) | 31512 (3128) | 40069 (4014) |
| $R_{work}/R_{free}$ | 0.2386/0.3222 | 0.2529/0.3231 | 0.2000/0.2265 | 0.2702/0.3076 |
| No. atoms | | | | |
| Protein | 2431 | 2401 | 2427 | 2433 |
| Water | 45 | 37 | 176 | 144 |
| *B* factors | | | | |
| Protein | 50.449 | 50.903 | 37.652 | 31.883 |
| Water | 45.343 | 46.359 | 41.474 | 34.449 |
| R.m.s deviations | | | | |
| Bond lengths (Å) | 0.010 | 0.010 | 0.009 | 0.009 |
| Bond angles (°) | 1.25 | 1.24 | 1.31 | 1.13 |

#### Table S1 – continued

| Protein/Ligand (PDI entry) | MPI21 (7RVU) | MPI22 (7RVV) | MPI23 (7RVW) | MPI24 (7RVX) |
|---|---|---|---|---|
| Data Collection | | | | |
| Space group | P1 | P1 | C1 | P1 |
| cell dimensions | | | | |
| *a*, *b*, *c* (Å) | 55.60, 60.93, 62.87 | 46.46, 52.26, 64.12 | 95.91, 81.31, 54.47 | 56.06, 61.15, 63.76 |
| α, β, γ (°) | 79.27, 68.84, 69.73 | 108.82, 97.3, 98.14 | 90.00, 116.98, 90.00 | 79.94, 68.33, 70.17 |
| Resolution (Å) | 57.76-2.50 (2.60-2.50) | 48.52-3.00 (3.18-3.00) | 24.27-1.85 (1.89-1.85) | 49.69-1.85 (1.89-1.85) |
| $R_{merge}$ | 47.1 (91.8) | 10.5 (170.0) | 13.6 (98.3) | 8.3 (51.4) |
| $I/\sigma I$ | 2.1 (0.8) | 6.1 (4.0) | 13.3 (2.1) | 7.6 (1.5) |
| Completeness (%) | 98.9 (98.7) | 93.2 (95.9) | 99.9 (100.0) | 89.1 (85.9) |
| Redundancy | 5.4 (5.6) | 2.8 (2.9) | 11.4 (9.1) | 3.8 (3.8) |
| Refinement | | | | |
| Resolution (Å) | 43.51-2.50 | 48.52-3.00 | 24.27-1.85 | 49.69-1.85 |
| No. Reflections | 12573 (1249) | 10372 (1108) | 31811 (3152) | 56153 (5444) |
| $R_{work}/R_{free}$ | 0.2515/0.3285 | 0.3414/0.4279 | 0.1898/0.2175 | 0.2353/0.2651 |
| No. atoms | | | | |
| Protein | 2378 | 2376 | 2401 | 4840 |
| Water | 43 | 0 | 223 | 229 |
| *B* factors | | | | |
| Protein | 29.165 | 22.608 | 30.173 | 40.269 |
| Water | 26.273 | 0 | 34.473 | 42.174 |
| R.m.s deviations | | | | |
| Bond lengths (Å) | 0.010 | 0.010 | 0.009 | 0.012 |
| Bond angles (°) | 1.28 | 1.38 | 1.24 | 1.50 |

| Protein/Ligand (PDI entry) | MPI25 (7RVY) | MPI26 (7RVZ) | MPI27 (7RV0) | MPI28 (7RV1) |
|---|---|---|---|---|
| Data Collection | | | | |
| Space group | C1 | C121 | C1 | P1 |
| cell dimensions | | | | |
| *a*, *b*, *c* (Å) | 96.50, 81.24, 54.48 | 98.33, 80.95, 51.82 | 95.66, 80.81, 54.44 | 55.64, 61.09, 63.87 |
| α, β, γ (°) | 90.00, 117.06, 90.00 | 90.00, 115.19, 90.00 | 90.00, 116.96, 90.00 | 79.27, 68.06, 69.37 |
| Resolution (Å) | 24.21-1.85 (1.89-1.85) | 59.88-1.90 (1.94-1.90) | 24.26-1.85 (1.89-1.85) | 59.13-2.50 (2.60-2.50) |
| $R_{merge}$ | 6.5 (38.7) | 10.1 (141.9) | 12.2 (90.8) | 26.2 (37.8) |
| $I/\sigma I$ | 12.2 (2.5) | 8.7 (1.3) | 8.8 (1.7) | 4.7 (2.8) |
| Completeness (%) | 92.6 (81.3) | 96.1 (95.6) | 99.8 (98.9) | 97.1 (97.4) |
| Redundancy | 4.2 (3.7) | 6.4 (6.5) | 6.8 (4.0) | 2.9 (3.1) |
| Refinement | | | | |
| Resolution (Å) | 24.21-1.85 | 43.45-1.90 | 24.26-1.85 | 46.10 - 2.50 |
| No. Reflections | 29597 (2613) | 27763 (2745) | 31429 (3102) | 24135 (2442) |
| $R_{work}/R_{free}$ | 0.2309/0.2680 | 0.1911/0.2199 | 0.2220/0.2534 | 0.2566/0.3160 |
| No. atoms | | | | |
| Protein | 2407 | 2358 | 2403 | 4722 |
| Water | 232 | 103 | 200 | 122 |
| *B* factors | | | | |
| Protein | 22.457 | 44.839 | 28.397 | 16.575 |
| Water | 25.228 | 44.799 | 31.384 | 15.514 |
| R.m.s deviations | | | | |
| Bond lengths (Å) | 0.009 | 0.008 | 0.011 | 0.010 |
| Bond angles (°) | 1.19 | 1.73 | 1.39 | 1.24 |

### NMR Spectroscopies of Synthesized Compounds

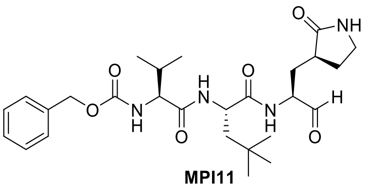

**MPI11**

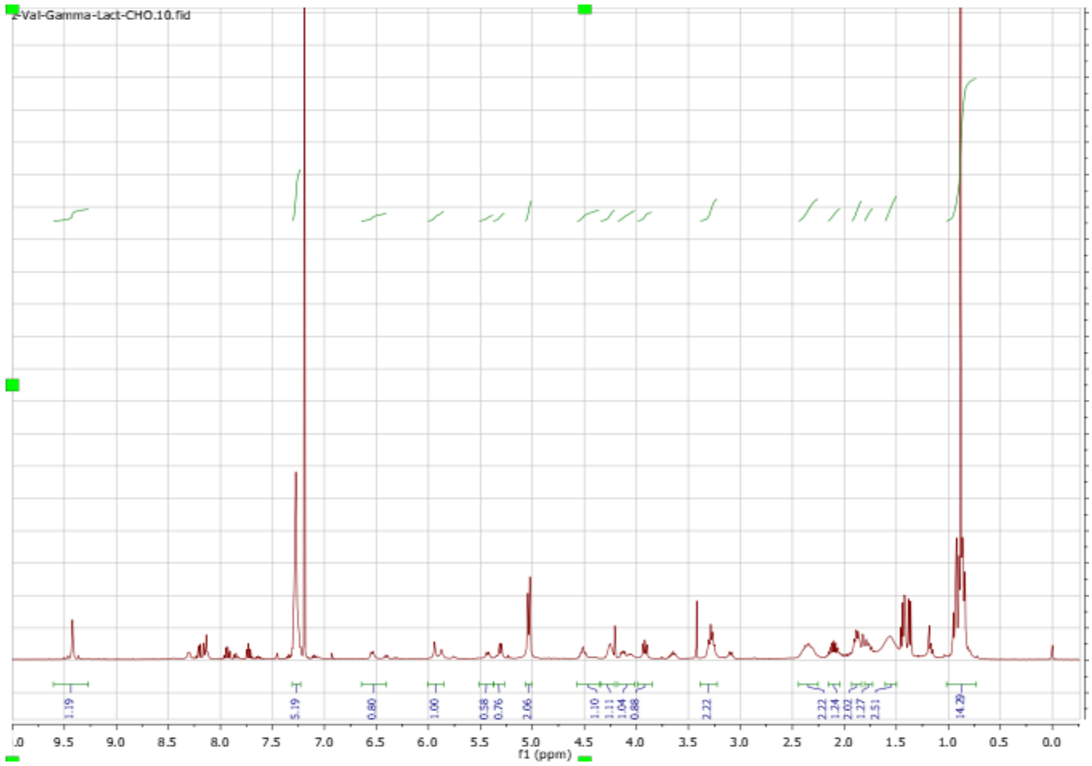

Z-Val-Gamma-Lact-CHO.10.fid
1.19
5.19
0.80
1.00
0.58
0.76
2.06
1.10
1.11
1.04
0.88
2.22
2.22
1.24
2.02
1.27
2.51
14.29
f1 (ppm)


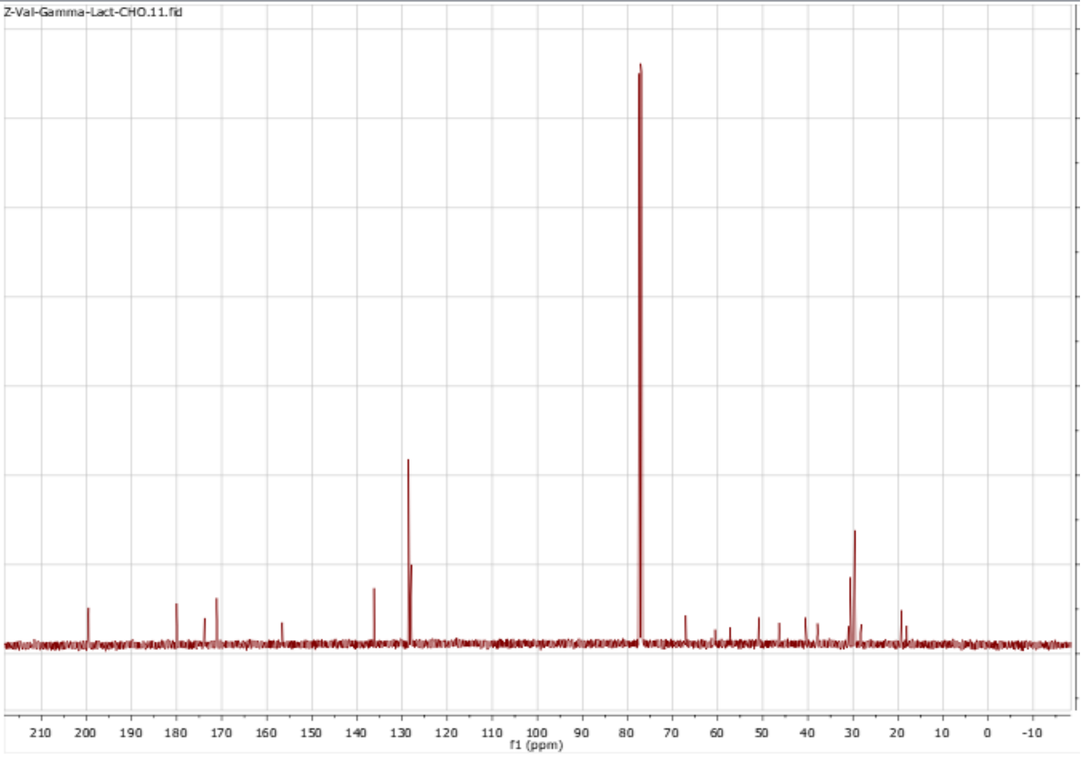

Z-Val-Gamma-Lact-CHO.11.fid
f1 (ppm)

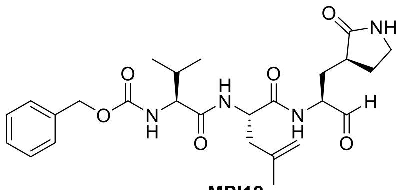

**MPI12**

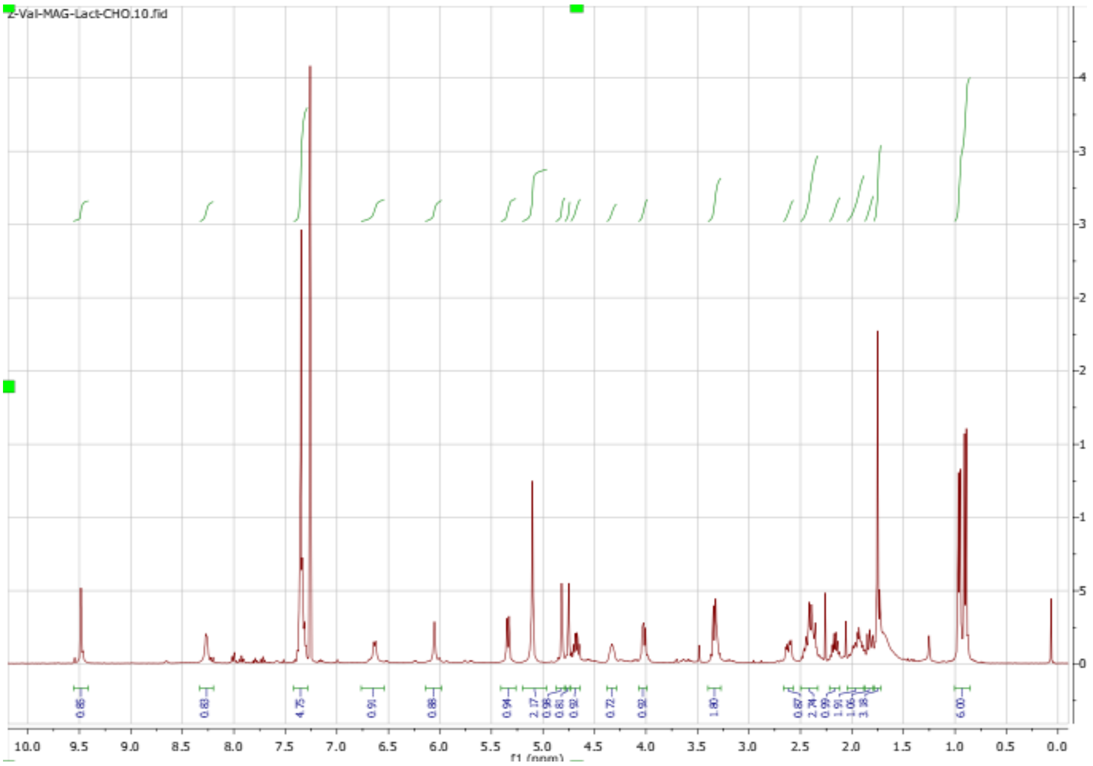


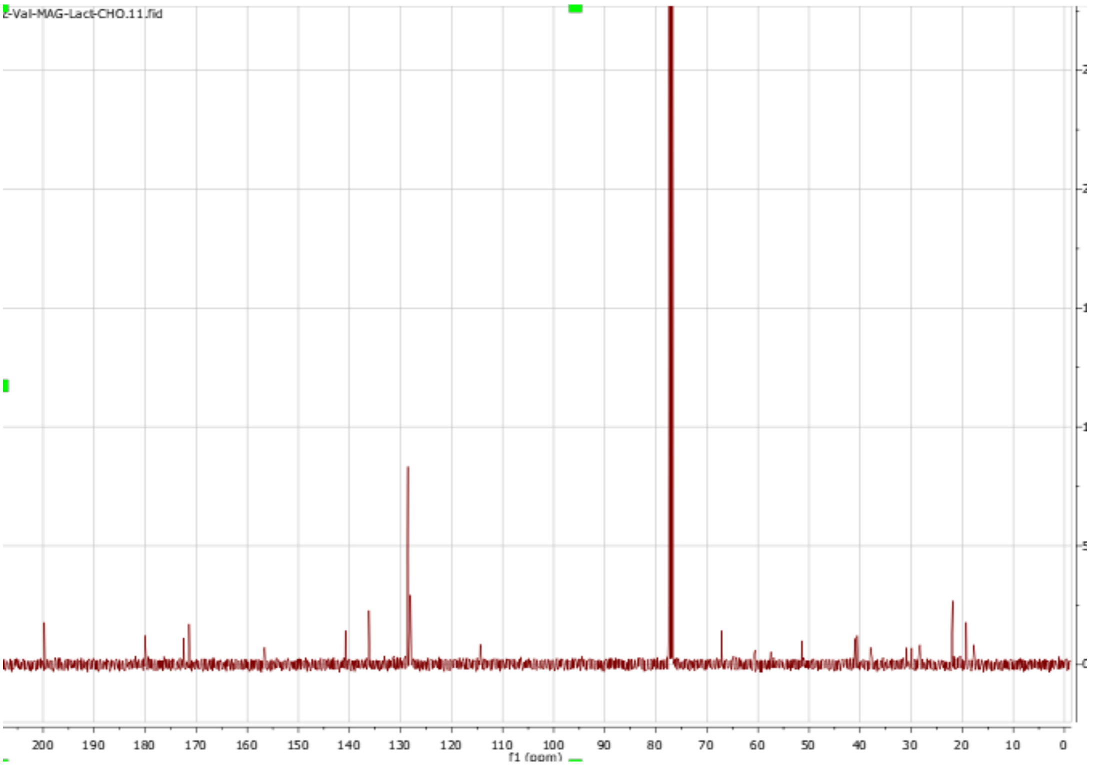

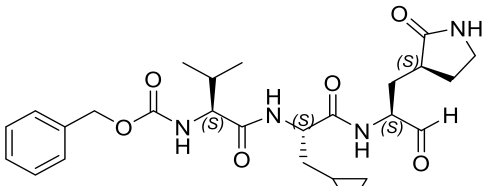
O
NH
(S)
O
H
O
N
H
(S)
N
H
(S)
O
O
N
H
(S)
H
O

**MPI13**

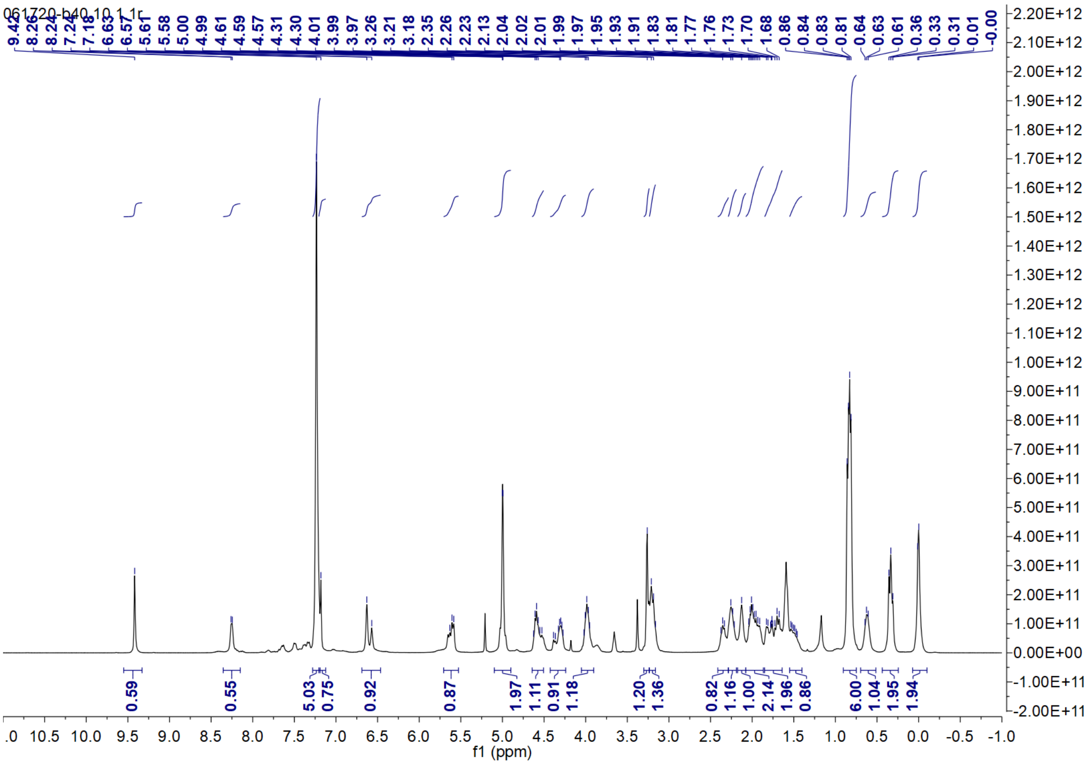
061720-b40.10.1.1r
9.42 8.26 8.24 7.24 7.18 6.63 6.57 5.61 5.58 5.00 4.99 4.61 4.59 4.57 4.31 4.30 4.01 3.99 3.97 3.26 3.21 3.18 2.35 2.26 2.23 2.13 2.04 2.02 2.01 1.99 1.97 1.95 1.93 1.91 1.83 1.81 1.77 1.76 1.73 1.70 1.68 0.86 0.84 0.83 0.81 0.64 0.63 0.61 0.36 0.33 0.31 0.01 -0.00
0.59 0.55 5.03 0.75 0.92 0.87 1.97 1.11 0.91 1.18 1.20 1.36 0.82 1.16 1.00 2.14 1.96 0.86 6.00 1.04 1.95 1.94
f1 (ppm)

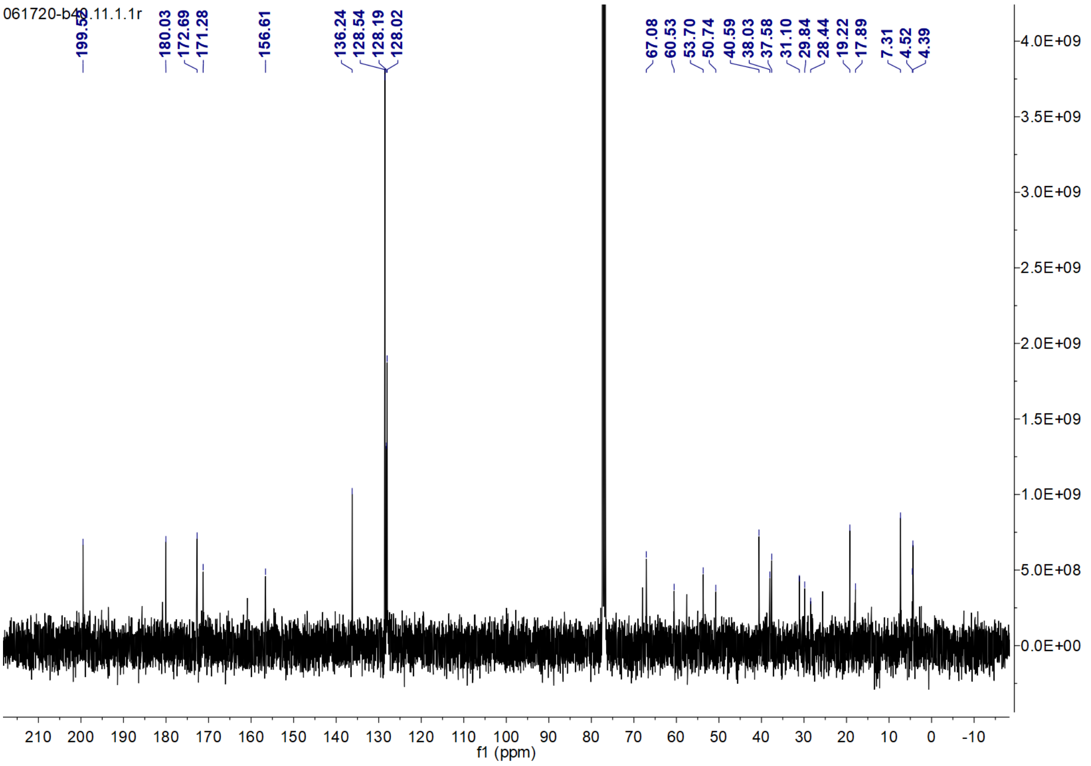
061720-b40.11.1.1r
199.52 180.03 172.69 171.28 156.61 136.24 128.54 128.19 128.02 67.08 60.53 53.70 50.74 40.59 38.03 37.58 31.10 29.84 28.44 19.22 17.89 7.31 4.52 4.39
f1 (ppm)

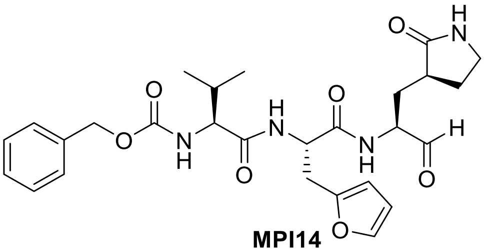

**MPI14**

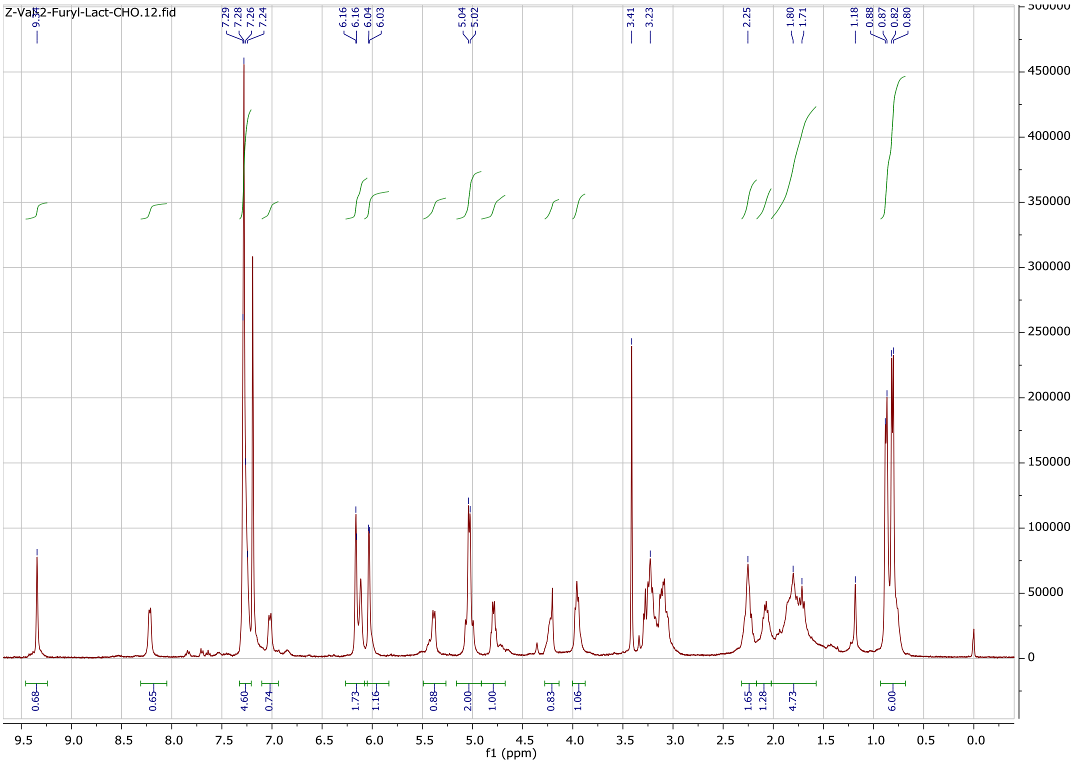
Z-Val-2-Furyl-Lact-CHO.12.fid
f1 (ppm)

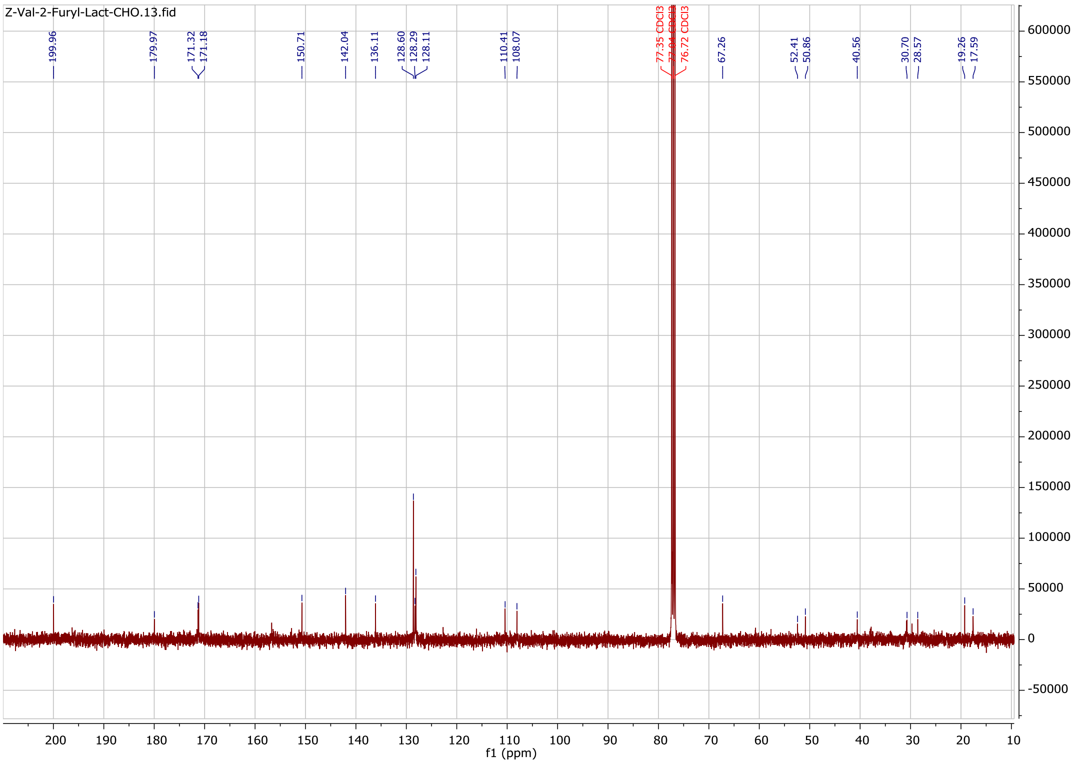
Z-Val-2-Furyl-Lact-CHO.13.fid
199.96
179.97
171.32
171.18
150.71
142.04
136.11
128.60
128.29
128.11
110.41
108.07
77.35 CDCl3
76.72 CDCl3
67.26
52.41
50.86
40.56
30.70
28.57
19.26
17.59
f1 (ppm)

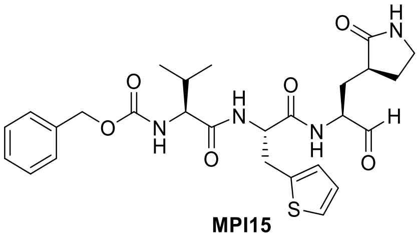

**MPI15**

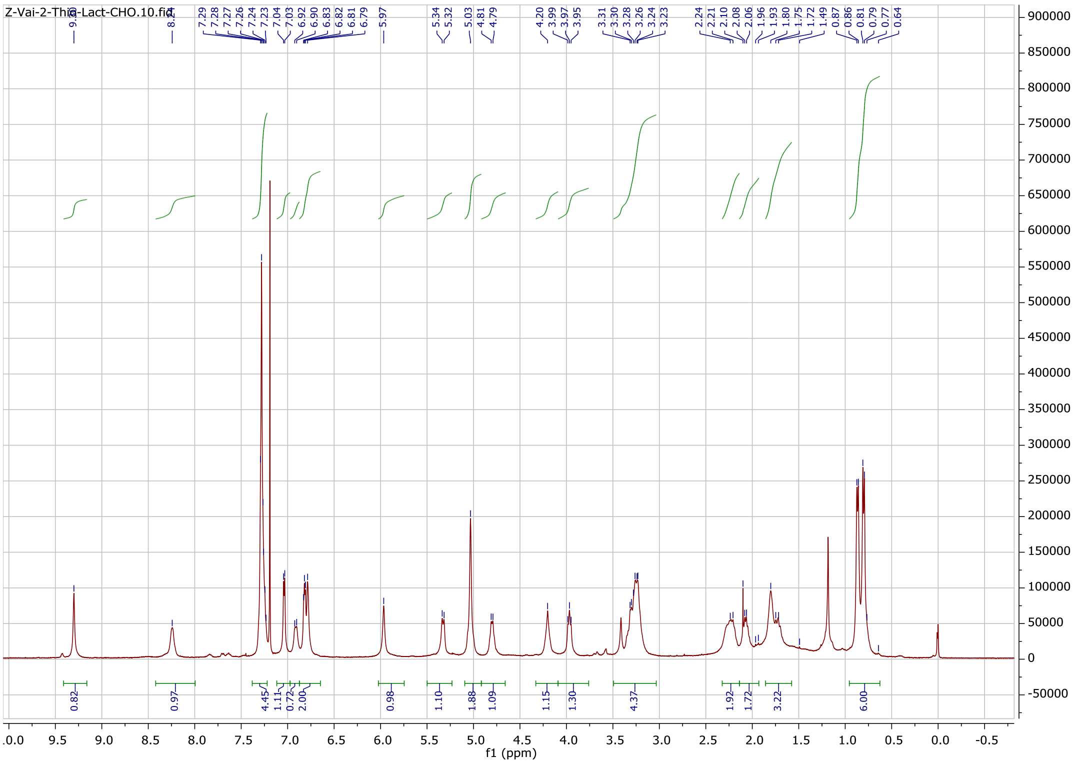

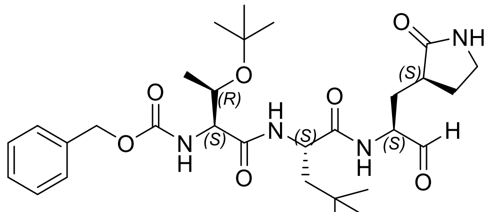

**MPI16**

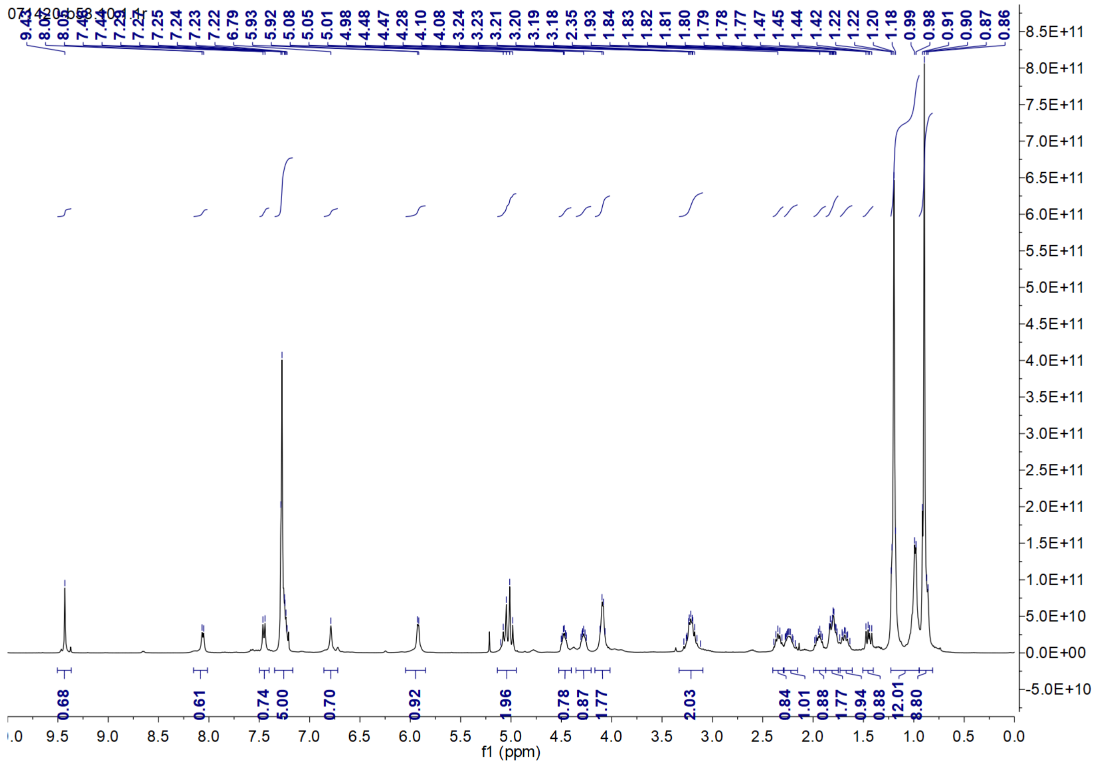

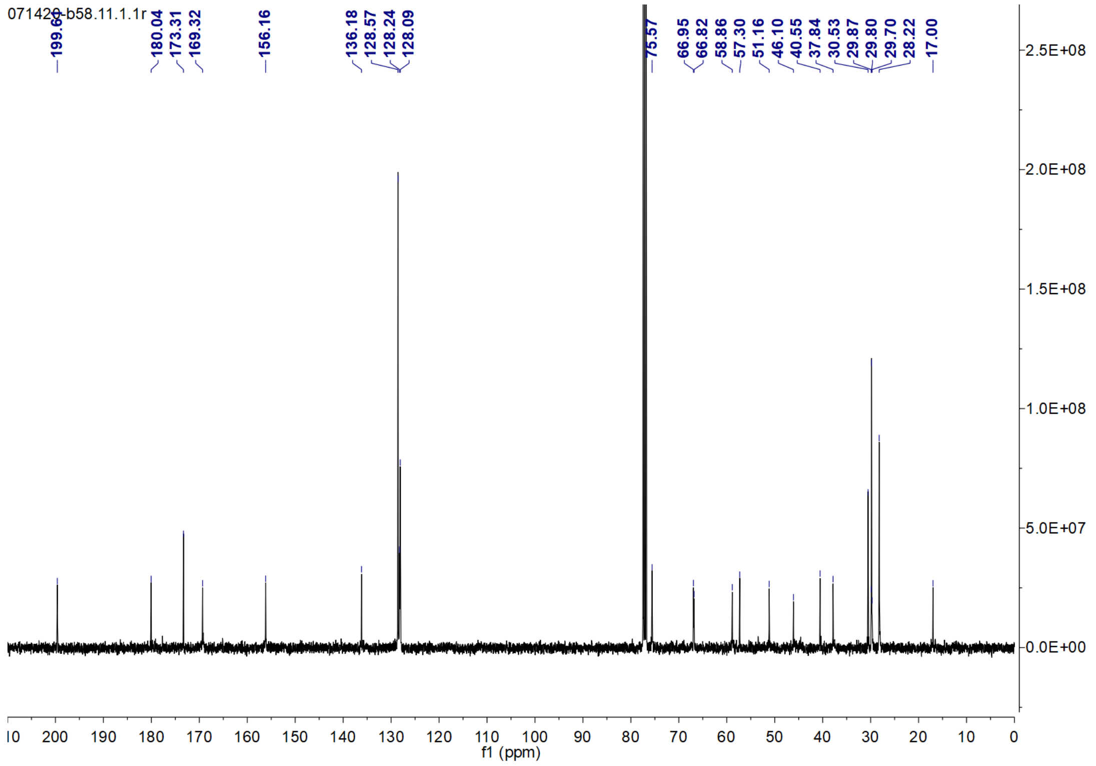

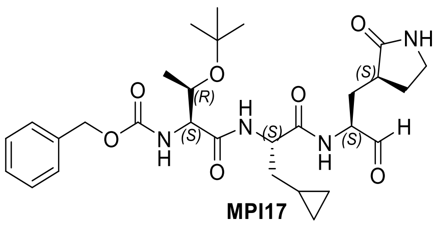

**MPI17**

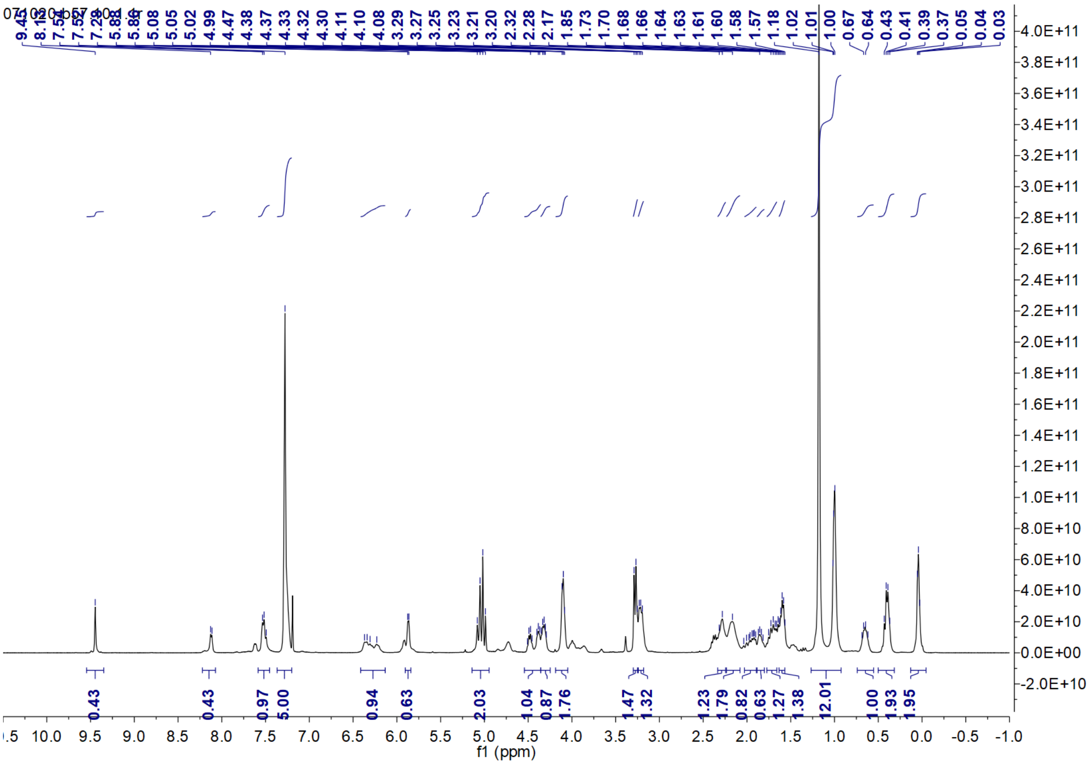

7.54
7.52
7.29
5.88
5.86
5.08
5.05
5.02
4.99
4.47
4.38
4.37
4.33
4.32
4.30
4.11
4.10
4.08
3.29
3.27
3.25
3.23
3.21
3.20
2.32
2.28
2.17
1.85
1.73
1.70
1.68
1.66
1.64
1.63
1.61
1.60
1.58
1.57
1.18
1.02
1.01
1.00
0.67
0.64
0.43
0.41
0.39
0.37
0.05
0.04
0.03
0.43
0.43
0.97
5.00
0.94
0.63
2.03
1.04
0.87
1.76
1.47
1.32
1.23
1.79
0.82
0.63
1.27
1.38
12.01
1.00
1.93
1.95
f1 (ppm)


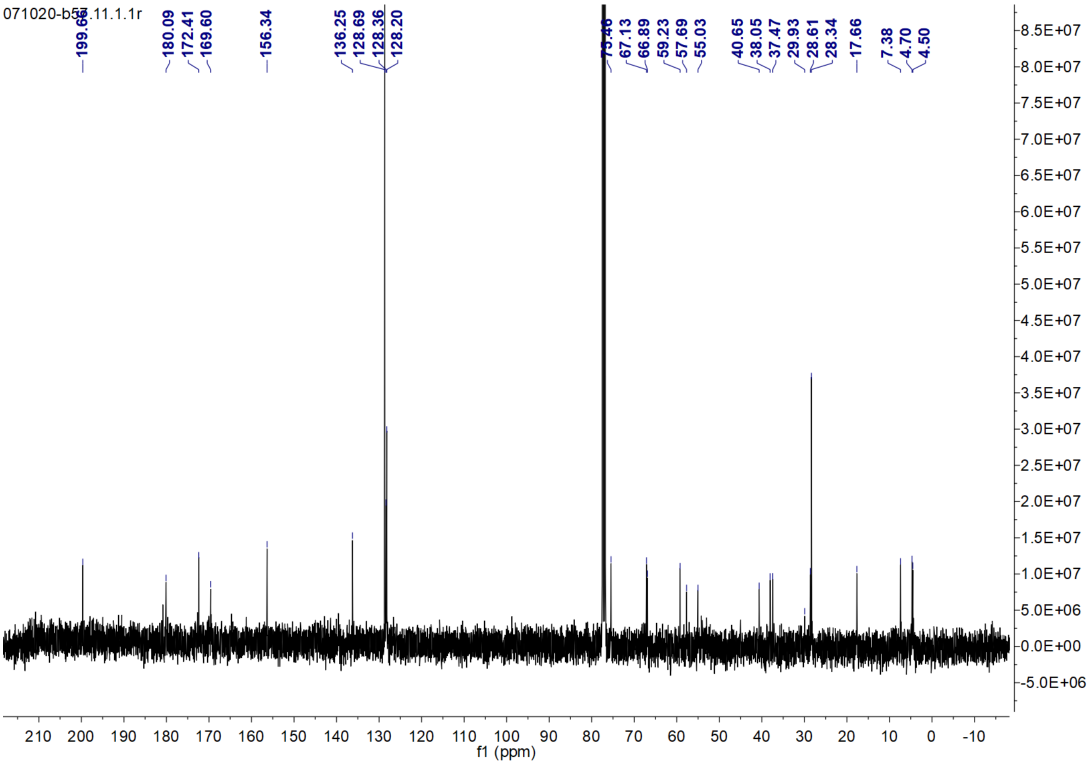

180.09
172.41
169.60
156.34
136.25
128.69
128.36
128.20
67.13
66.89
59.23
57.69
55.03
40.65
38.05
37.47
29.93
28.61
28.34
17.66
7.38
4.70
4.50
f1 (ppm)

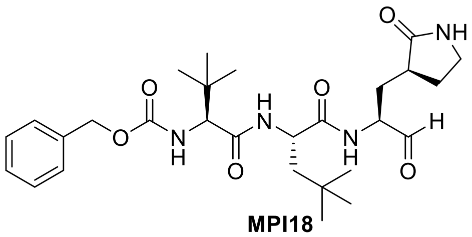
O
NH
O
O
N
H
H
N
O
N
H
H
O

**MPI18**

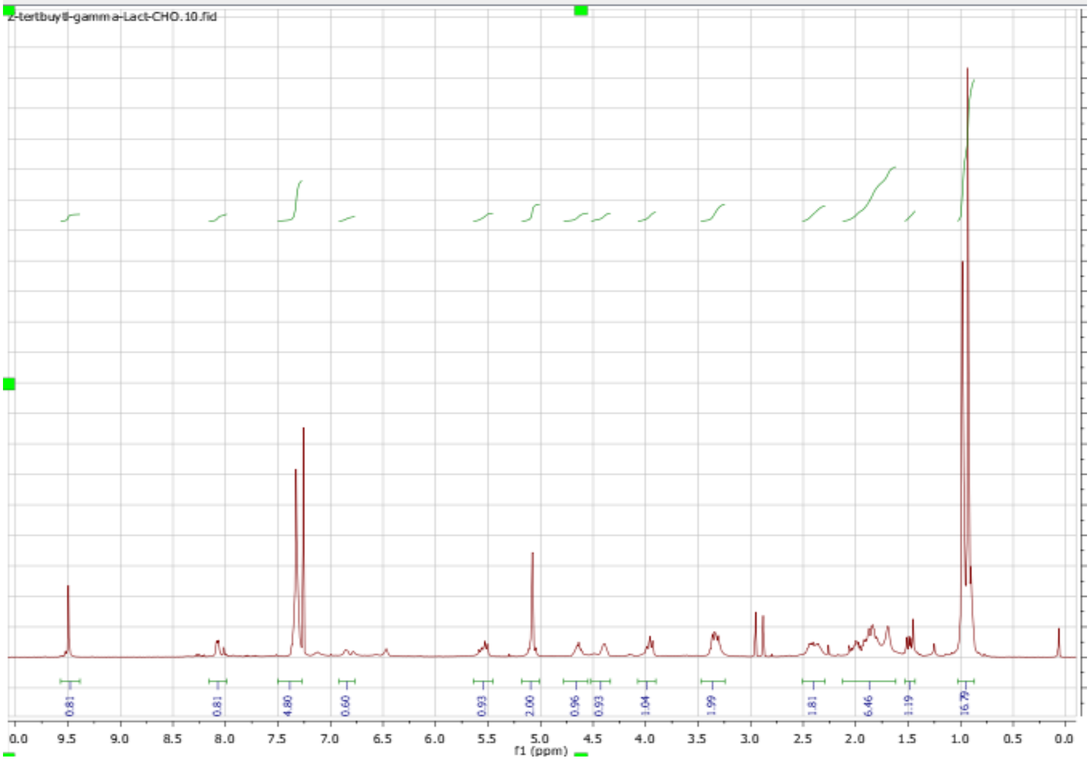
Z-tertbutyll-gamma-Lact-CHO.10.fid
0.81
0.81
4.80
0.60
0.93
2.00
0.96
0.93
1.04
1.99
1.81
6.46
1.19
16.79
f1 (ppm)

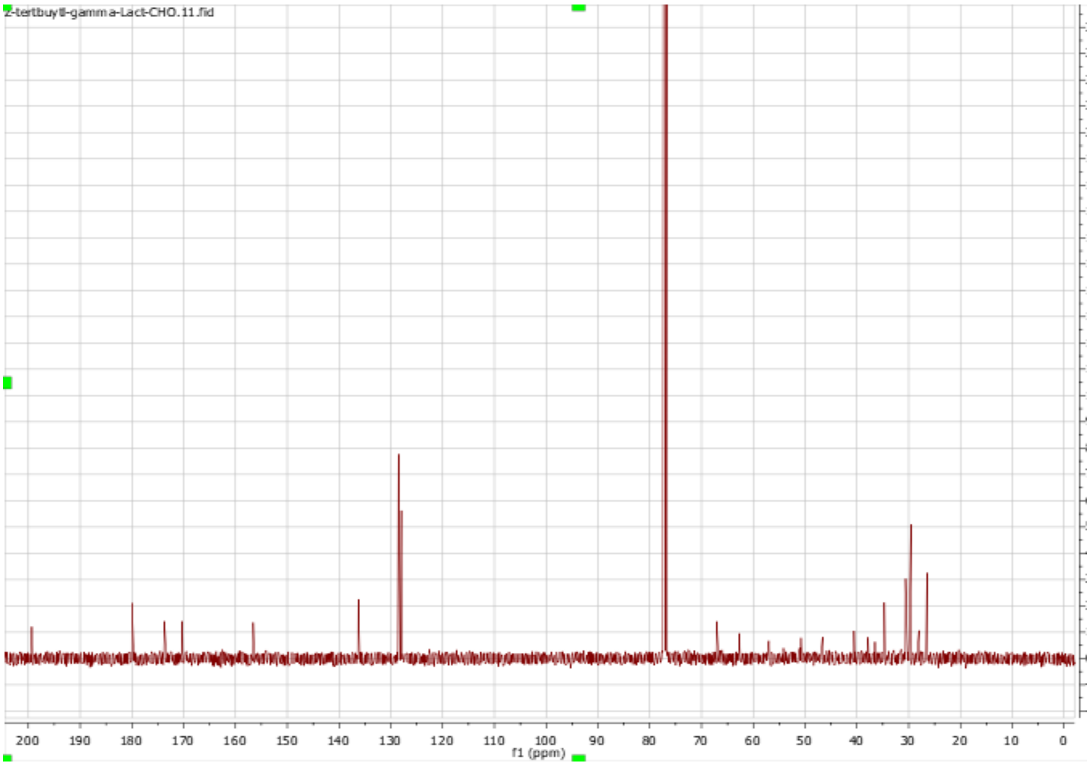
Z-tertbutyll-gamma-Lact-CHO.11.fid
f1 (ppm)

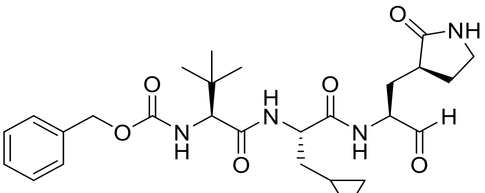

**MPI19**

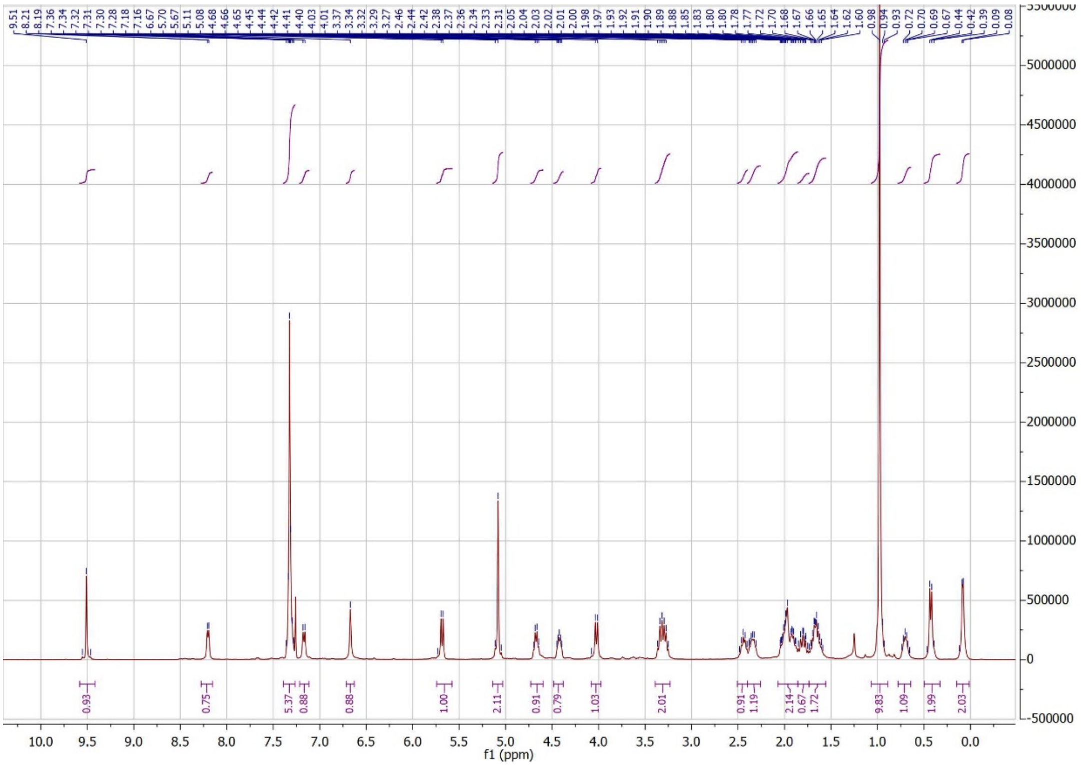
9.51
8.21
8.19
7.36
7.34
7.32
7.31
7.30
7.28
7.18
7.16
6.67
5.70
5.67
5.11
5.08
4.68
4.66
4.65
4.45
4.44
4.42
4.41
4.40
4.03
4.01
3.37
3.34
3.32
3.29
3.27
2.46
2.44
2.42
2.38
2.37
2.36
2.34
2.33
2.31
2.05
2.04
2.03
2.02
2.01
2.00
1.98
1.97
1.93
1.92
1.91
1.90
1.89
1.88
1.85
1.83
1.80
1.80
1.78
1.77
1.72
1.70
1.68
1.67
1.66
1.65
1.64
1.62
1.60
0.98
0.94
0.93
0.72
0.70
0.69
0.67
0.44
0.42
0.39
0.09
0.08
0.93
0.75
5.37
0.88
0.88
1.00
2.11
0.91
0.79
1.03
2.01
0.91
1.19
2.14
0.67
1.72
9.83
1.09
1.99
2.03
f1 (ppm)

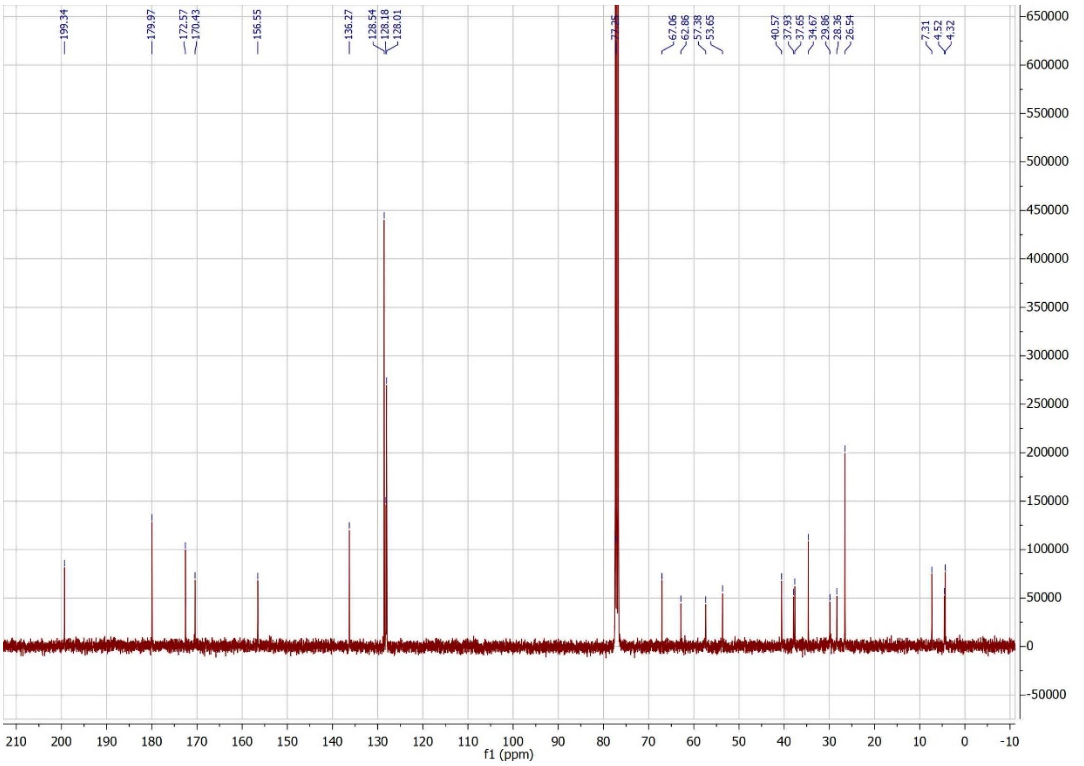
199.34
179.97
172.57
170.43
156.55
136.27
128.54
128.18
128.01
67.06
62.86
57.38
53.65
40.57
37.93
37.65
34.67
29.86
28.36
26.54
7.31
4.52
4.32
f1 (ppm)

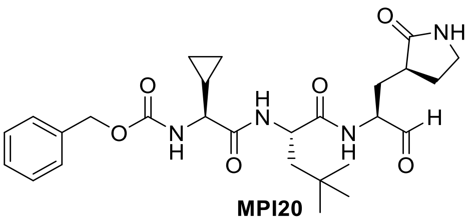

**MPI20**

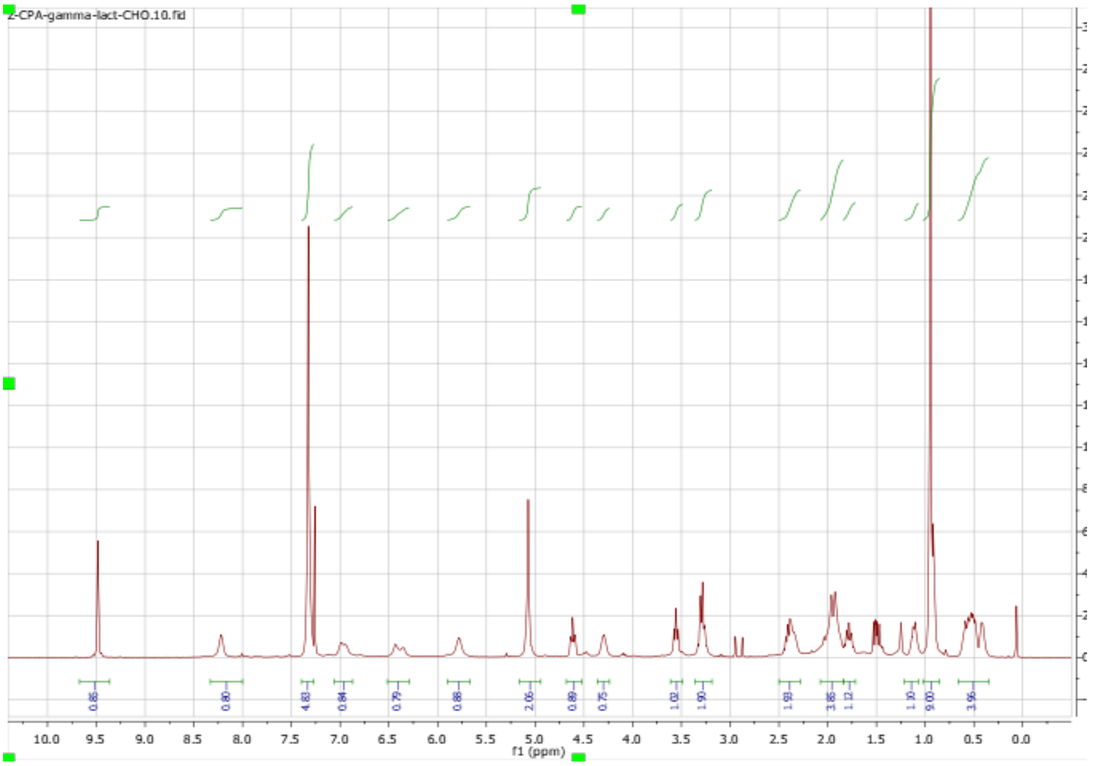

Z-CPA-gamma-lact-CHO.10.fid
0.86 0.80 4.83 0.84 0.79 0.88 2.06 0.89 0.75 1.02 1.90 1.93 3.86 1.12 1.30 9.00 3.96
10.0 9.5 9.0 8.5 8.0 7.5 7.0 6.5 6.0 5.5 5.0 4.5 4.0 3.5 3.0 2.5 2.0 1.5 1.0 0.5 0.0
f1 (ppm)


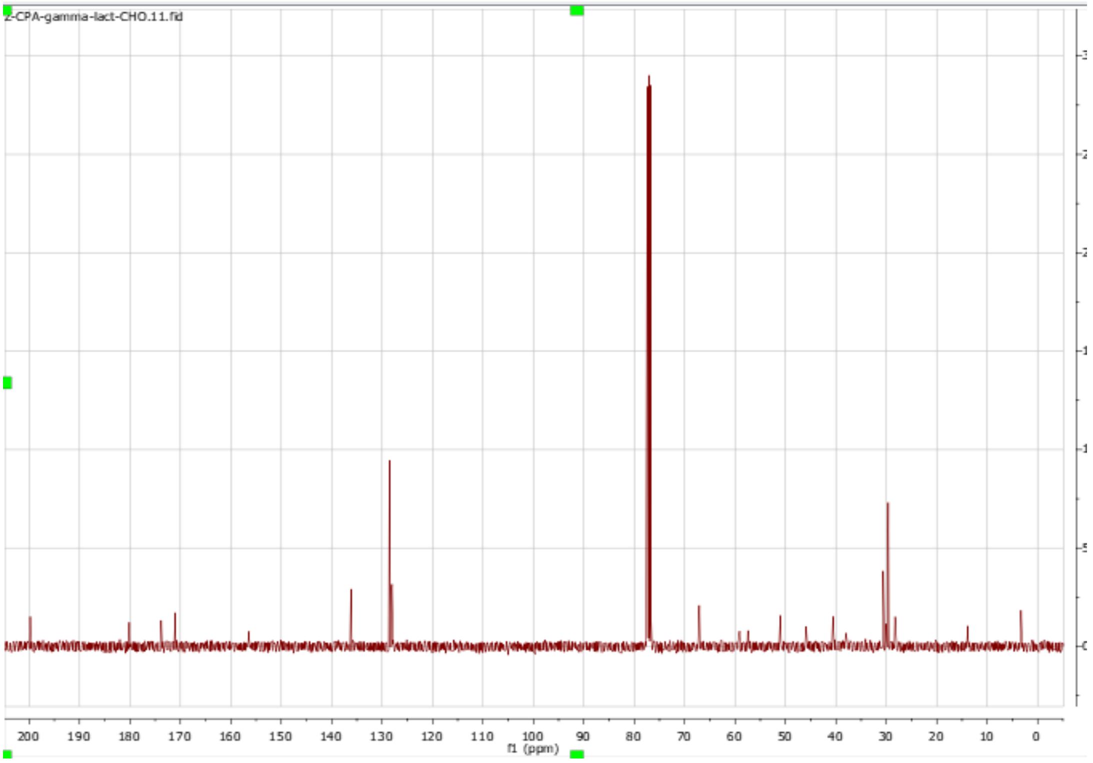

Z-CPA-gamma-lact-CHO.11.fid
200 190 180 170 160 150 140 130 120 110 100 90 80 70 60 50 40 30 20 10 0
f1 (ppm)

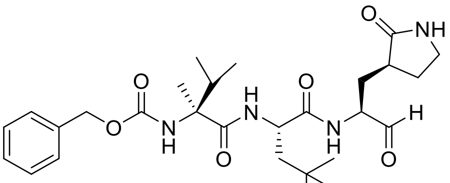

**MPI21**

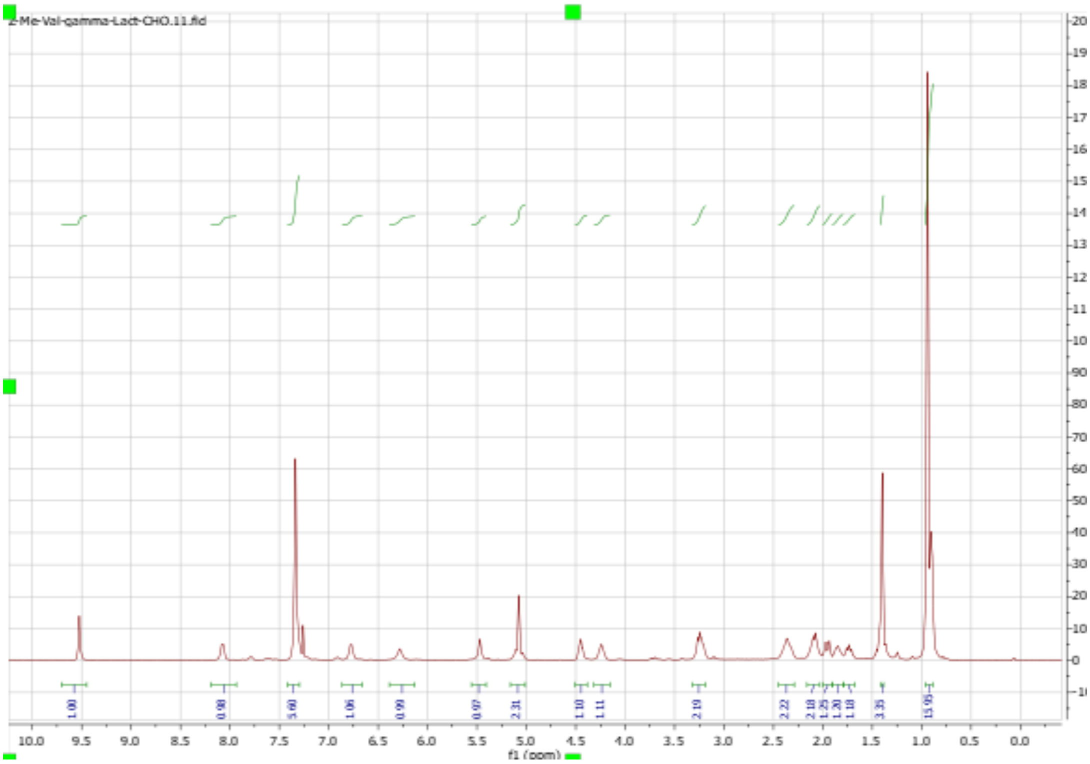
Z-Me-Val-gamma-Lact-CHO.11.fid
f1 (ppm)

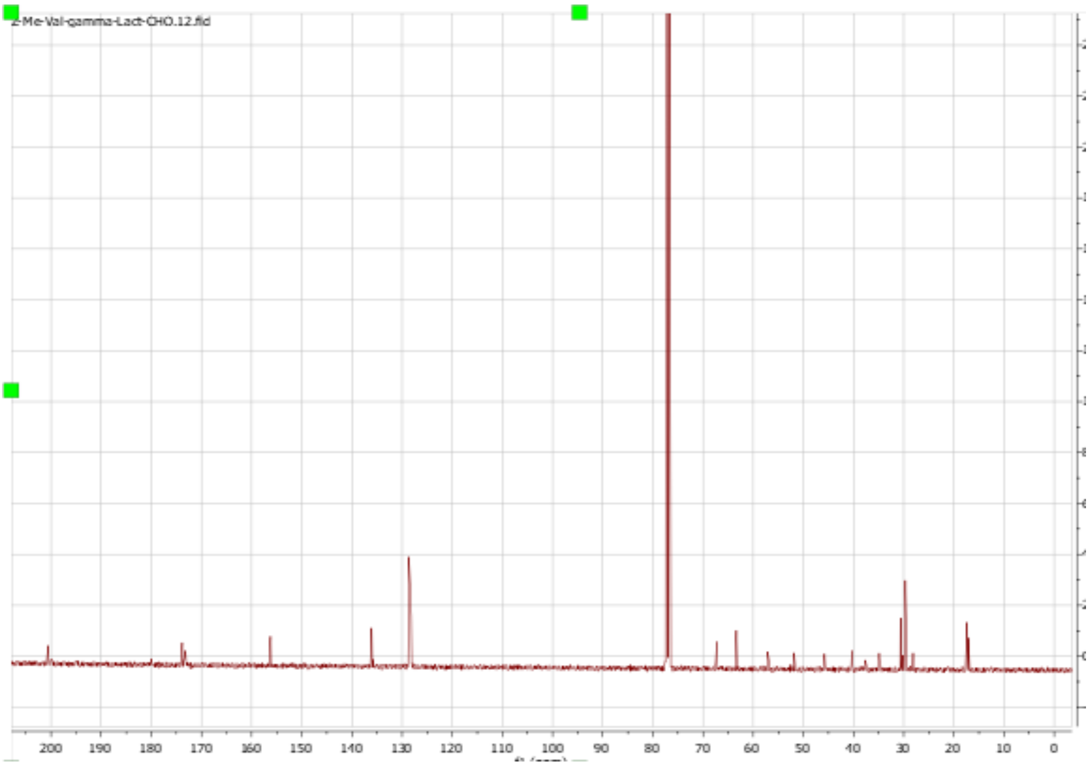
Z-Me-Val-gamma-Lact-CHO.12.fid

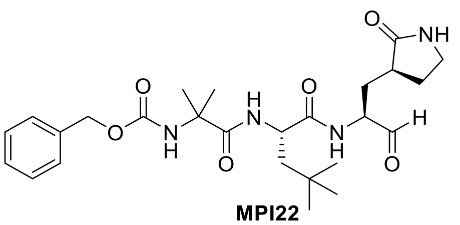

**MPI22**

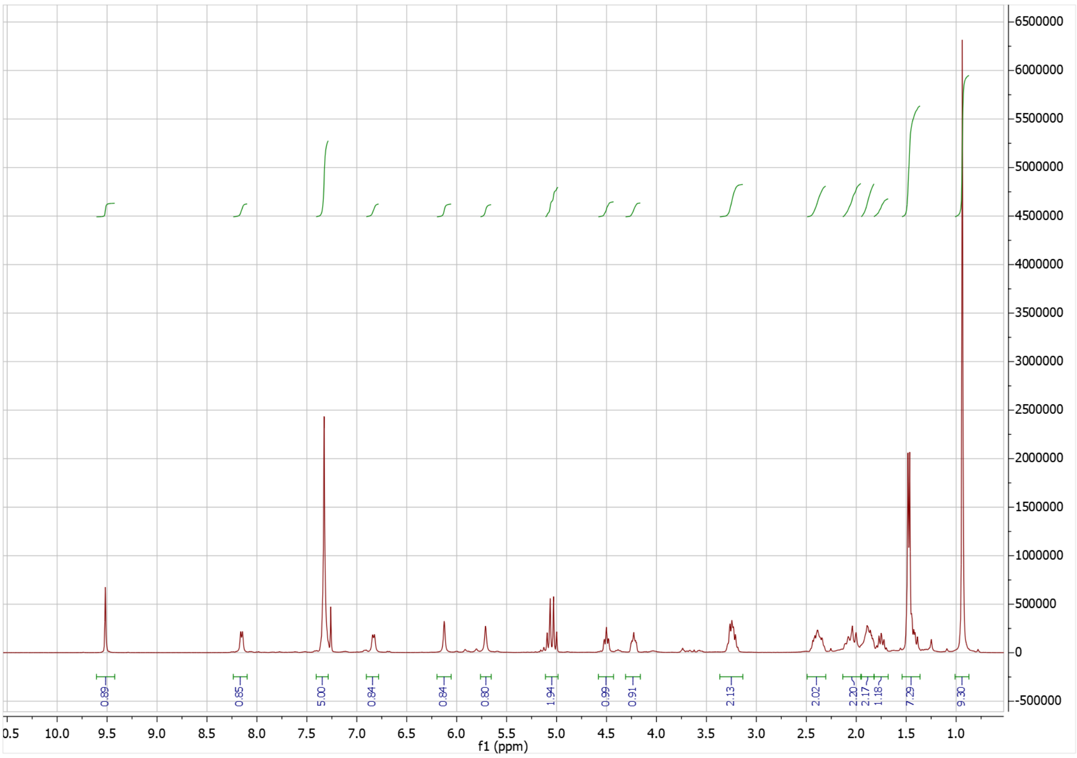

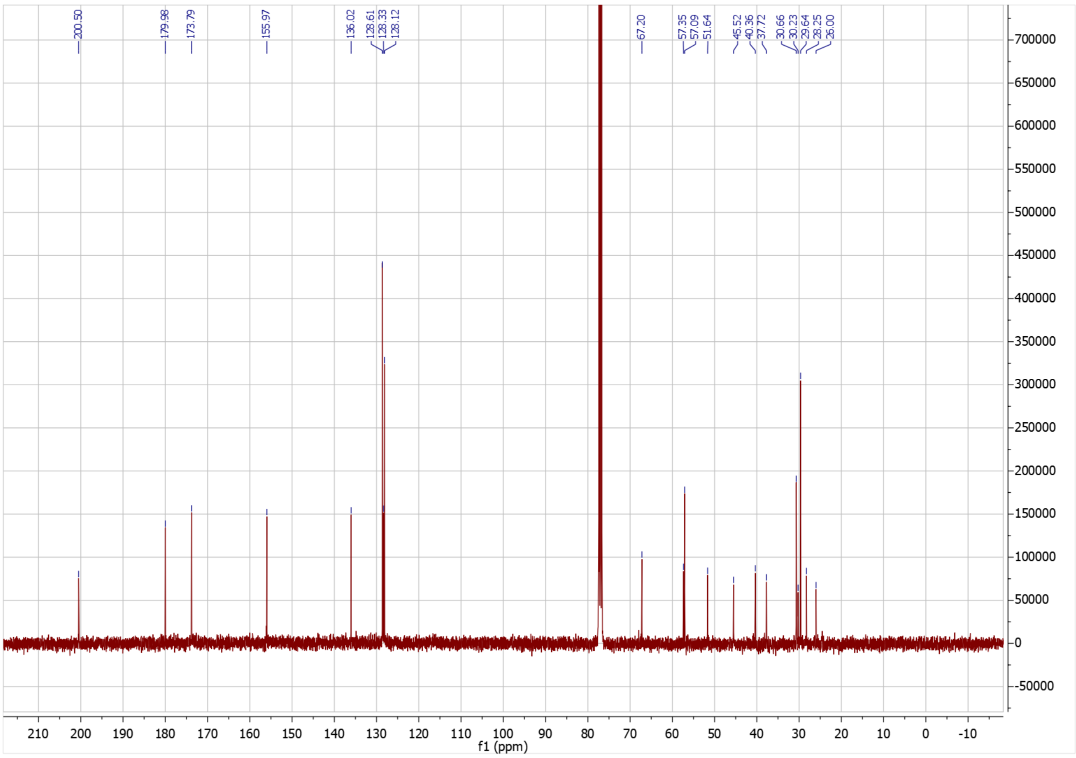

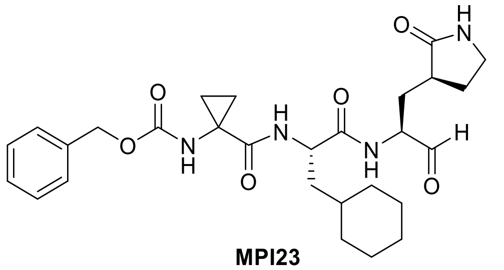

**MPI23**

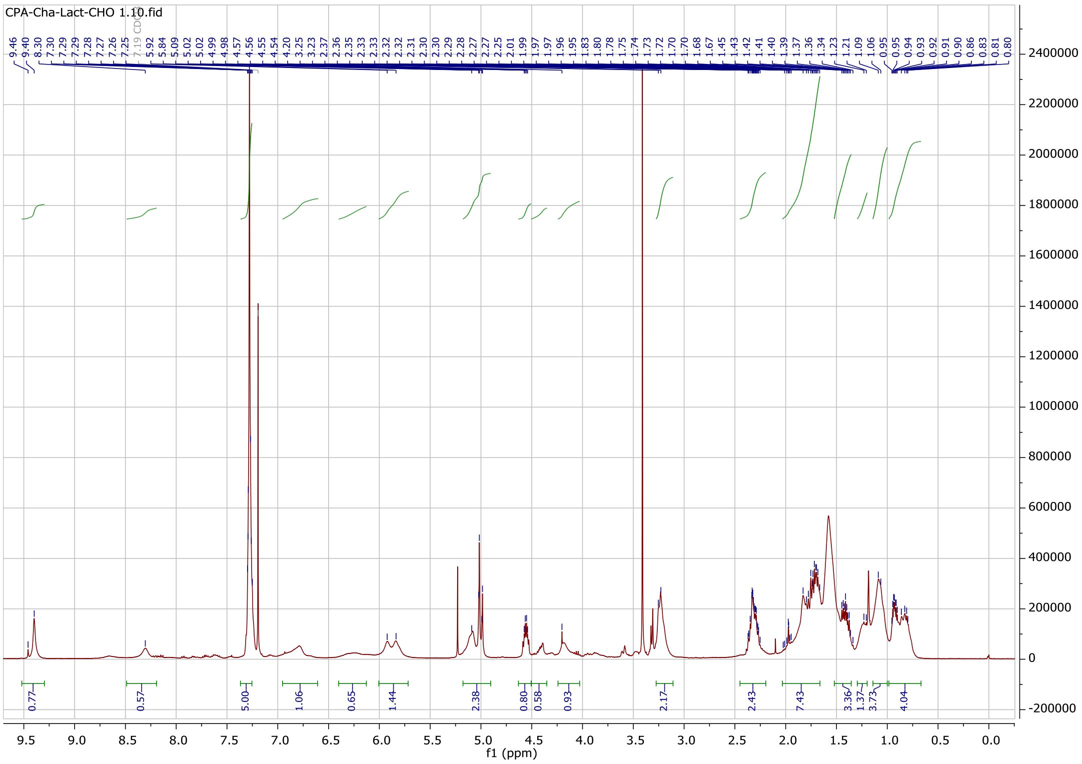

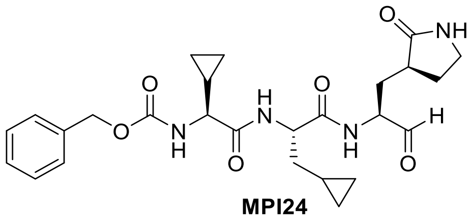

**MPI24**

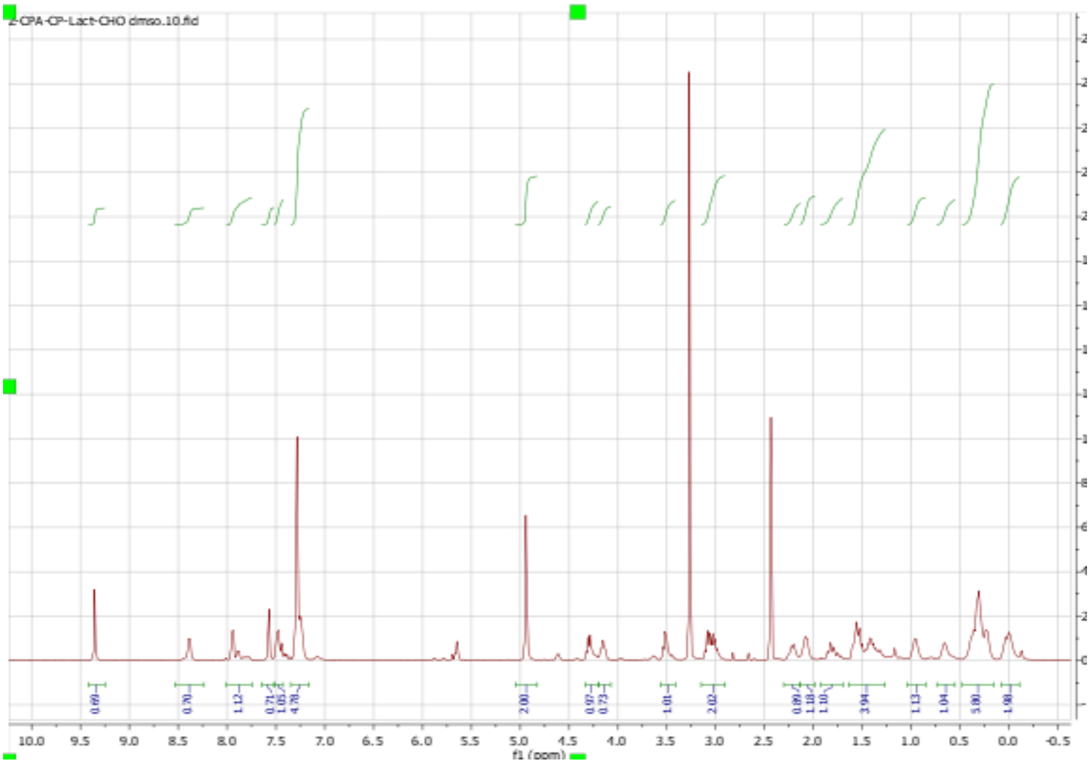
Z-CPA-CP-Lact-CHO dmso.10.fid
f1 (ppm)

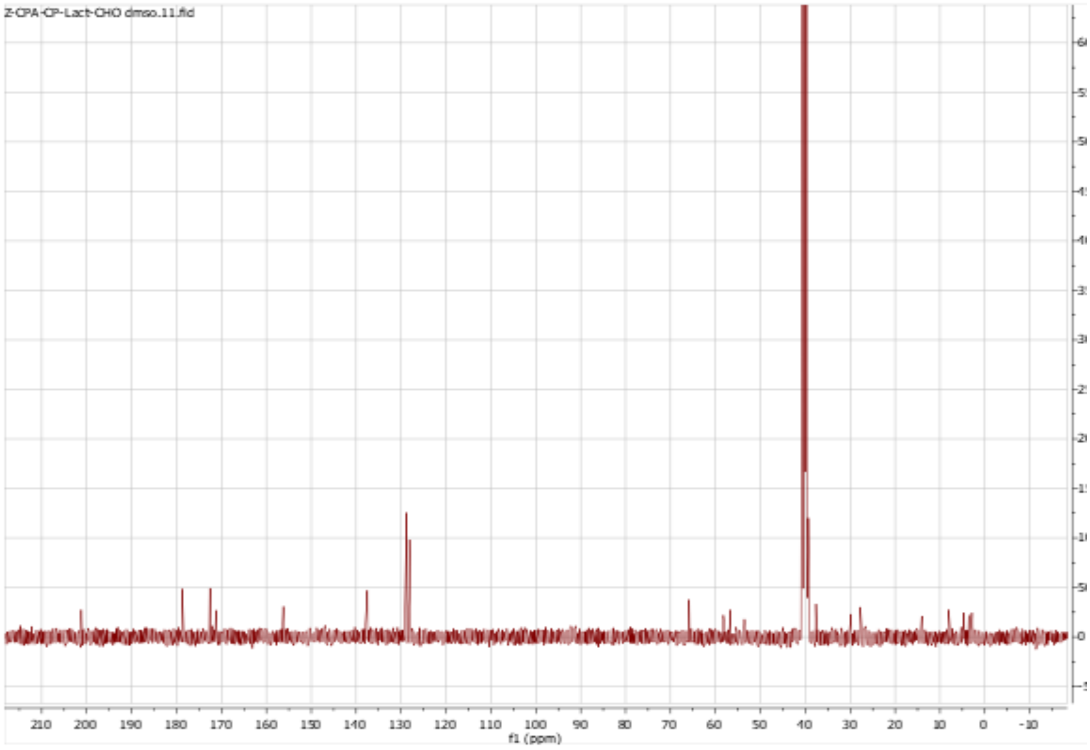
Z-CPA-CP-Lact-CHO dmso.11.fid
f1 (ppm)

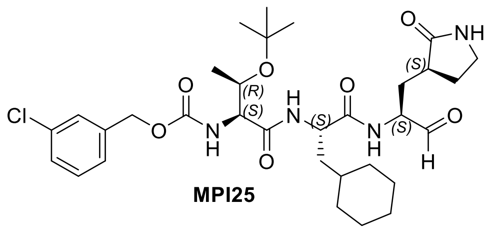
Cl
O
NH
(R)
(S)
(S)
(S)
H

**MPI25**

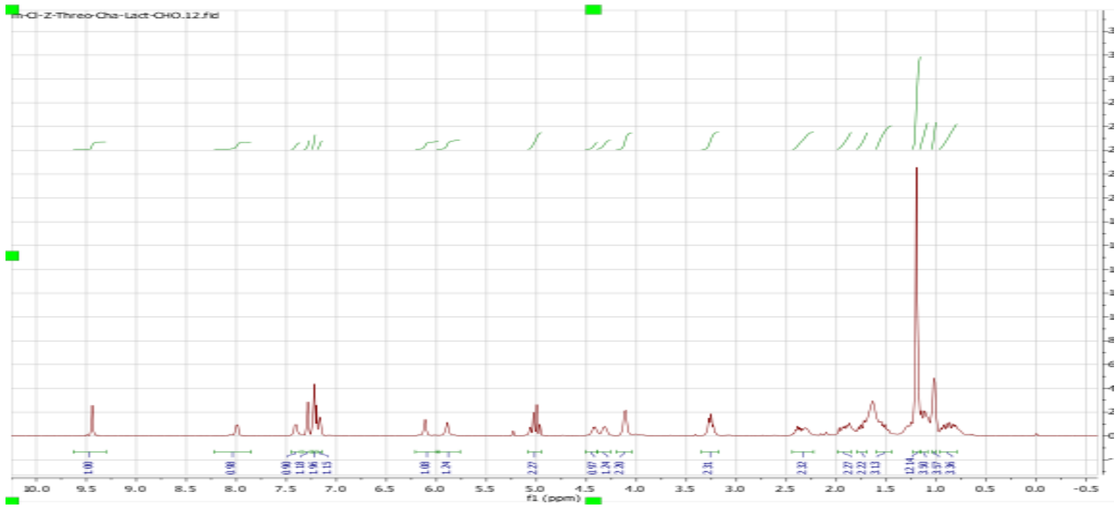
m-Cl-Z-Threo-Cha-Lact-CHO.12.fid
f1 (ppm)

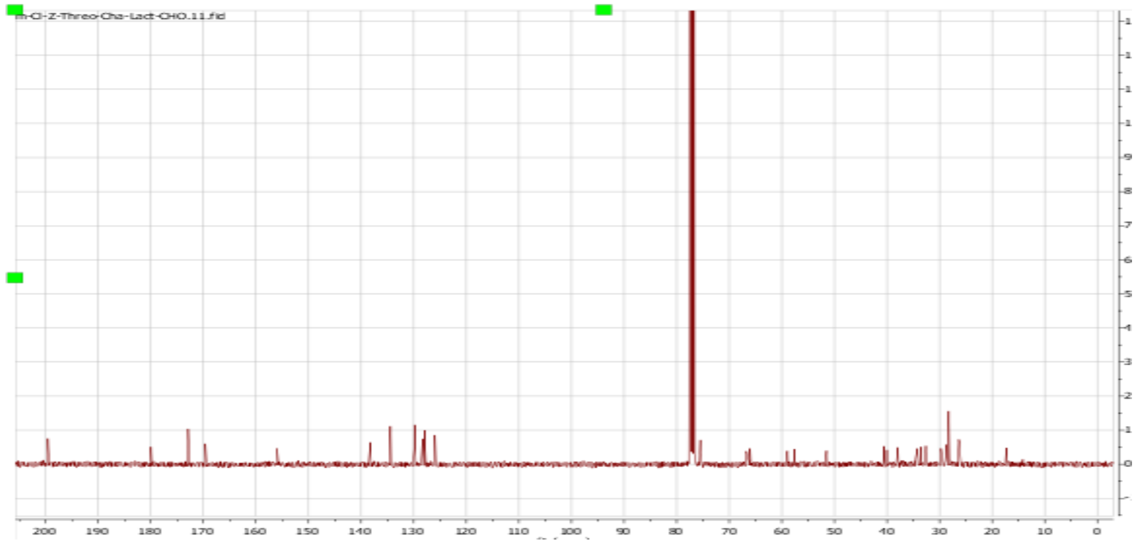
m-Cl-Z-Threo-Cha-Lact-CHO.11.fid

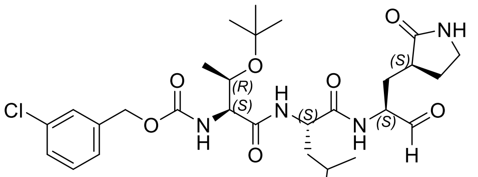
Cl
O
N
H
(R)
(S)
NH


**MPI26**

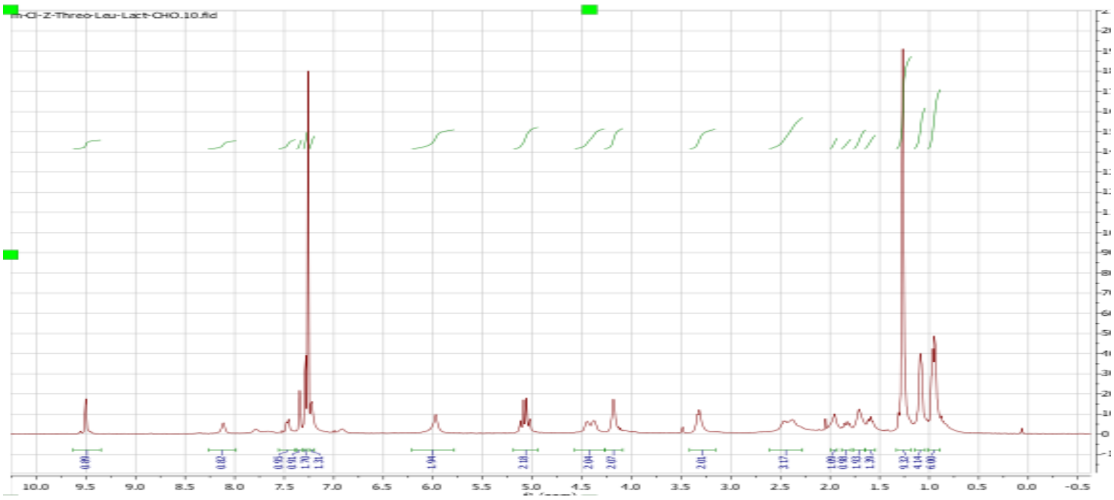
m-Cl-Z-Threo-Leu-Lact-CHO.10.fid
10.0 9.5 9.0 8.5 8.0 7.5 7.0 6.5 6.0 5.5 5.0 4.5 4.0 3.5 3.0 2.5 2.0 1.5 1.0 0.5 0.0 -0.5


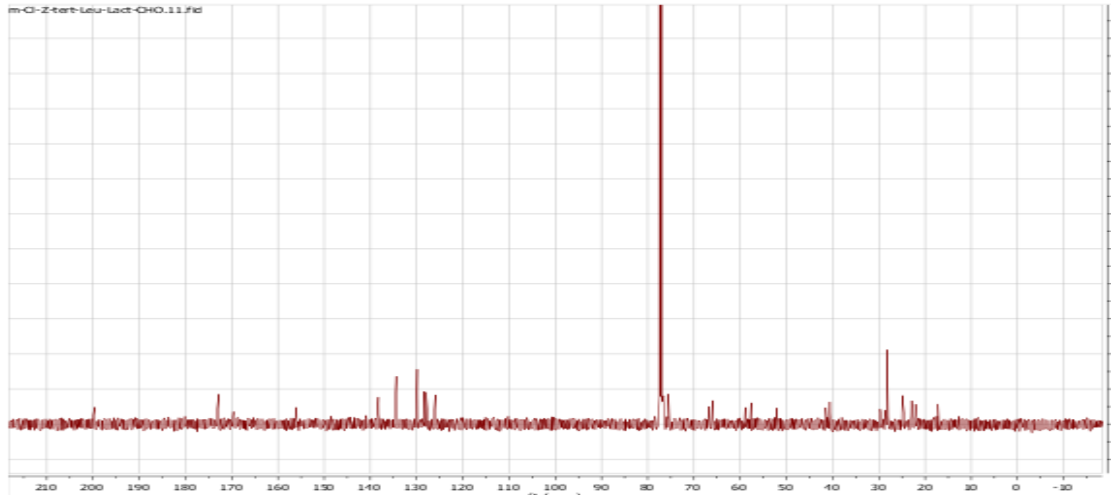
m-Cl-Z-tert-Leu-Lact-CHO.11.fid
210 200 190 180 170 160 150 140 130 120 110 100 90 80 70 60 50 40 30 20 10 0 -10

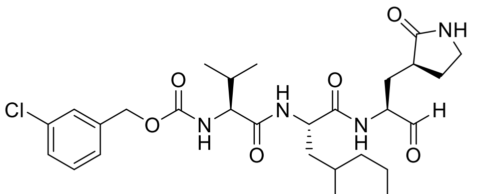

Cl
O
N
H
O
N
H
NH


**MPI27**

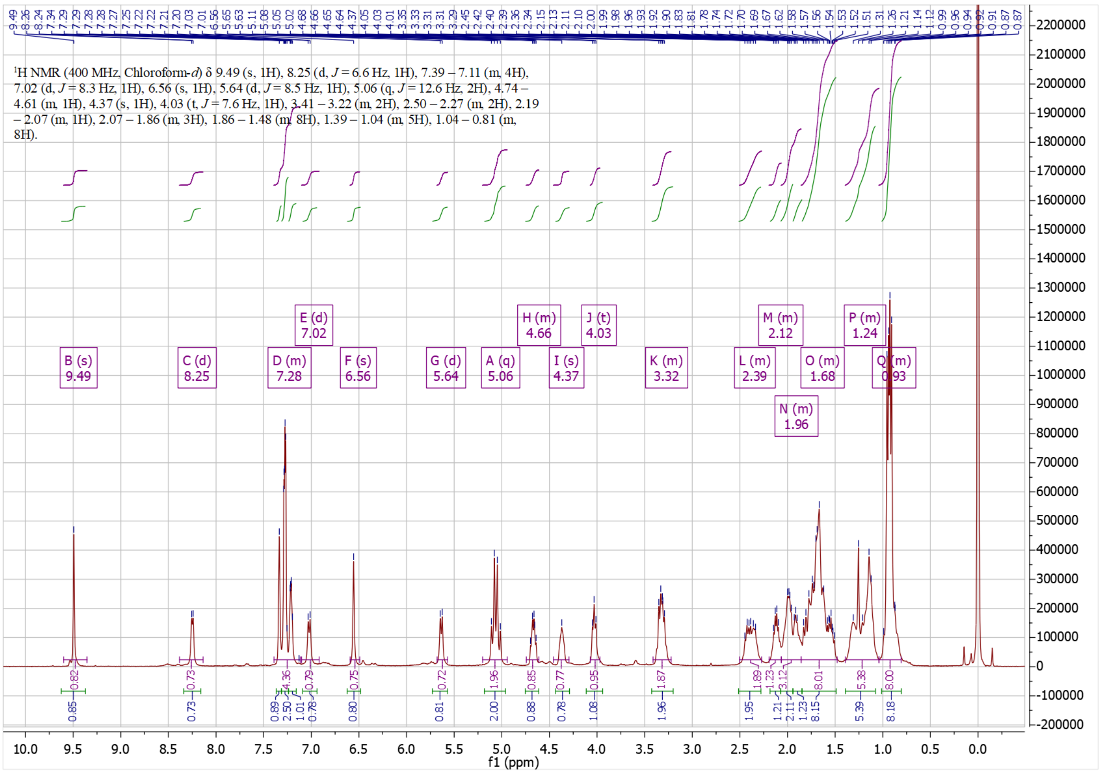

[1]H NMR (400 MHz, Chloroform-*d*) δ 9.49 (s, 1H), 8.25 (d, $J$ = 6.6 Hz, 1H), 7.39 – 7.11 (m, 4H), 7.02 (d, $J$ = 8.3 Hz, 1H), 6.56 (s, 1H), 5.64 (d, $J$ = 8.5 Hz, 1H), 5.06 (q, $J$ = 12.6 Hz, 2H), 4.74 – 4.61 (m, 1H), 4.37 (s, 1H), 4.03 (t, $J$ = 7.6 Hz, 1H), 3.41 – 3.22 (m, 2H), 2.50 – 2.27 (m, 2H), 2.19 – 2.07 (m, 1H), 2.07 – 1.86 (m, 3H), 1.86 – 1.48 (m, 8H), 1.39 – 1.04 (m, 5H), 1.04 – 0.81 (m, 8H).
f1 (ppm)


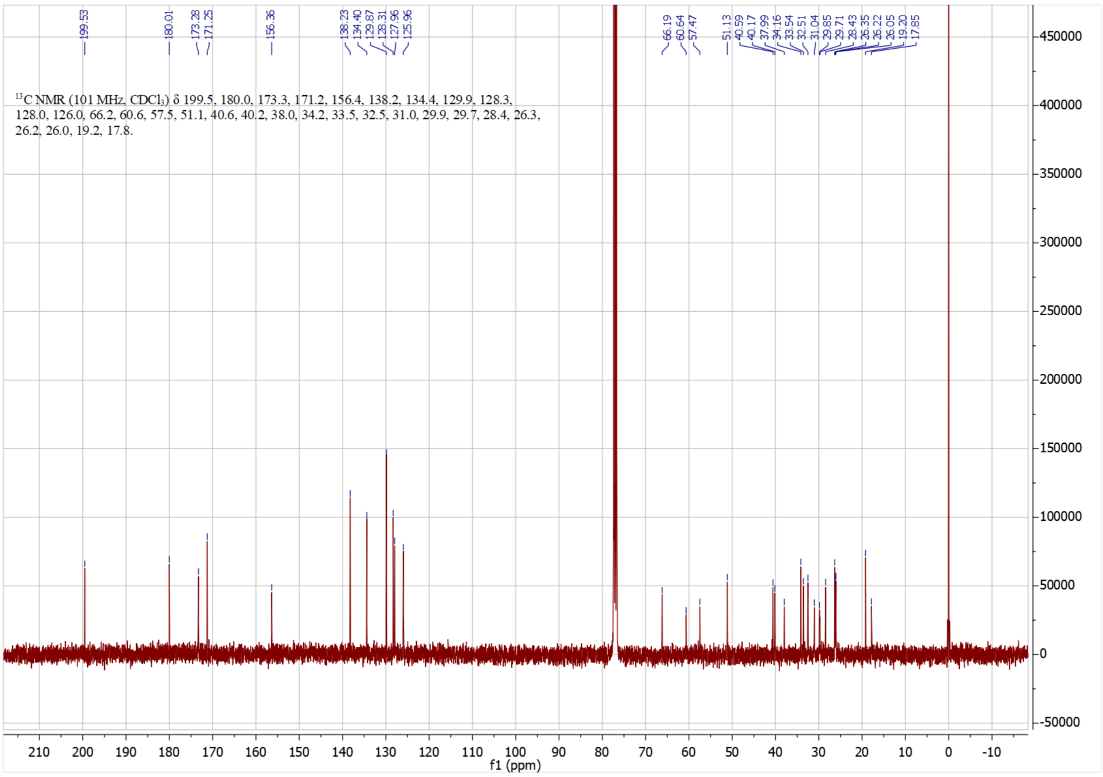

[13]C NMR (101 MHz, $CDCl_3$) δ 199.5, 180.0, 173.3, 171.2, 156.4, 138.2, 134.4, 129.9, 128.3, 128.0, 126.0, 66.2, 60.6, 57.5, 51.1, 40.6, 40.2, 38.0, 34.2, 33.5, 32.5, 31.0, 29.9, 29.7, 28.4, 26.3, 26.2, 26.0, 19.2, 17.8.
f1 (ppm)

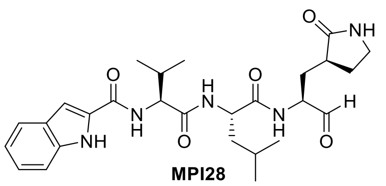

**MPI28**

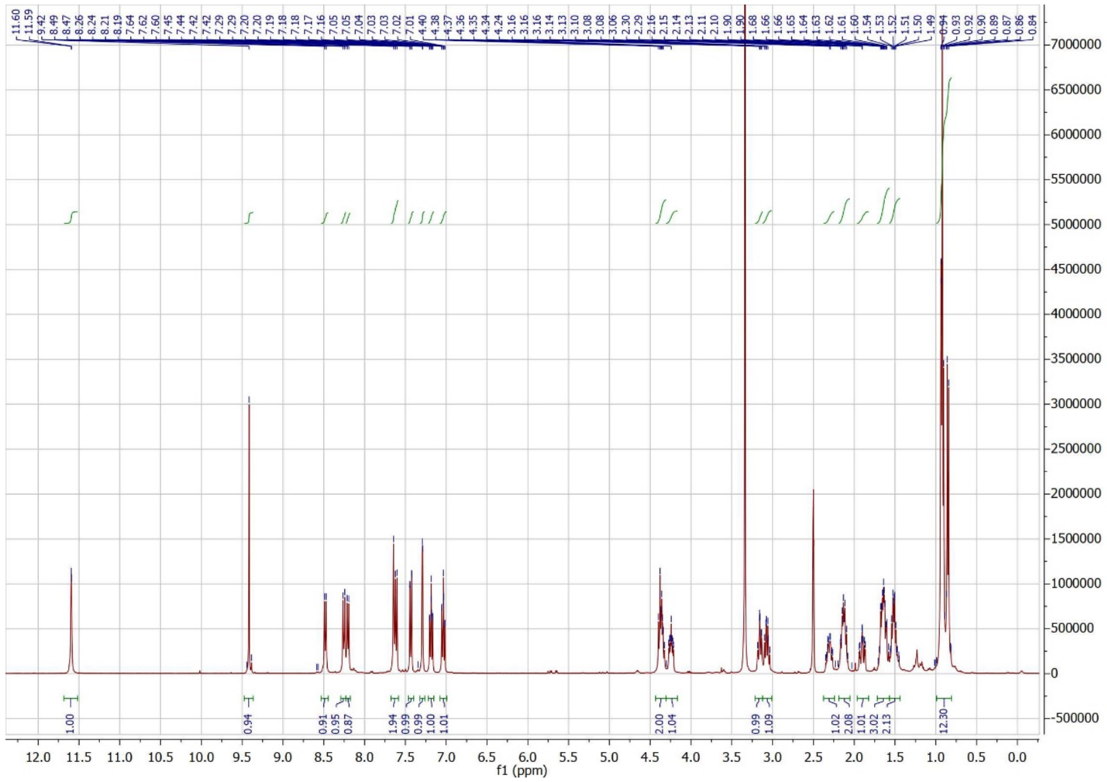
f1 (ppm)

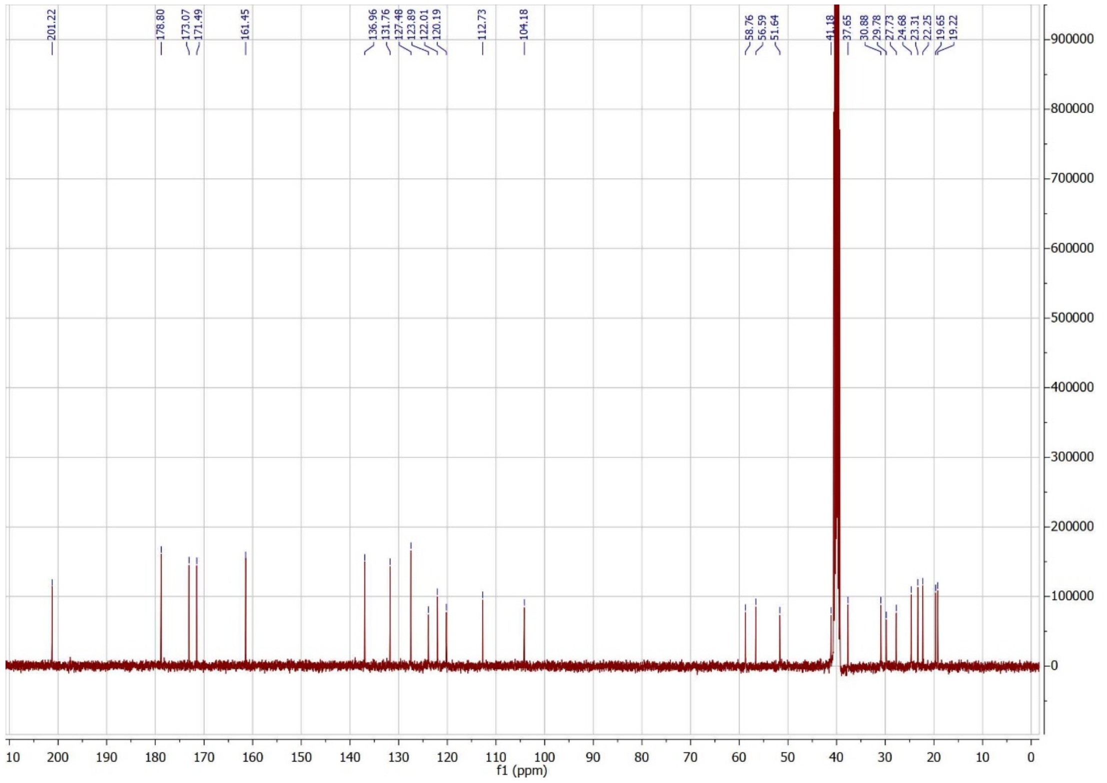
201.22
178.80
173.07
171.49
161.45
136.96
131.76
127.48
123.89
122.01
120.19
112.73
104.18
58.76
56.59
51.64
41.18
37.65
30.88
29.78
27.73
24.68
23.31
22.25
19.65
19.22
f1 (ppm)

#### HPLC Chromatographies of Potent MPI Compounds

### MPI 16

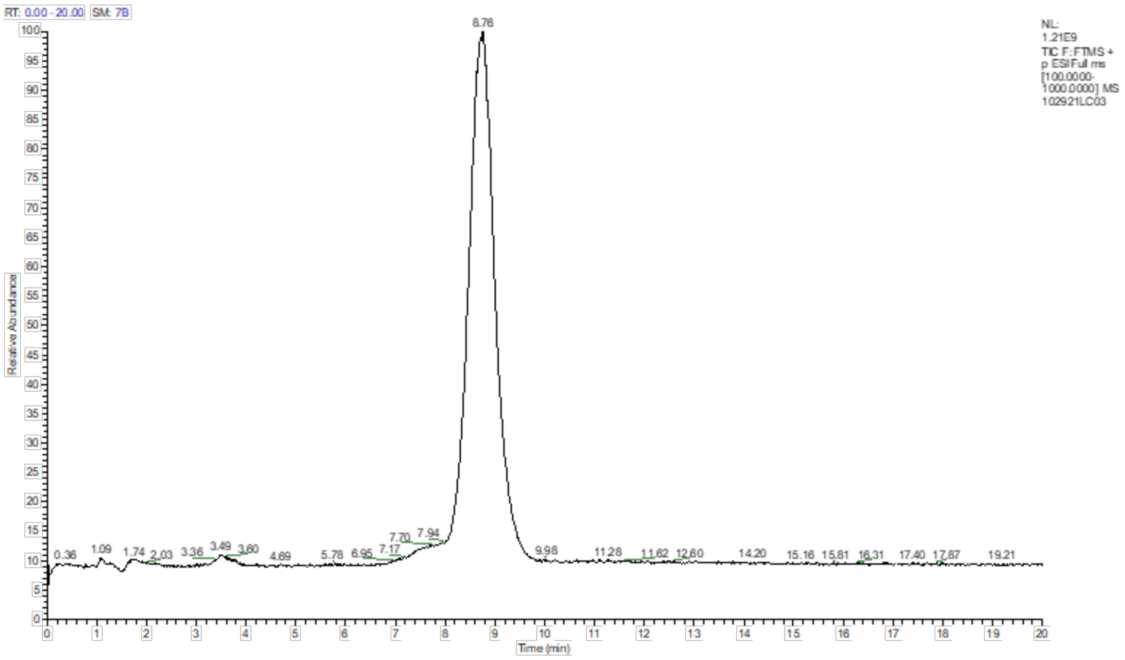


| | | | | | | |
|---|---|---|---|---|---|---|
| 1 | **Sample ID** | **MPI-16** | | | | |
| 2 | | | | | | |
| 3 | PEAK LIST | | | | | |
| 4 | 102921LC03.raw | | | | | |
| 5 | RT: 0.00 - 20.00 | | | | | |
| 6 | Number of detected peaks: 3 | | | | | |
| 7 | Apex RT | Start RT | End RT | Area | %Area | **Purity** |
| 8 | 3.5 | 2.61 | 4.08 | 111055586.4 | 0.6 | |
| 9 | 7.79 | 6.7 | 7.79 | 256284473.3 | 1.3 | |
| 10 | 8.76 | 7.83 | 10.11 | 18769702494 | 98.1 | **98%** |

| **No** | | | | | | |
|---|---|---|---|---|---|---|
| **1** | 0.000 | Equilibration | | | | |
| **2** | 0.000 | 0.300 | 0.0 | 70.0 | 0.0 | 5 |
| **3** | *New Row* | | | | | |
| **4** | 0.000 | Run | | | | |
| **5** | 5.000 | 0.300 | 0.0 | 70.0 | 0.0 | 5 |
| **6** | *New Row* | | | | | |
| **7** | 20.000 | Stop Run | | | | |

### MPI 17

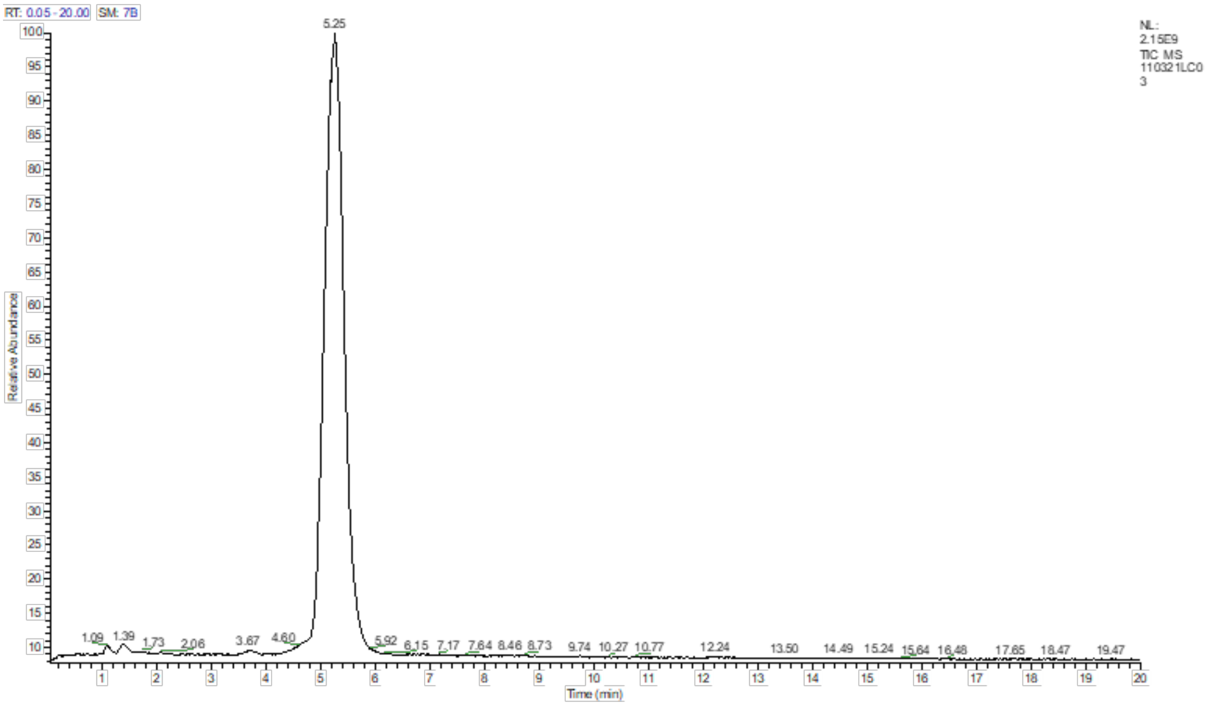


| | | | | | | |
|---|---|---|---|---|---|---|
| 1 | **Sample ID** | **MPI-17** | | | | |
| 2 | | | | | | |
| 3 | PEAK LIST | | | | | |
| 4 | 110321LC03.raw | | | | | |
| 5 | RT: 0.00 - 20.00 | | | | | |
| 6 | Number of detected peaks: 2 | | | | | |
| 7 | Apex RT | Start RT | End RT | Area | %Area | **PURITY** |
| 8 | 3.67 | 3.48 | 3.92 | 2.04E+08 | 0.41 | |
| 9 | 5.25 | 4.31 | 5.95 | 4.94E+10 | 99.59 | **99.6%** |

#### Experimental Condition

| No | Time | Flow [ml/min] | %B | %C | %D | Curve |
|---|---|---|---|---|---|---|
| 1 | 0.000 | Equilibration | | | | |
| 2 | 0.000 | 0.300 | 0.0 | 70.0 | 0.0 | 5 |
| 3 | *New Row* | | | | | |
| 4 | 0.000 | Run | | | | |
| 5 | 5.000 | 0.300 | 0.0 | 70.0 | 0.0 | 5 |
| 6 | *New Row* | | | | | |
| 7 | 20.000 | Stop Run | | | | |

### MPI 18

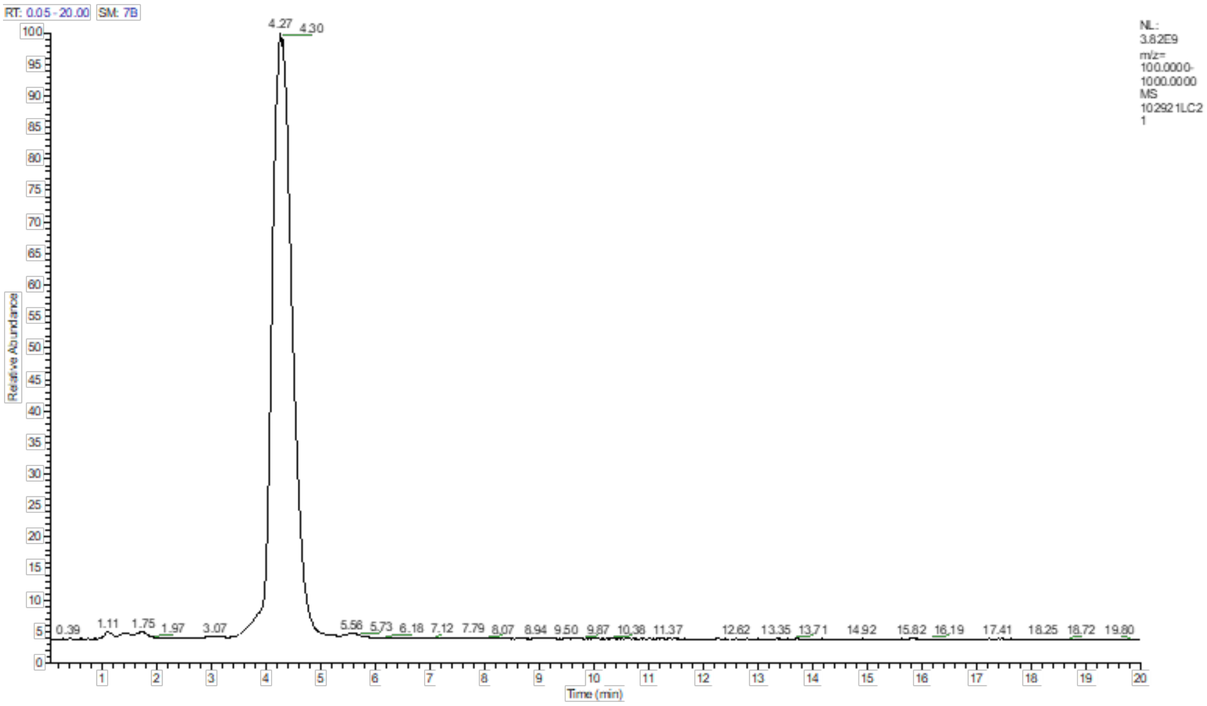


| 1 | Sample ID | MPI-18 | | | | |
|---|---|---|---|---|---|---|
| 2 | | | | | | |
| 3 | PEAK LIST | | | | | |
| 4 | 102921LC21.raw | | | | | |
| 5 | RT: 0.00 - 20.00 | | | | | |
| 6 | Number of detected peaks: 2 | | | | | |
| 7 | Apex RT | Start RT | End RT | Area | %Area | **Purity** |
| 8 | 4.27 | 3.52 | 4.98 | 9.61E+10 | 99.7 | **99.7%** |
| 9 | 5.56 | 5.33 | 5.78 | 2.87E+08 | 0.3 | |

#### Experimental Condition

| No | Time | Flow [ml/min] | %B | %C | %D | Curve |
|---|---|---|---|---|---|---|
| 1 | 0.000 | Equilibration | | | | |
| 2 | 0.000 | 0.300 | 0.0 | 70.0 | 0.0 | 5 |
| 3 | *New Row* | | | | | |
| 4 | 0.000 | Run | | | | |
| 5 | 5.000 | 0.300 | 0.0 | 70.0 | 0.0 | 5 |
| 6 | *New Row* | | | | | |
| 7 | 20.000 | Stop Run | | | | |

### MPI 25

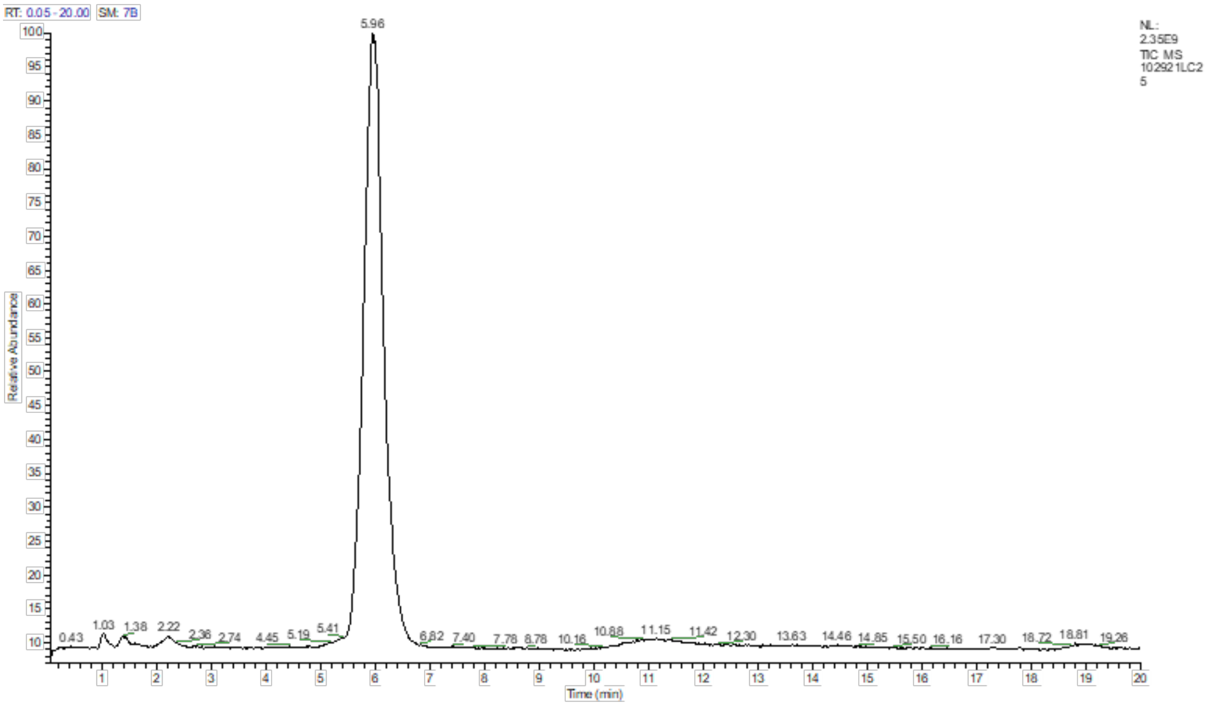


| 1 | Sample ID | MPI-25 | | | |
|---|---|---|---|---|---|
| 2 | | | | | |
| 3 | PEAK LIST | | | | |
| 4 | 102921LC25.raw | | | | |
| 5 | RT: 0.00 - 20.00 | | | | |
| 6 | Number of detected peaks: 3 | | | | |
| 7 | Apex RT | Start RT | End RT | Area | %Area | Purity |
| 8 | 2.22 | 1.92 | 2.6 | 6.28E+08 | 1.08 | |
| 9 | 5.96 | 5 | 6.72 | 5.54E+10 | 95.68 | 96% |
| 10 | 11.15 | 10.13 | 12.18 | 1.87E+09 | 3.23 | |

#### Experimental Condition

| No | Time | Flow [ml/min] | %B | %C | %D | Curve |
|---|---|---|---|---|---|---|
| 1 | 0.000 | Equilibration | | | | |
| 2 | 0.000 | 0.300 | 0.0 | 80.0 | 0.0 | 5 |
| 3 | *New Row* | | | | | |
| 4 | 0.000 | Run | | | | |
| 5 | 5.000 | 0.300 | 0.0 | 80.0 | 0.0 | 5 |
| 6 | *New Row* | | | | | |
| 7 | 20.000 | Stop Run | | | | |

### MPI 26

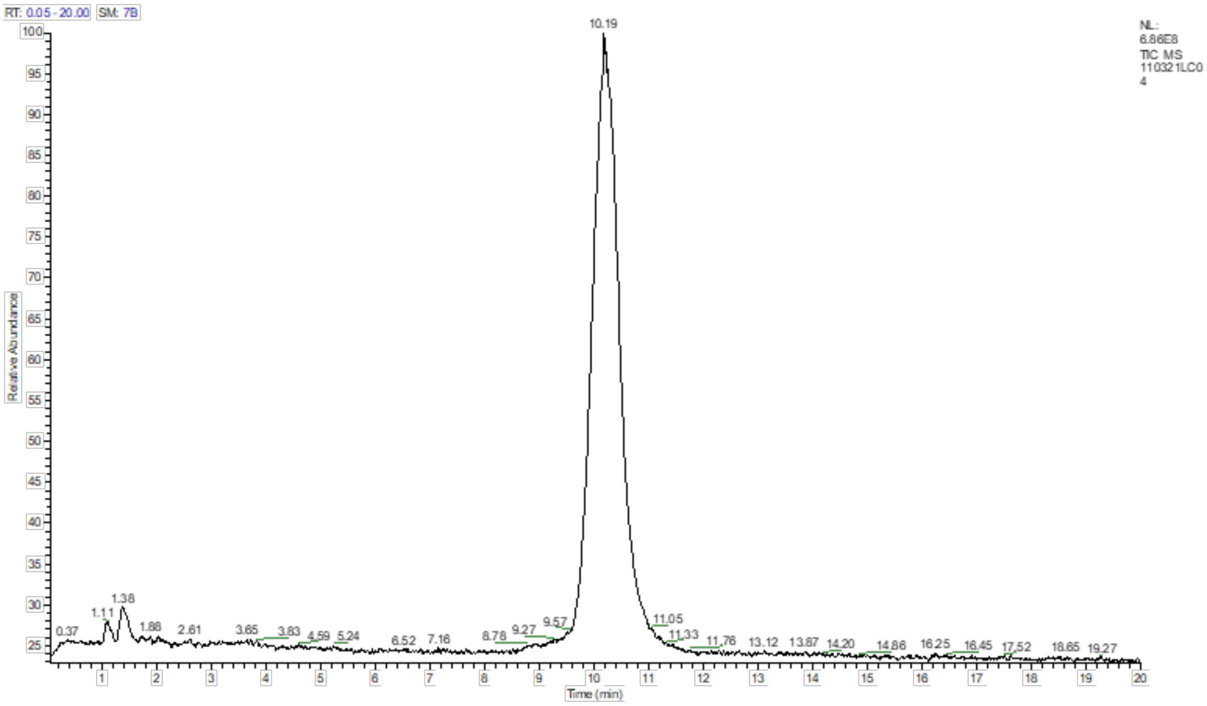

| | | | | | | |
|---|---|---|---|---|---|---|
| 1 | **Sample ID** | **MPI-26** | | | | |
| 2 | | | | | | |
| 3 | PEAK LIST | | | | | |
| 4 | 110321LC04.raw | | | | | |
| 5 | RT: 0.00 - 20.00 | | | | | |
| 6 | Number of detected peaks: 2 | | | | | |
| 7 | Apex RT | Start RT | End RT | Area | %Area | **PURITY** |
| 8 | 8.91 | 8.67 | 9.1 | 97515510 | 0.55 | |
| 9 | 10.19 | 9.43 | 11.59 | 1.77E+10 | 99.45 | **99.5%** |

#### Experimental Condition

| No | Time | Flow [ml/min] | %B | %C | %D | Curve |
|---|---|---|---|---|---|---|
| 1 | 0.000 | Equilibration | | | | |
| 2 | 0.000 | 0.300 | 0.0 | 70.0 | 0.0 | 5 |
| 3 | *New Row* | | | | | |
| 4 | 0.000 | Run | | | | |
| 5 | 5.000 | 0.300 | 0.0 | 70.0 | 0.0 | 5 |
| 6 | *New Row* | | | | | |
| 7 | 20.000 | Stop Run | | | | |

### MPI 27

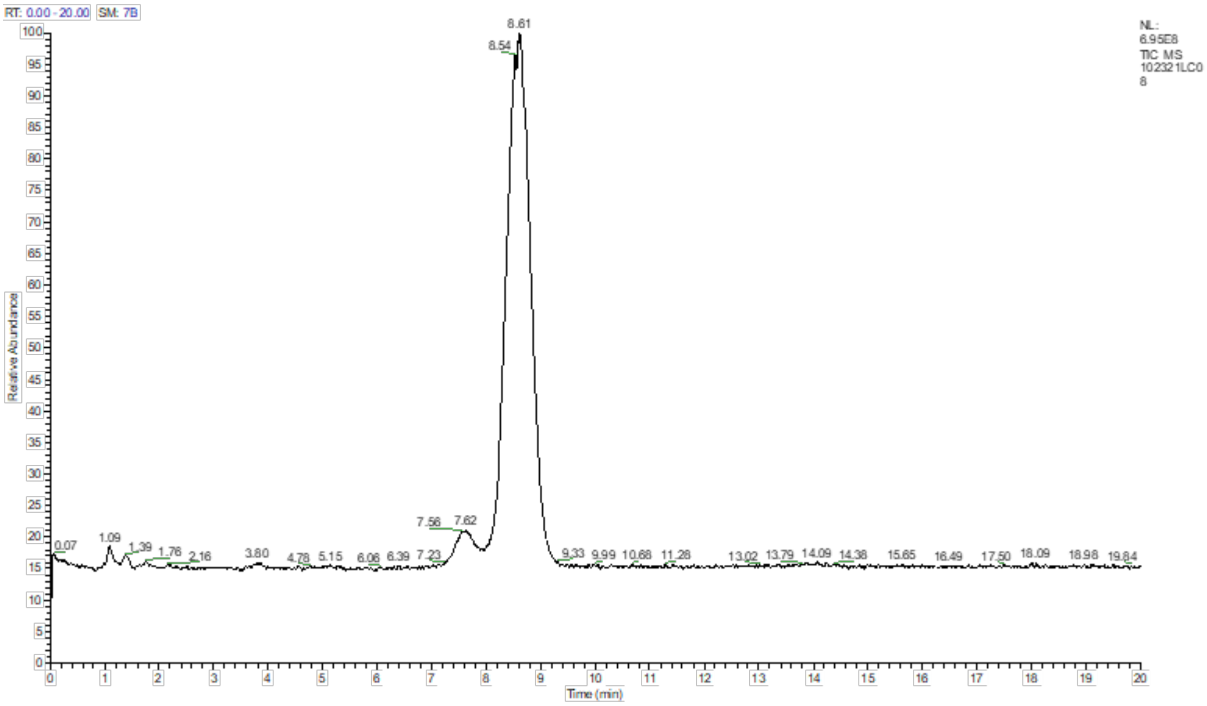


| | | | | | | |
|---|---|---|---|---|---|---|
| 1 | **Sample ID** | **MPI27** | | | | |
| 2 | | | | | | |
| 3 | PEAK LIST | | | | | |
| 4 | 102321LC08.raw | | | | | |
| 5 | RT: 0.00 - 20.00 | | | | | |
| 6 | Number of detected peaks: 2 | | | | | |
| 7 | Apex RT | Start RT | End RT | Area | %Area | **PURITY** |
| 8 | 7.62 | 7.21 | 7.89 | 9.44E+08 | 4.87 | |
| 9 | 8.61 | 7.89 | 9.31 | 1.85E+10 | 95.13 | **95%** |

#### Experimental Condition

| No | Time | Flow [ml/min] | %B | %C | %D | Curve |
|---|---|---|---|---|---|---|
| 1 | 0.000 | Equilibration | | | | |
| 2 | 0.000 | 0.300 | 0.0 | 70.0 | 0.0 | 5 |
| 3 | *New Row* | | | | | |
| 4 | 0.000 | Run | | | | |
| 5 | 5.000 | 0.300 | 0.0 | 70.0 | 0.0 | 5 |
| 6 | *New Row* | | | | | |
| 7 | 20.000 | Stop Run | | | | |

**Flow Cytometry Images for MPIs**

(compound concentration labeled)

### MPI11

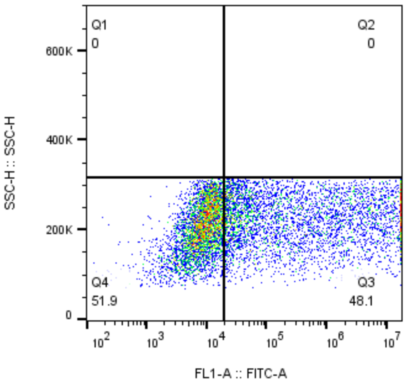


MPI11-10uM
HEK
8503

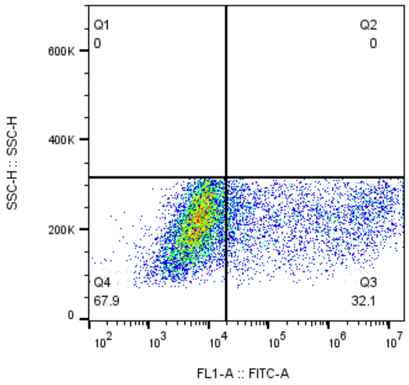


MPI11-2uM
HEK
8864

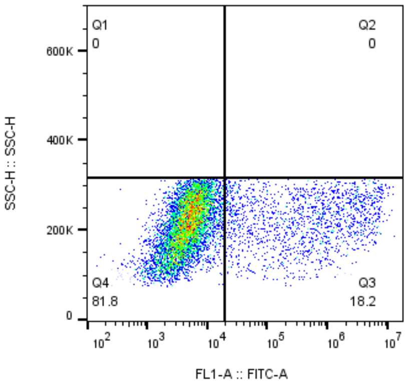


MPI11-400nM
HEK
8814

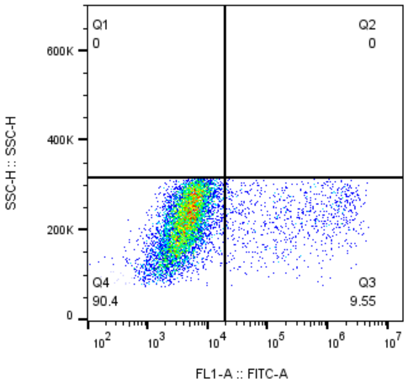


MPI11-80nM
HEK
8437

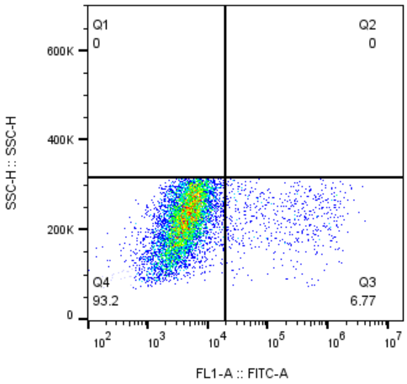


MPI11-16nM
HEK
8238

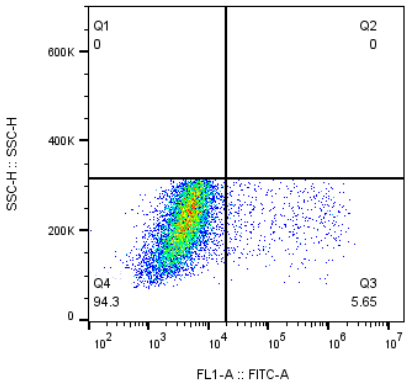


MPI11-3.2nM
HEK
8566

### MPI12

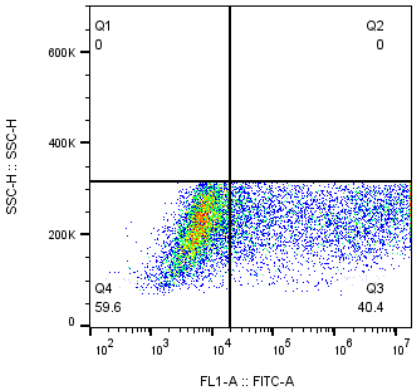


MPI12-10uM
HEK
9939

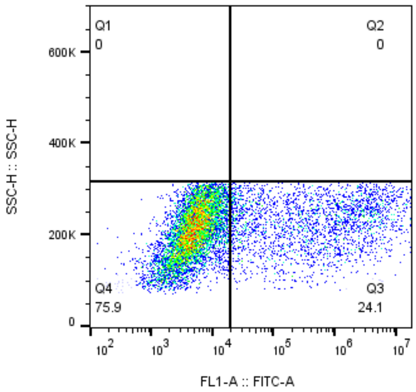


MPI12-2uM
HEK
10342

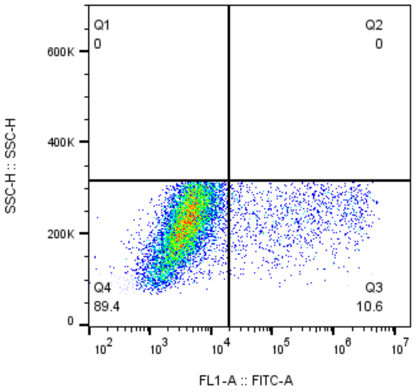


MPI12-400nM
HEK
9916

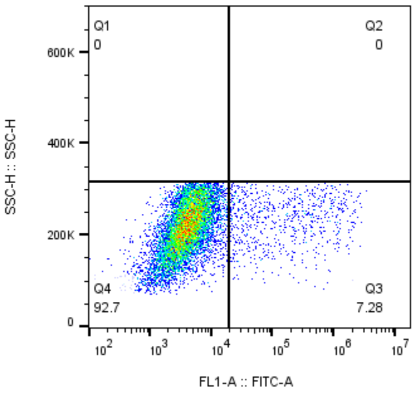


MPI12-80nM
HEK
10198

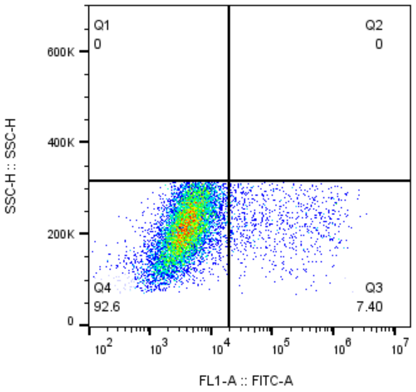


MPI12-16nM
HEK
10331

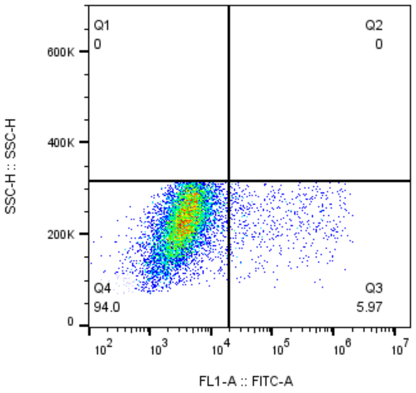


MPI12-3.2nM
HEK
8893

### MPI13

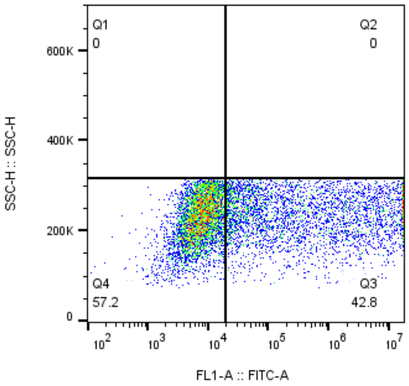


MPI13-10uM
HEK
8217

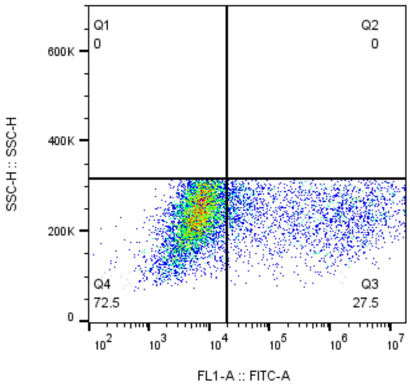


MPI13-2uM
HEK
8105

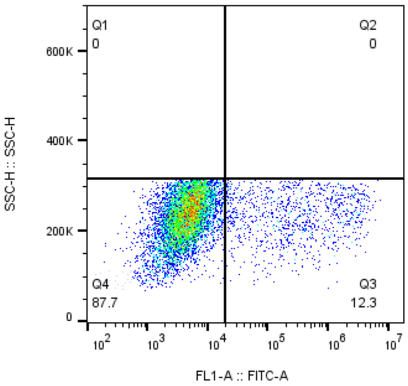


MPI13-400nM
HEK
8537

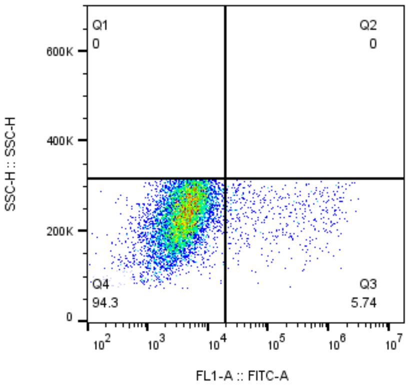


MPI13-80nM
HEK
8138

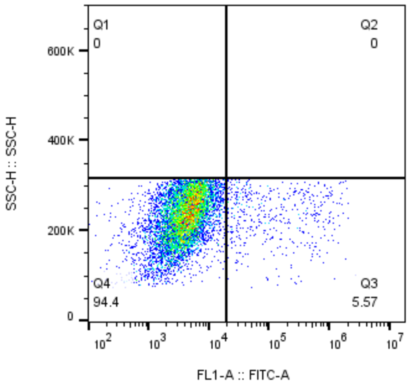


MPI13-16nM
HEK
8397

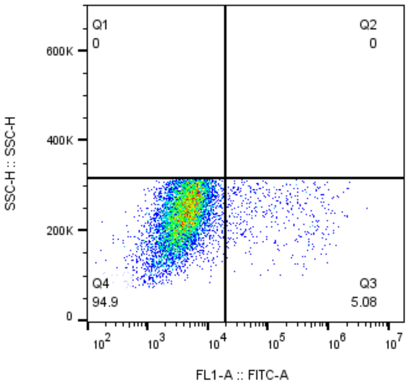


MPI13-3.2nM
HEK
8191

### MPI14

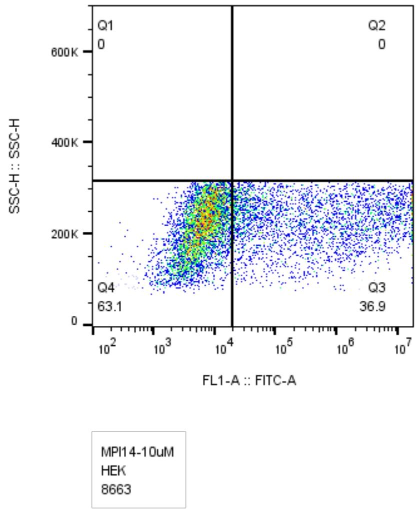
Q1
0
Q2
0
Q4
63.1
Q3
36.9
SSC-H :: SSC-H
FL1-A :: FITC-A
MPI14-10uM
HEK
8663

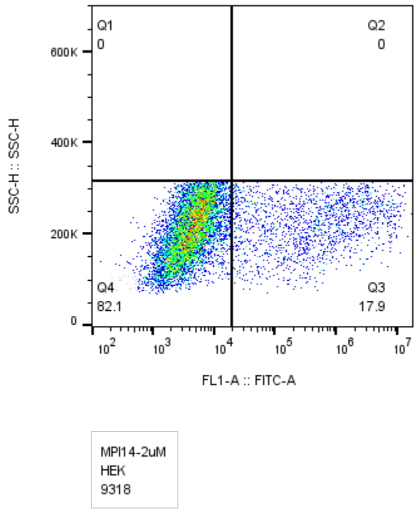
Q1
0
Q2
0
Q4
82.1
Q3
17.9
SSC-H :: SSC-H
FL1-A :: FITC-A
MPI14-2uM
HEK
9318

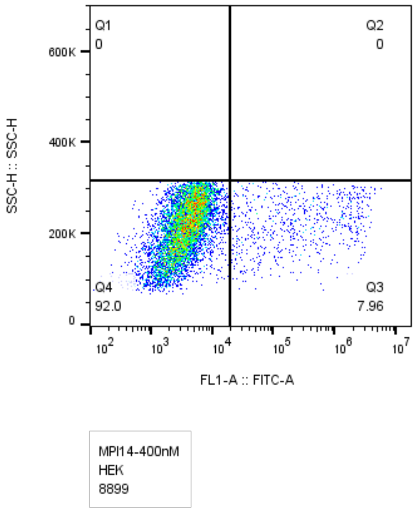
Q1
0
Q2
0
Q4
92.0
Q3
7.96
SSC-H :: SSC-H
FL1-A :: FITC-A
MPI14-400nM
HEK
8899

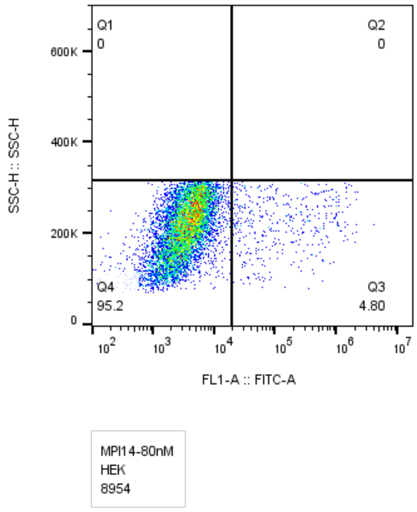
Q1
0
Q2
0
Q4
95.2
Q3
4.80
SSC-H :: SSC-H
FL1-A :: FITC-A
MPI14-80nM
HEK
8954

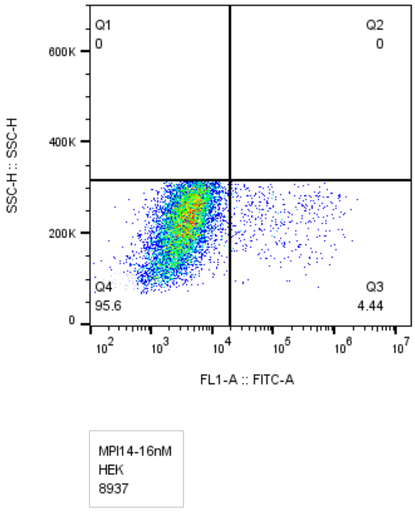
Q1
0
Q2
0
Q4
95.6
Q3
4.44
SSC-H :: SSC-H
FL1-A :: FITC-A
MPI14-16nM
HEK
8937

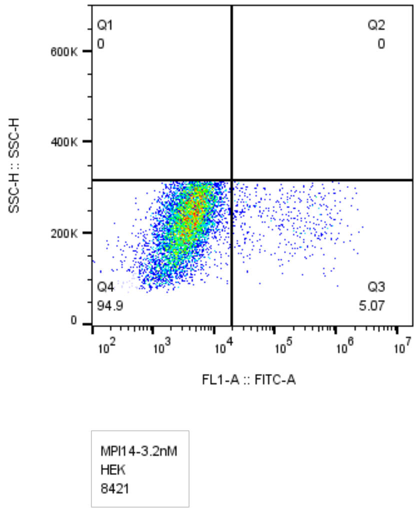
Q1
0
Q2
0
Q4
94.9
Q3
5.07
SSC-H :: SSC-H
FL1-A :: FITC-A
MPI14-3.2nM
HEK
8421

### MPI16

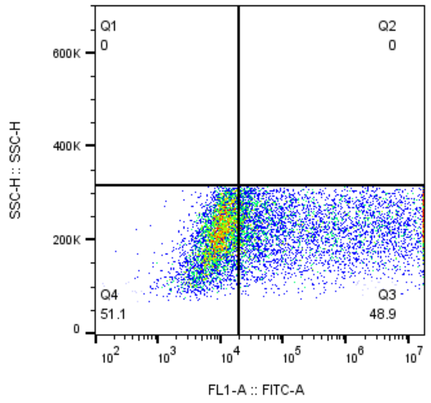


MPI16-10uM
HEK
9121

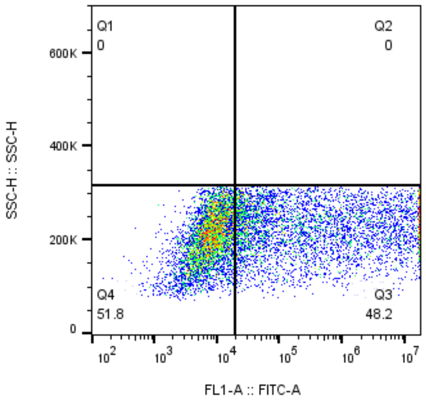


MPI16-2uM
HEK
9316

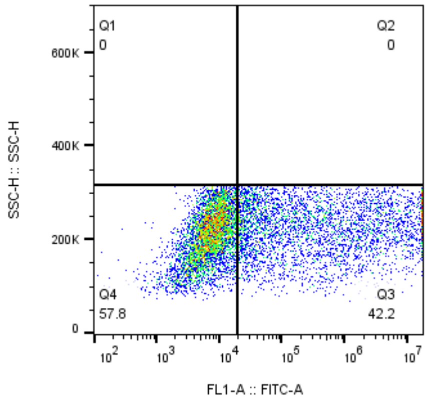


MPI16-400nM
HEK
9078

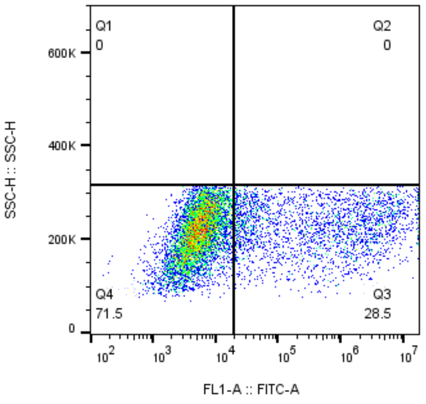


MPI16-80nM
HEK
9190

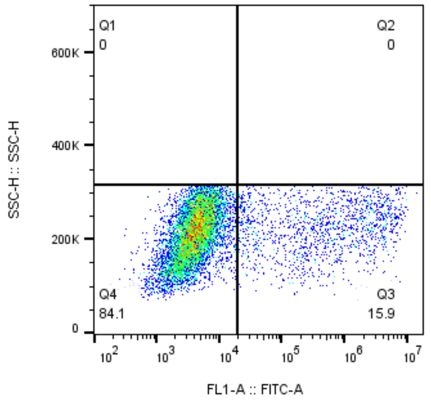


MPI16-16nM
HEK
9193

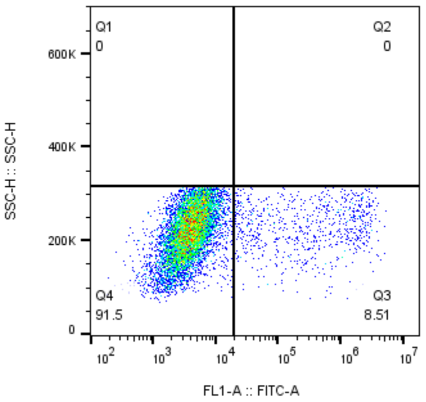


MPI16-3.2nM
HEK
8670

### MPI17

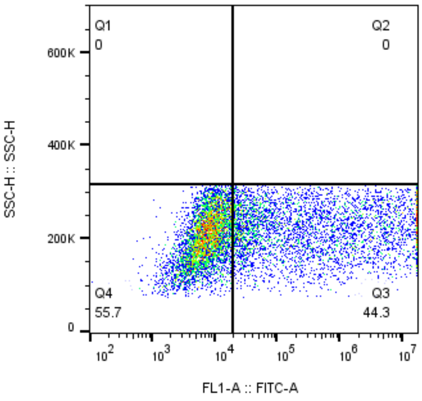


MPI17-10uM
HEK
9139

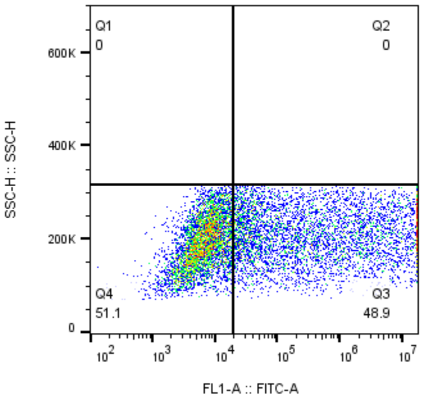


MPI17-2uM
HEK
9915

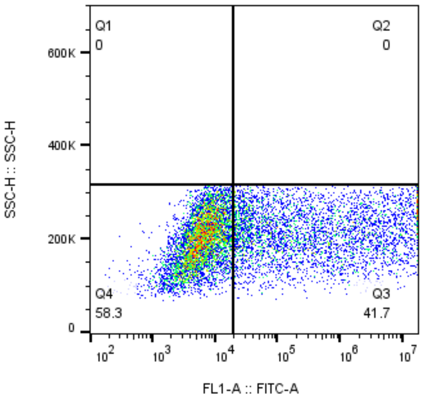


MPI17-400nM
HEK
9653

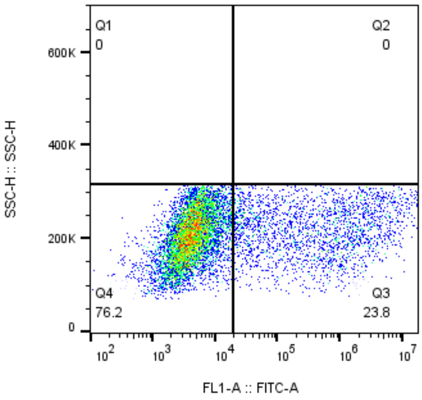


MPI17-80nM
HEK
9446

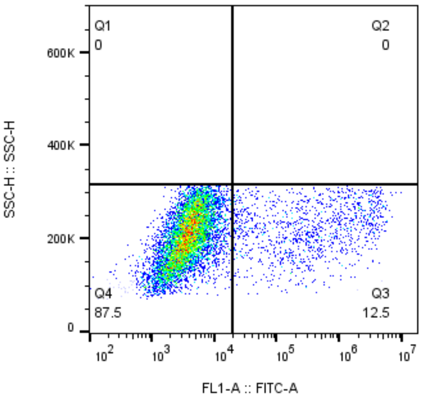


MPI17-16nM
HEK
9369

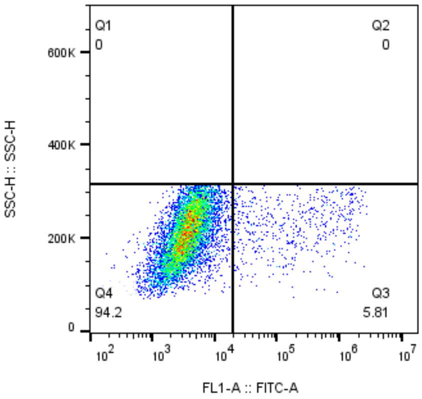


MPI17-3.2nM
HEK
9257

### MPI18

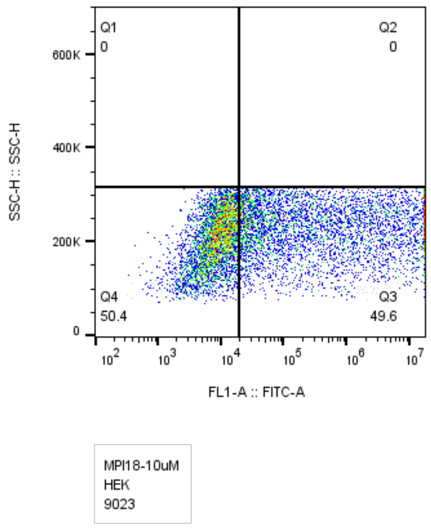
Q1
0
Q2
0
Q4
50.4
Q3
49.6
SSC-H :: SSC-H
FL1-A :: FITC-A
MPI18-10uM
HEK
9023

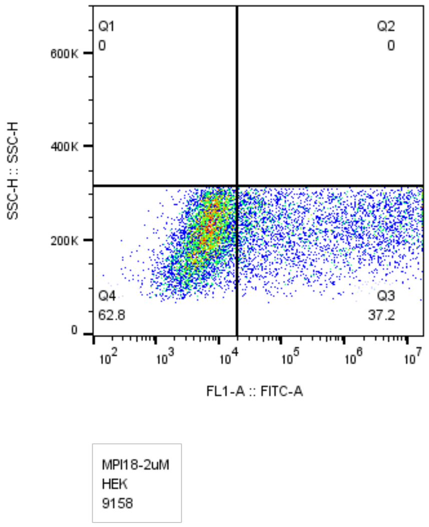
Q1
0
Q2
0
Q4
62.8
Q3
37.2
SSC-H :: SSC-H
FL1-A :: FITC-A
MPI18-2uM
HEK
9158

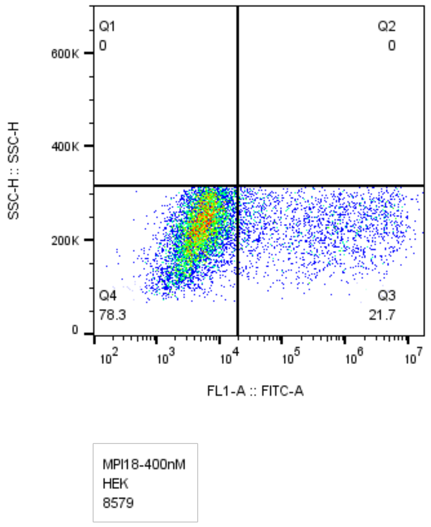
Q1
0
Q2
0
Q4
78.3
Q3
21.7
SSC-H :: SSC-H
FL1-A :: FITC-A
MPI18-400nM
HEK
8579

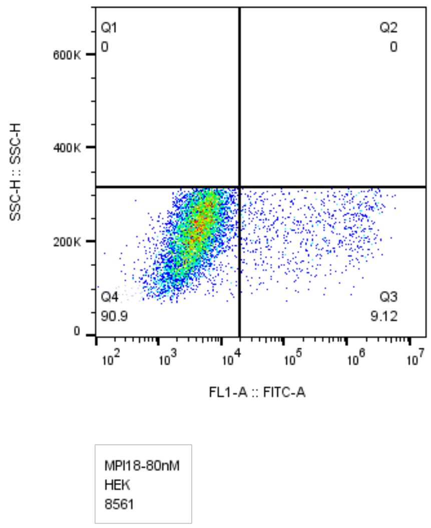
Q1
0
Q2
0
Q4
90.9
Q3
9.12
SSC-H :: SSC-H
FL1-A :: FITC-A
MPI18-80nM
HEK
8561

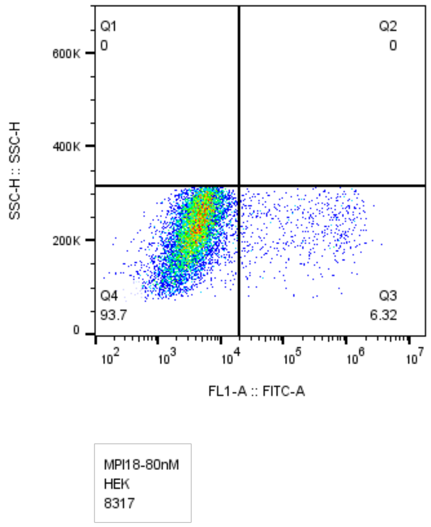
Q1
0
Q2
0
Q4
93.7
Q3
6.32
SSC-H :: SSC-H
FL1-A :: FITC-A
MPI18-80nM
HEK
8317

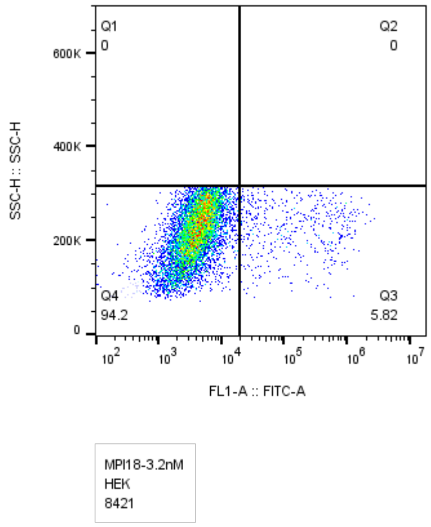
Q1
0
Q2
0
Q4
94.2
Q3
5.82
SSC-H :: SSC-H
FL1-A :: FITC-A
MPI18-3.2nM
HEK
8421

### MPI19

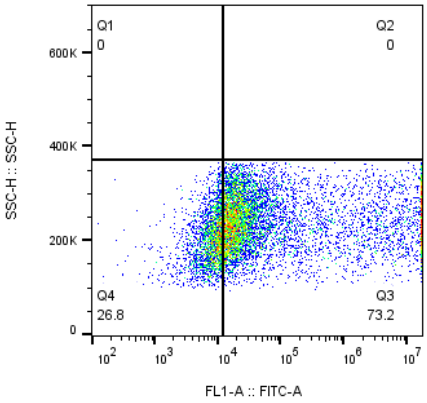


MPI19-10uM
HEK
9577

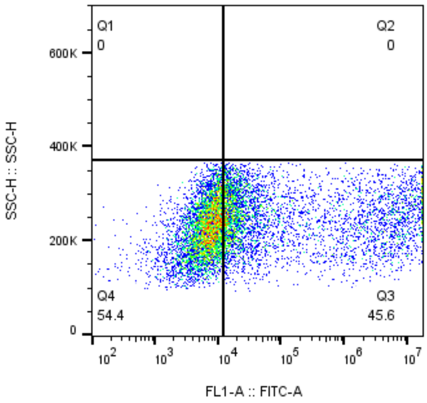


MPI19-2uM
HEK
9958

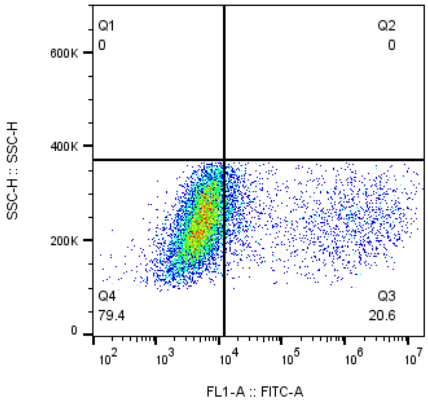


MPI19-400nM
HEK
9672

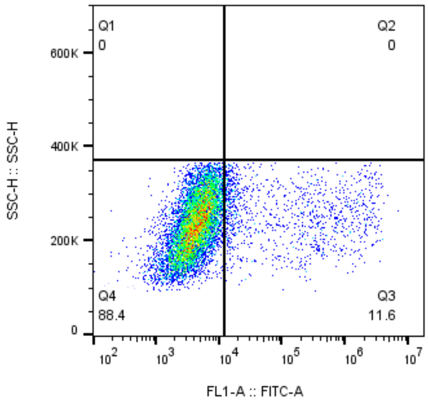


MPI19-80nM
HEK
9771

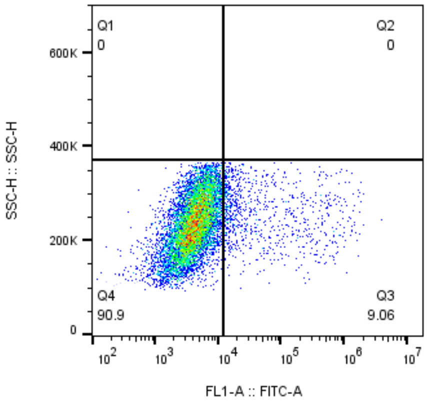


MPI19-16nM
HEK
9756

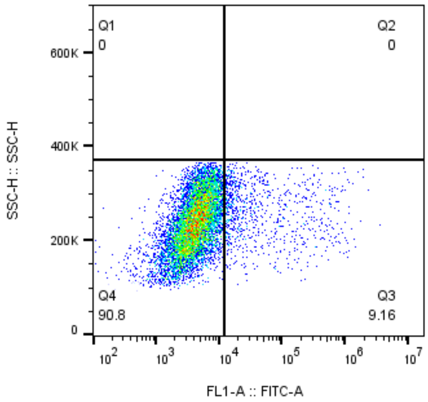


MPI19-3.2nM
HEK
9501

### MPI20

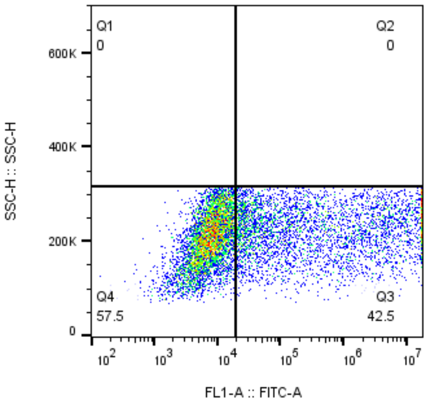


MPI20-10uM
HEK
9252

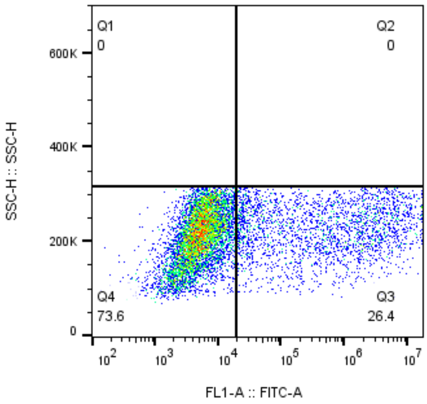


MPI20-2uM
HEK
9242

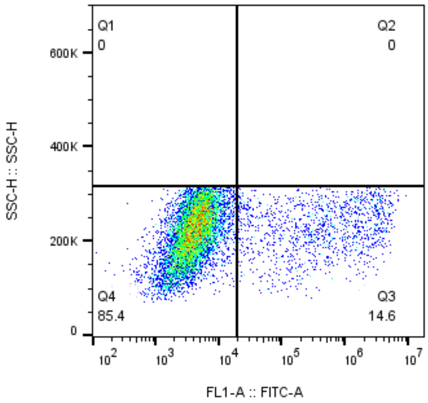


MPI20-400nM
HEK
8719

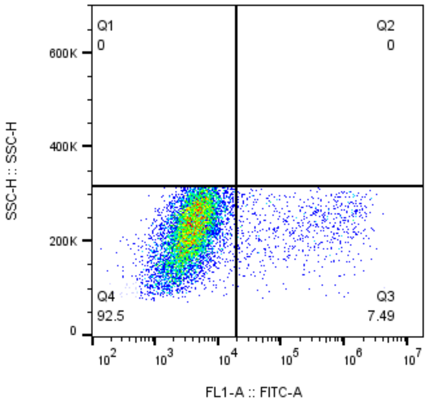


MPI20-80nM
HEK
8763

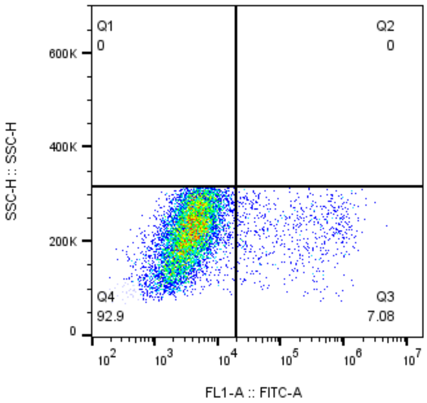


MPI20-16nM
HEK
8988

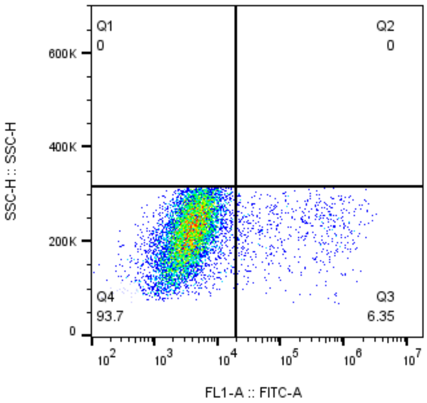


MPI20-3.2nM
HEK
8886

### MPI21

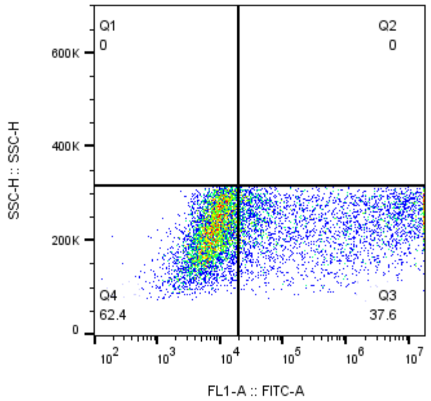


MPI21-10uM
HEK
8396

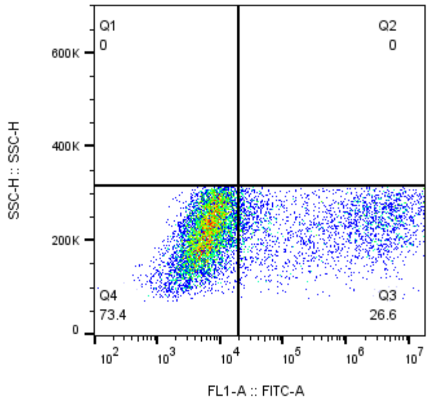


MPI21-2uM
HEK
8758

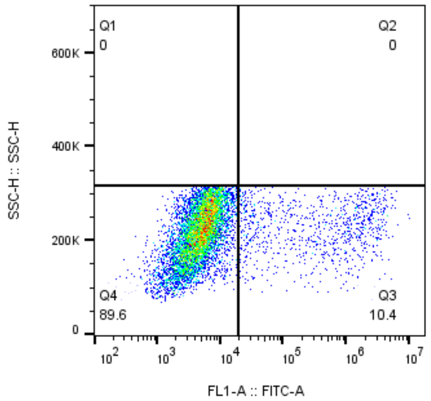


MPI21-400nM
HEK
8693

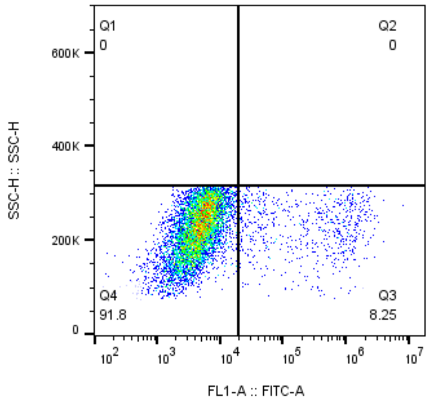


MPI21-80nM
HEK
8101

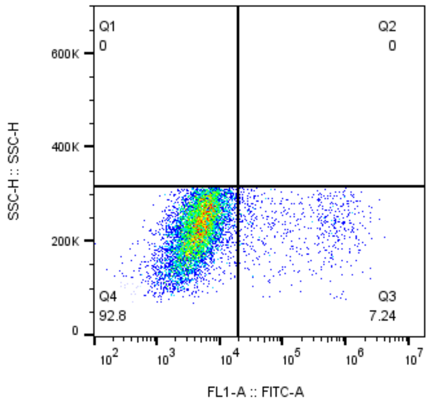


MPI21-16nM
HEK
8162

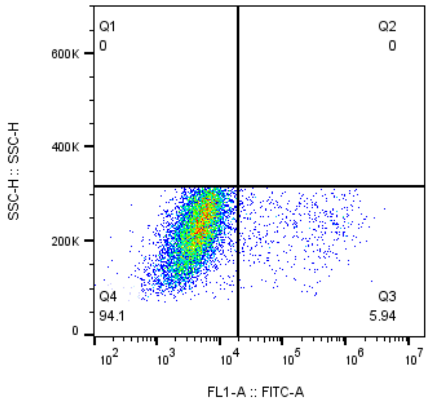


MPI21-3.2nM
HEK
8402

### MPI22

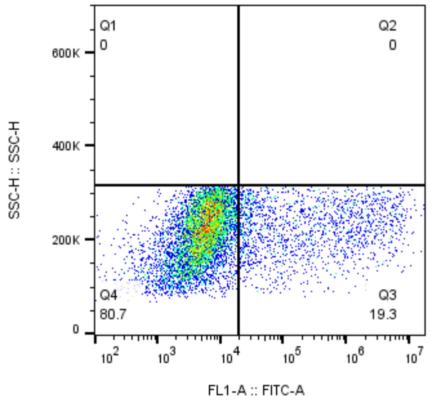


MPI22-10uM
HEK
9010

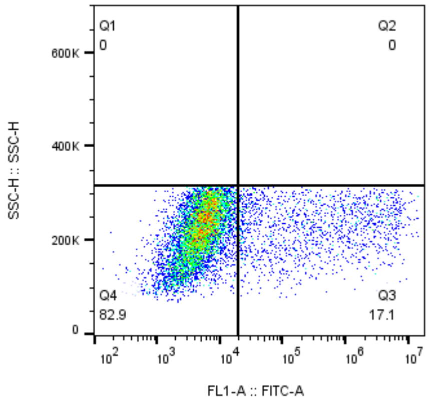


MPI22-2uM
HEK
8654

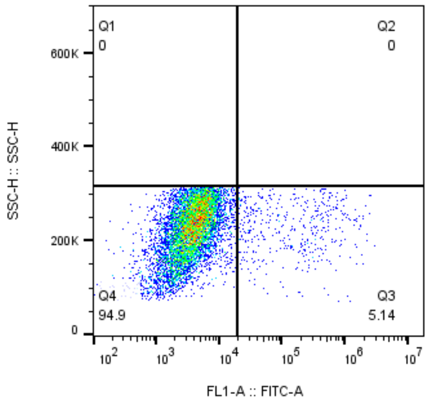


MPI22-400nM
HEK
8364

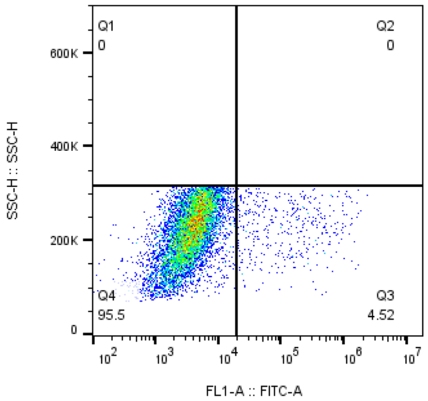


MPI22-80nM
HEK
8718

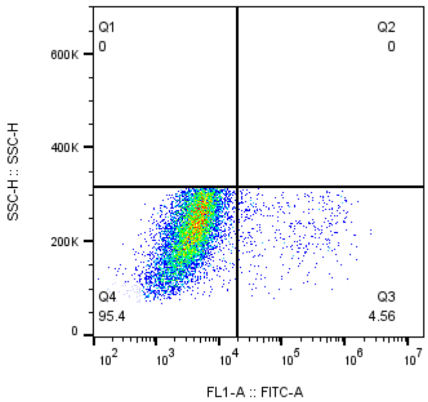


MPI22-16nM
HEK
8647

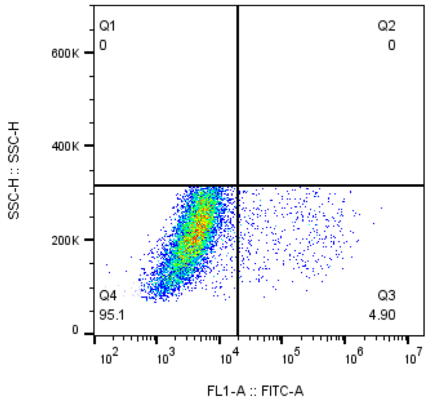


MPI22-3.2nM
HEK
8774

### MPI23

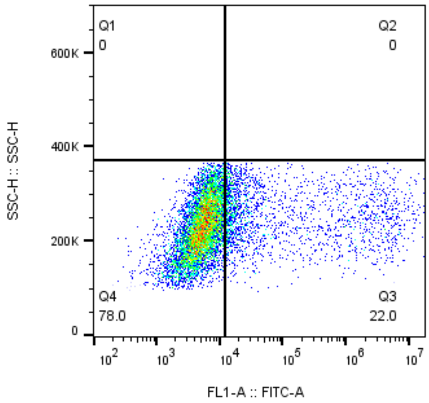


MPI23-10uM
HEK
9295

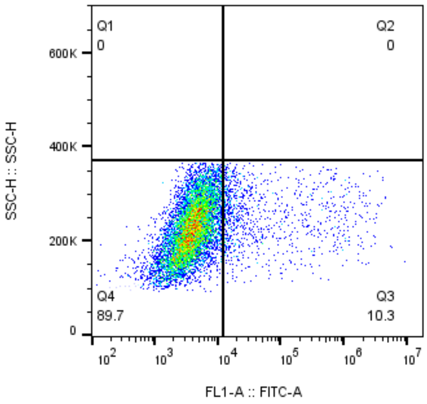


MPI23-2uM
HEK
9530

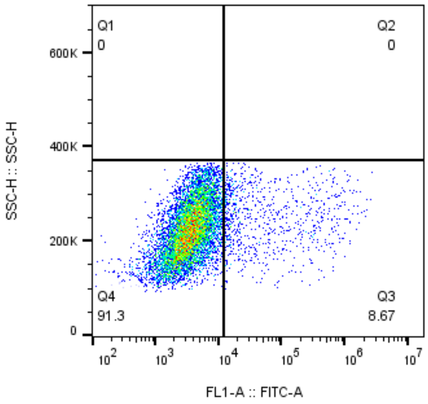


MPI23-400nM
HEK
9495

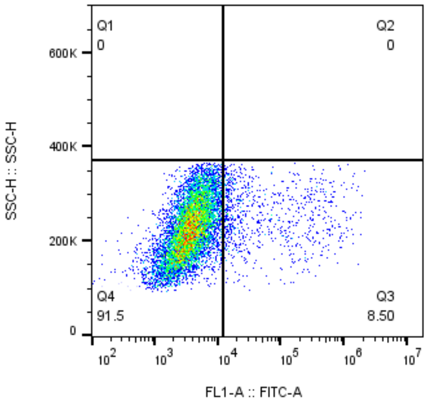


MPI23-80nM
HEK
10038

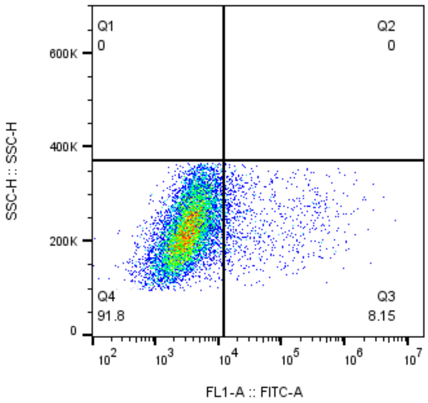


MPI23-16nM
HEK
9504

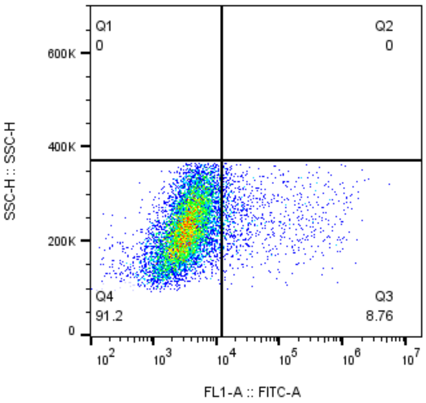


MPI23-3.2nM
HEK
9690

### MPI24

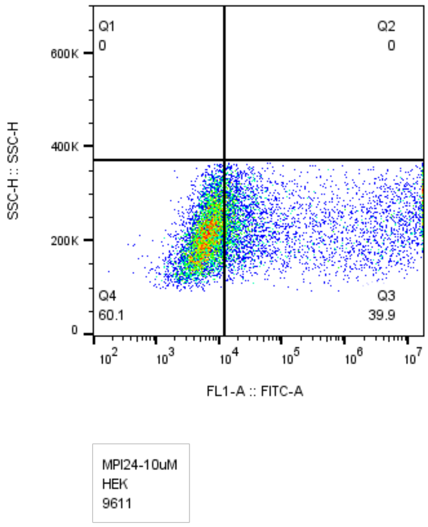
Q1
0
Q2
0
Q4
60.1
Q3
39.9
SSC-H :: SSC-H
FL1-A :: FITC-A
MPI24-10uM
HEK
9611

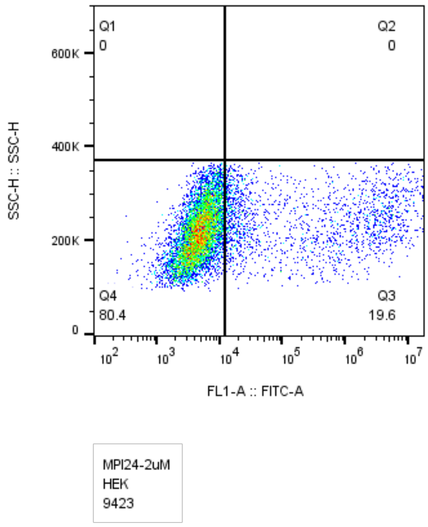
Q1
0
Q2
0
Q4
80.4
Q3
19.6
SSC-H :: SSC-H
FL1-A :: FITC-A
MPI24-2uM
HEK
9423

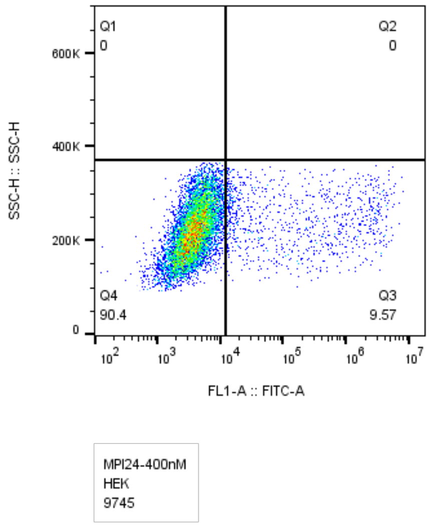
Q1
0
Q2
0
Q4
90.4
Q3
9.57
SSC-H :: SSC-H
FL1-A :: FITC-A
MPI24-400nM
HEK
9745

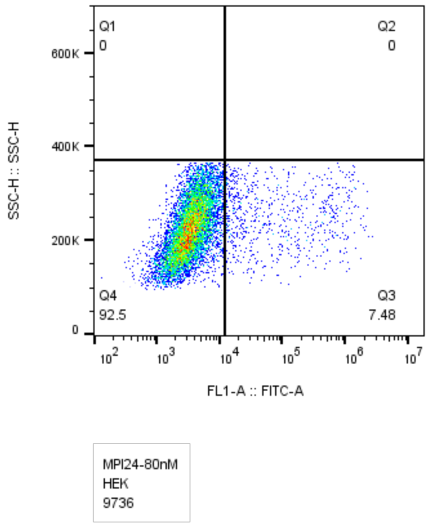
Q1
0
Q2
0
Q4
92.5
Q3
7.48
SSC-H :: SSC-H
FL1-A :: FITC-A
MPI24-80nM
HEK
9736

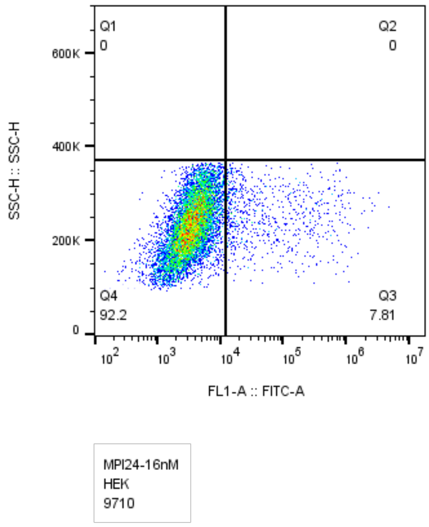
Q1
0
Q2
0
Q4
92.2
Q3
7.81
SSC-H :: SSC-H
FL1-A :: FITC-A
MPI24-16nM
HEK
9710

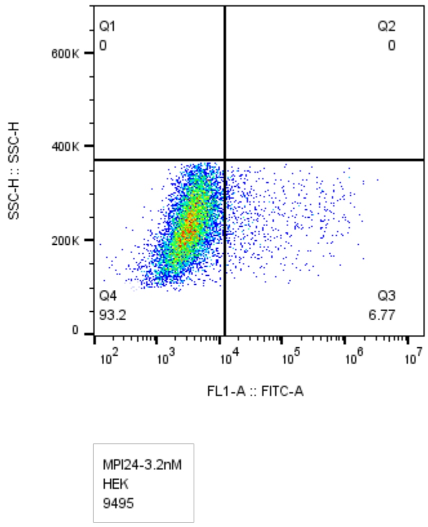
Q1
0
Q2
0
Q4
93.2
Q3
6.77
SSC-H :: SSC-H
FL1-A :: FITC-A
MPI24-3.2nM
HEK
9495

### MPI25

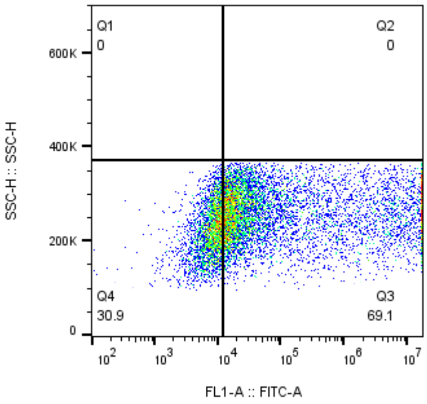


MPI25-10uM
HEK
9749

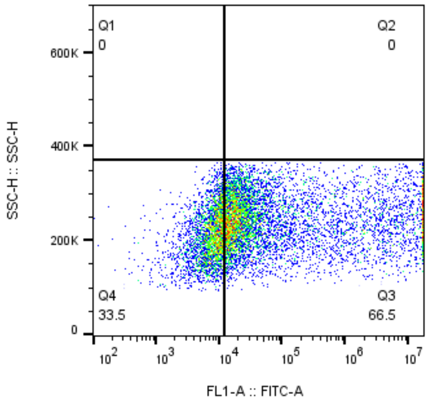


MPI25-2uM
HEK
9793

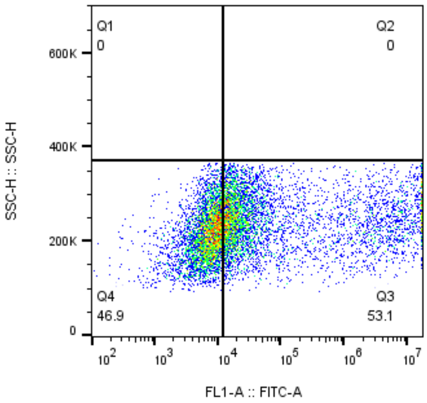


MPI25-400nM
HEK
9597

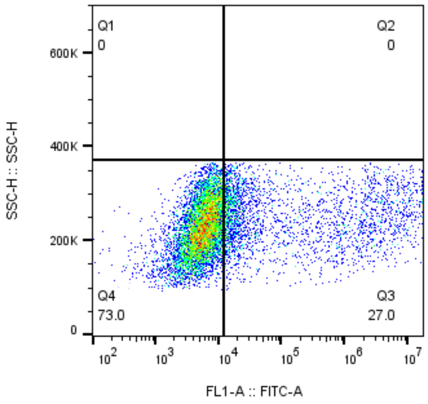


MPI25-80nM
HEK
9906

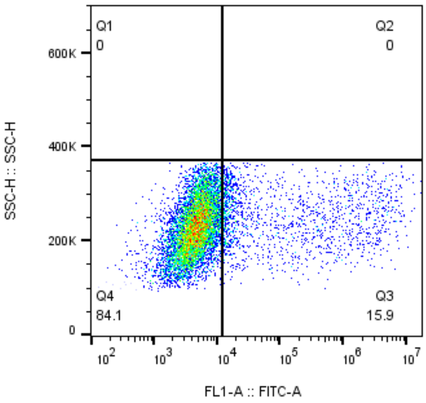


MPI25-16nM
HEK
10032

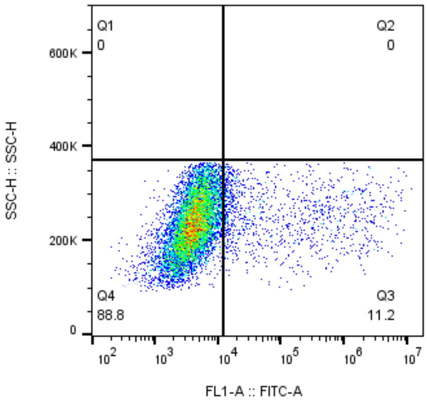


MPI25-3.2nM
HEK
9797

### MPI26

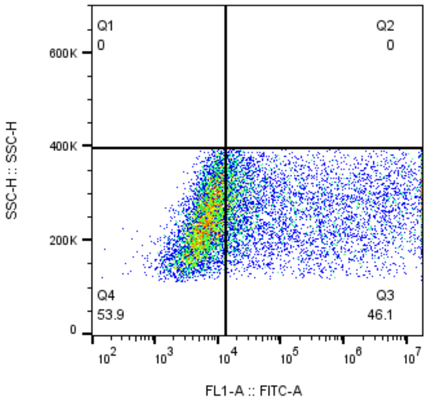


MPI26-10uM
HEK
11089

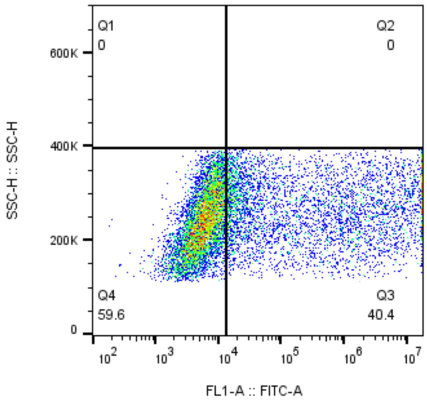


MPI26-2uM
HEK
11229

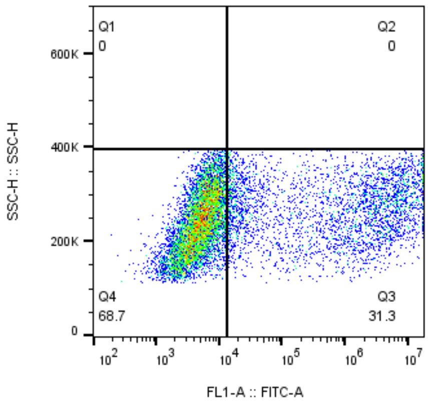


MPI26-400nM
HEK
10873

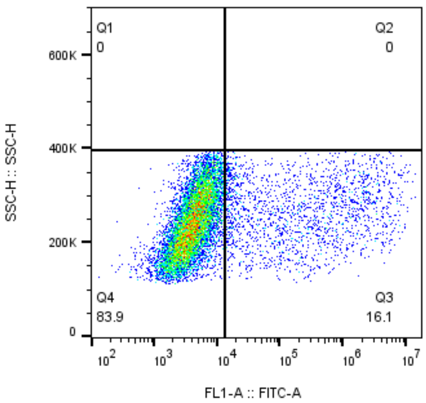


MPI26-80nM]
HEK
11537

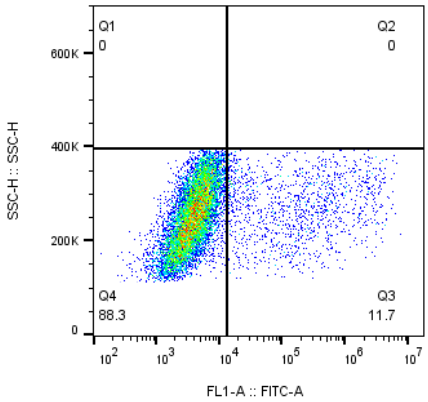


MPI26-16nM
HEK
11129

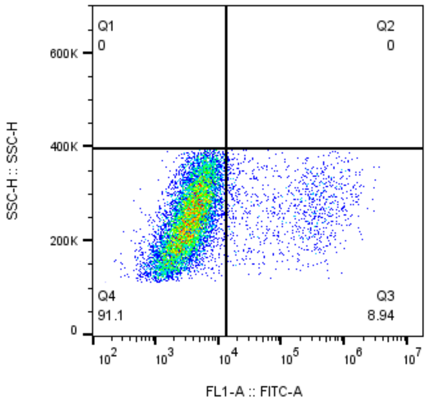


MPI26-3.2nM
HEK
11021

### MPI27

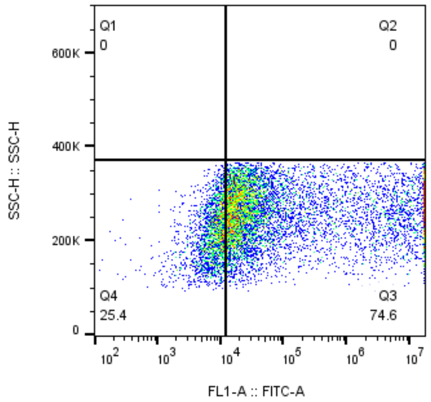


MPI27-10uM
HEK
9718

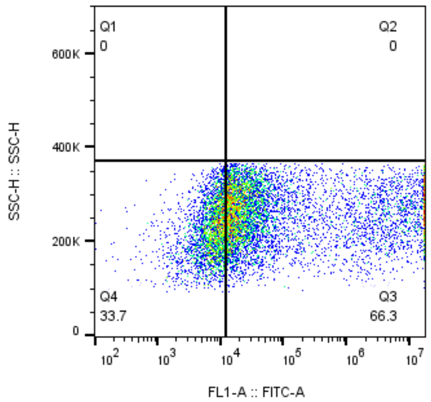


MPI27-2uM
HEK
9676

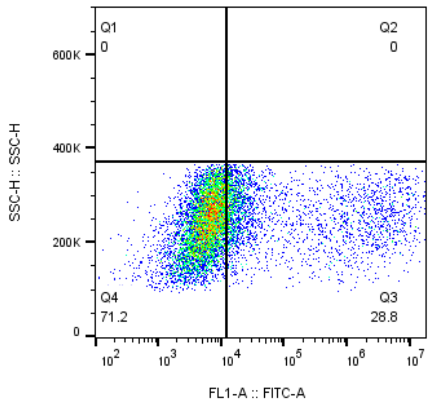


MPI27-400nM
HEK
9714

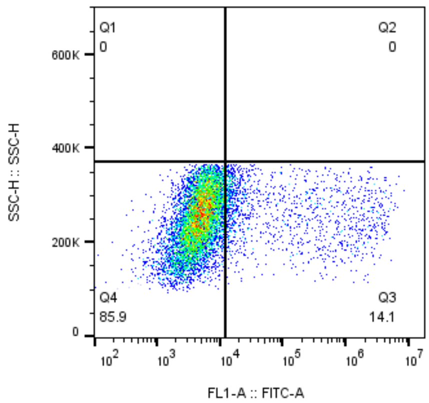


MPI27-80nM
HEK
9987

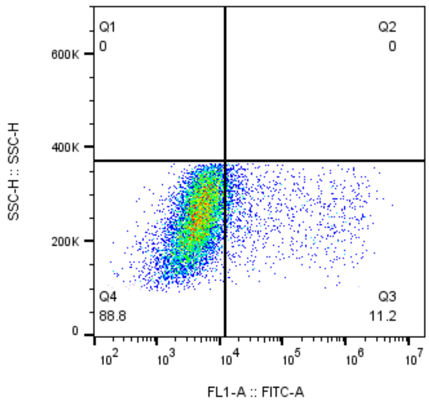


MPI27-16nM
HEK
9631

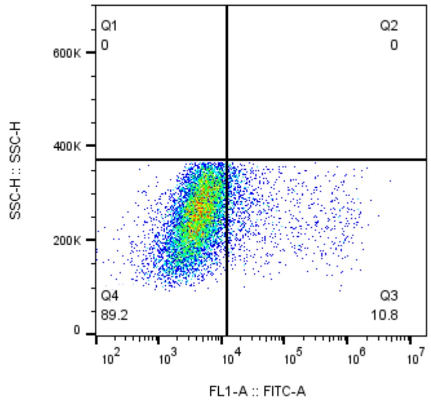


MPI27-3.2nM
HEK
9598

### MPI28

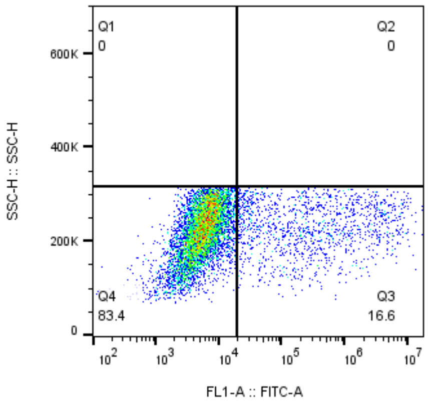


MPI28-10uM
HEK
8268

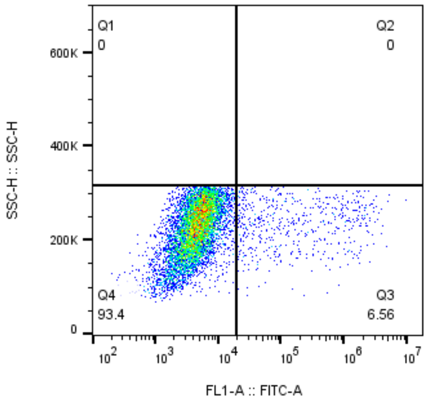


MPI28-2uM
HEK
8242

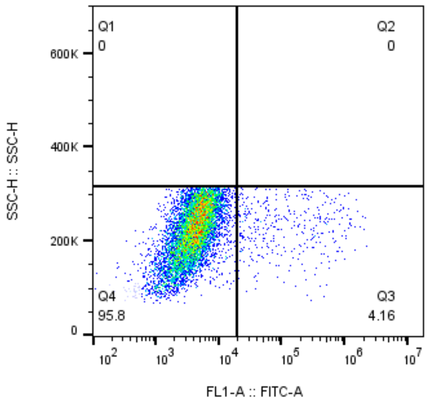


MPI28-400nM
HEK
8453

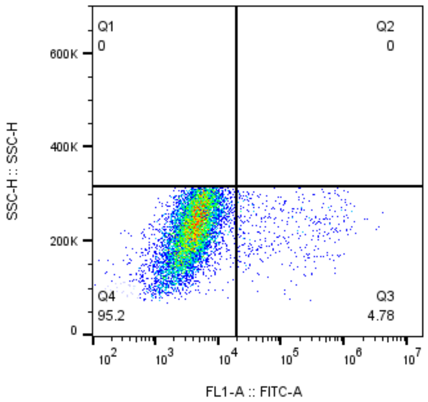


MPI28-80nM
HEK
8396

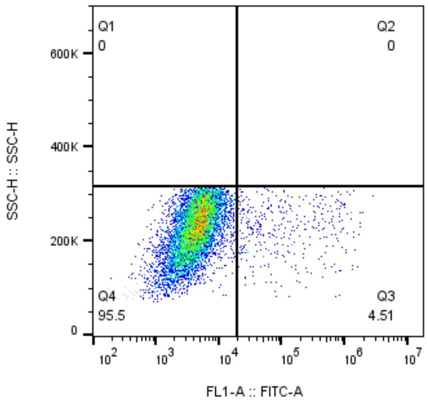


MPI28-16nM
HEK
8027

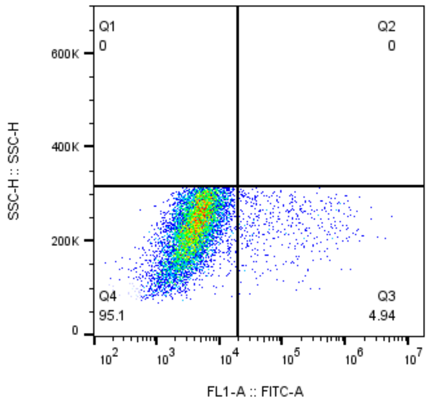


MPI28-3.2nM
HEK
8466
